# From Text-based Genome, Population Variations, and Transcriptome Datafiles to SQLite Database and Web Application: A Bioinformatical Study on Alfalfa

**DOI:** 10.1101/2021.08.25.457609

**Authors:** Shichen Qiao, Chen Shen

**Affiliations:** Shichen Qiao, Undergraduate Student, Department of Electrical and Computer Engineering, Department of Computer Sciences, University of Wisconsin-Madison, Madison, WI 53715; Chen Shen, State Key Laboratory of Agrobiotechnology, College of Biological Sciences, China Agricultural University, Beijing, China

## Abstract

In this study, a web database application with the Flask framework was developed to implement three types of queries and visualize the results over a bioinformatical dataset from Alfalfa (*Medicago sativa*). A backend SQLite database was constructed from genome FASTA, population variations, transcriptome, and annotation files with extensions “.fasta”, “.gff”, “vcf”, “.annotate”, etc. Further, a supplementary command-line-based Java application was also developed for faster access to the database without direct SQL programming. Overall, Python, Java, and HTML were the main programming languages used in this application. Those scripts and the development procedures are valuable for bioinformaticians to build online databases from similar raw datasets of other species.

## 1. Introduction

The field of bioinformatics has been brought to a new era as web server technologies, internet services, and database management tools develop rapidly in recent years. Serval standard data file formats, such as the FASTA Format (.fasta), the General Feature Format (.gff), and the Variant Call Format (.vcf), are used world-widely to store raw genetic sequences, genomic annotations, and population variation data (Gourlé, n.d.). There are many other text-based annotations with different file extensions like “.annotate”, “.pfam”, and “.interpro” as well. For many species with complex chromosomal compositions, these data can be fragmented and enormous as a whole. It is inconvenient and time-consuming to do cross-file searches and analyses on these data without help from some complex online platforms or tools such as BLAST (U.S. National Library of Medicine, n.d.). However, those platforms may not be customized or convenient enough to meet every research group’s needs. Thus, integrating the genome, variation, transcriptome, and annotation data of a species into a relational database, and then utilizing a personalized website application to interact with it could be a better approach for research labs to manage their datasets.

In this article, a case study on Alfalfa (*Medicago sativa*) was conducted with MacOS 11.2 and CentOS 7. The source dataset was adopted from a journal published on *Molecular Plant* (Shen et al., 2020, *The Chromosome-Level Genome Sequence of the Autotetraploid Alfalfa and Resequencing of Core Germplasms Provide Genomic Resources for Alfalfa Research*). SQLite database was chosen because it requires fewer computer resources and it has easy connections to Java and Python programs. R, Microsoft Excel, and different text editors were also used to process data.

The products of our case study include a SQLite database file (.db), a Flask-framework-based web application, and a supplementary, packaged, command-line-based Java application (.jar). Both applications have three query modes across all different-typed files in the original dataset: [A] querying about a gene ID for all related genome, transcriptome, variations, and annotation data; [B] querying about all gene IDs on file whose annotations contain some given keywords; [C] querying about all gene variation information on a certain portion of a chromosome. The web application visualizes the data through bar charts, table coloring, and categorization, and it is capable to link annotations to their source databases’ websites automatically.

## 2. Data Selection

The data involved in this study were adopted from genome sequencing results and transcriptome annotations on Alfalfa (Shen et al., 2020), stored as shown in **Figure 1**. After decompression, the FASTA, GFF, VCF, and annotation files were about 600 GB in total.

**Figure 1.**
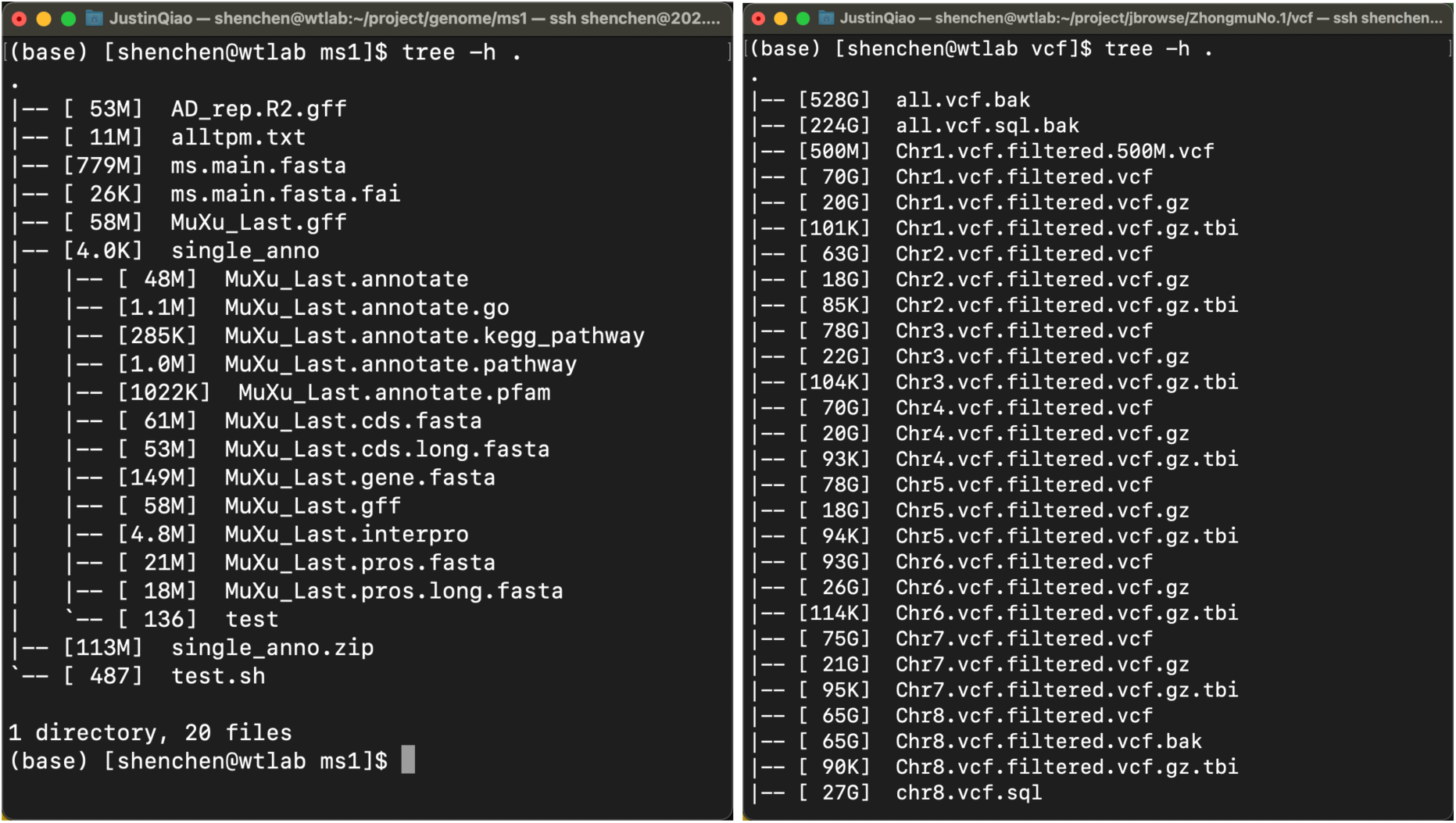
Complete dataset on our CentOS server.

After data selection, the files shown in **Figure 2** were downloaded to the Mac computer which we used primarily during the development stage. Although only 500 MB of the VCF records were kept locally, once the proper scripts were developed base on this small sample, the server would execute the same scripts and insert the complete VCF records into the database file. Further, ms.main.fasta was renamed as Ms_genome.fasta for clarity, and it was composed of the DNA sequences of 8 chromosomes and 649 contigs. Moreover, transcriptome data were stored in alltpm.txt, and the other files were annotations. A sketch map of these file formats is shown in **Figure 3**.

**Figure 2.**
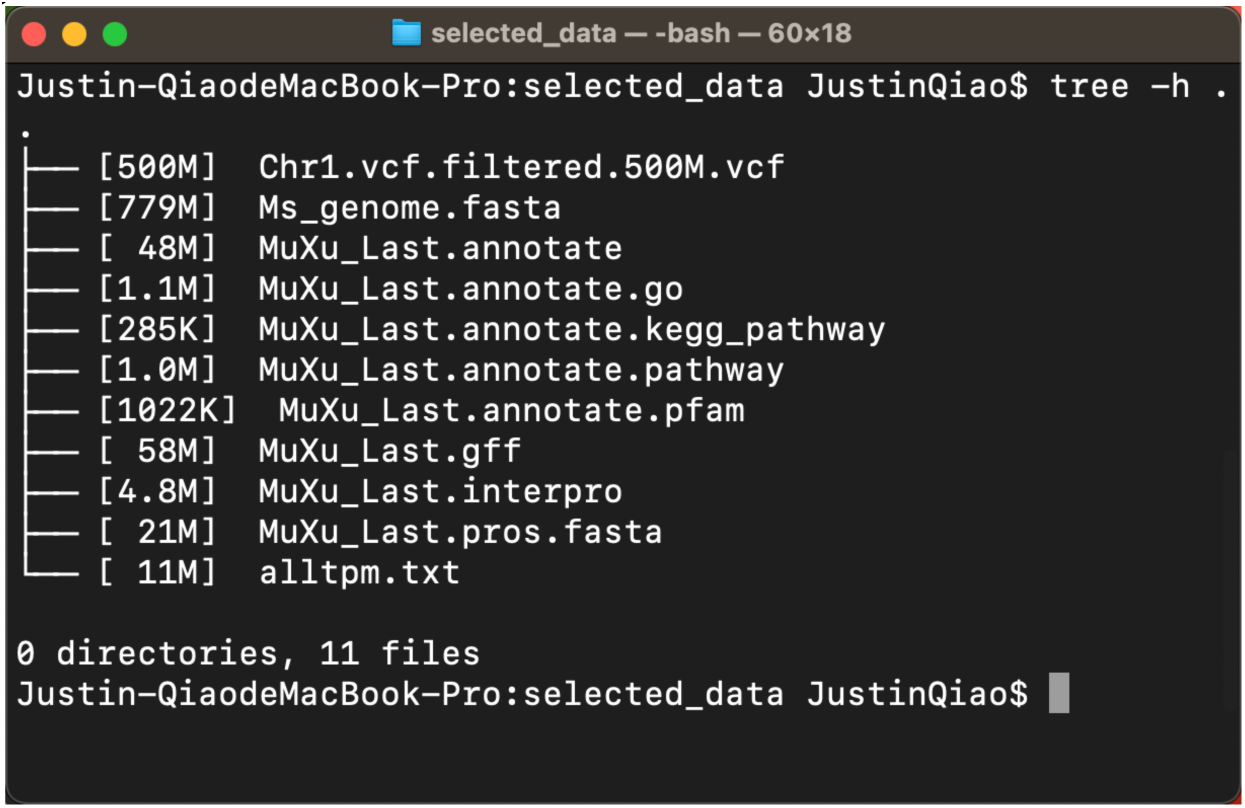
Selected dataset.

**Figure 3.**
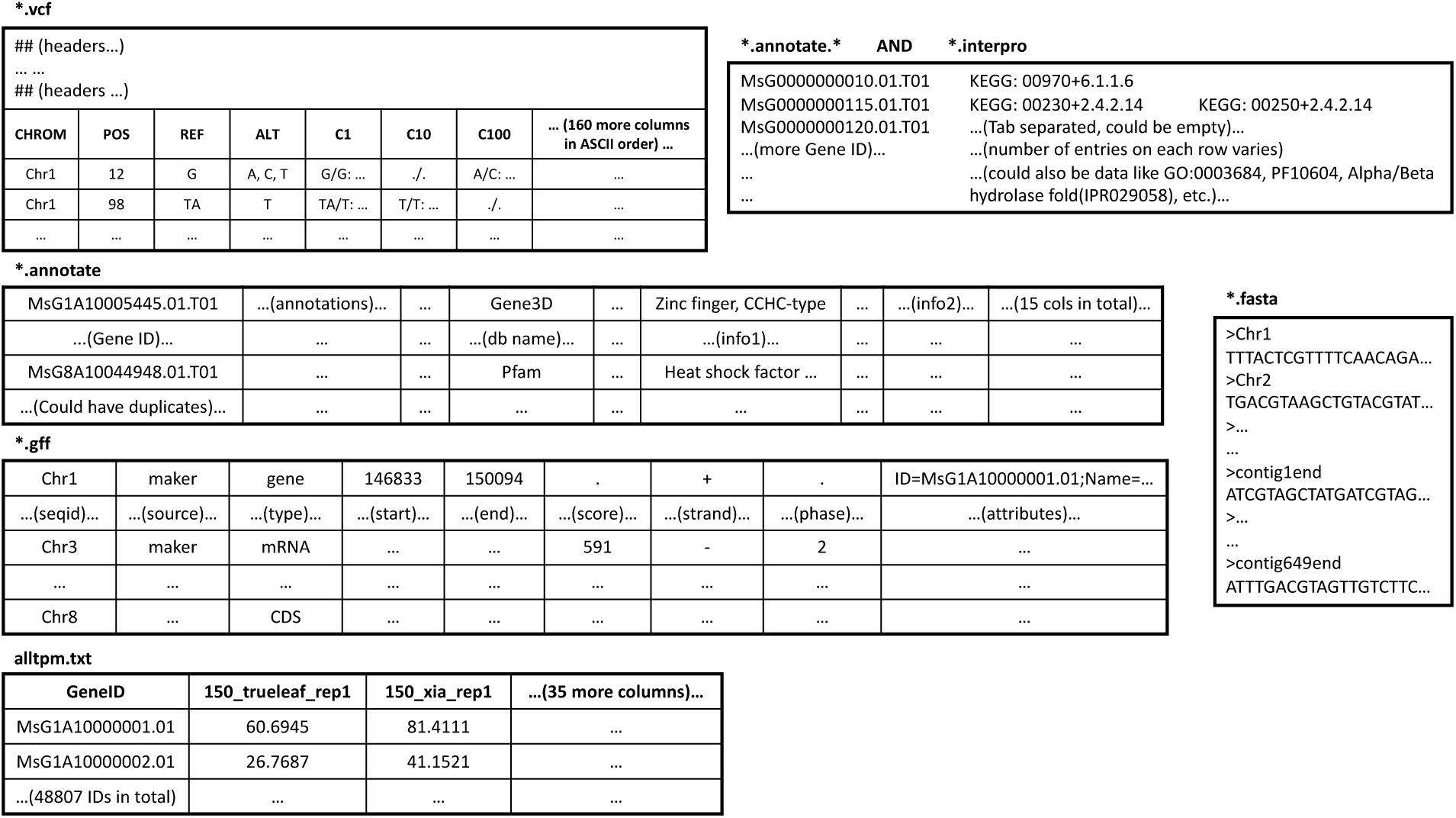
Selected datafiles’ formats.

## 3. Software Preparation

To minimize development costs, a MacBook Pro laptop with MacOS 11.2 was chosen to be the main development environment. A CentOS 7 Linux server was used to execute some Python, Java, and SQL scripts on the large raw datafiles, and it was also used to broadcast our final web application to the Local Area Network (LAN). The essential software applications used during our development process were recorded in this section.

### 3.1 SQL Environment and GUI

SQLite3 is a free, server-less, self-contained, cross-platform SQL database engine (SQLite Org, n.d.). The latest version of SQLite3 is pre-installed on most Mac machines. To utilize this database engine more intuitively, a visual management tool with graphical user interfaces (GUIs), DB Browser for SQLite, was used along with the procedure (**Figure 4**). With this application, developers could manage the structure of their database, browse and filter data, and execute SQL scripts more conveniently (Sqlitebrowser Org, 2021).

**Figure 4.**
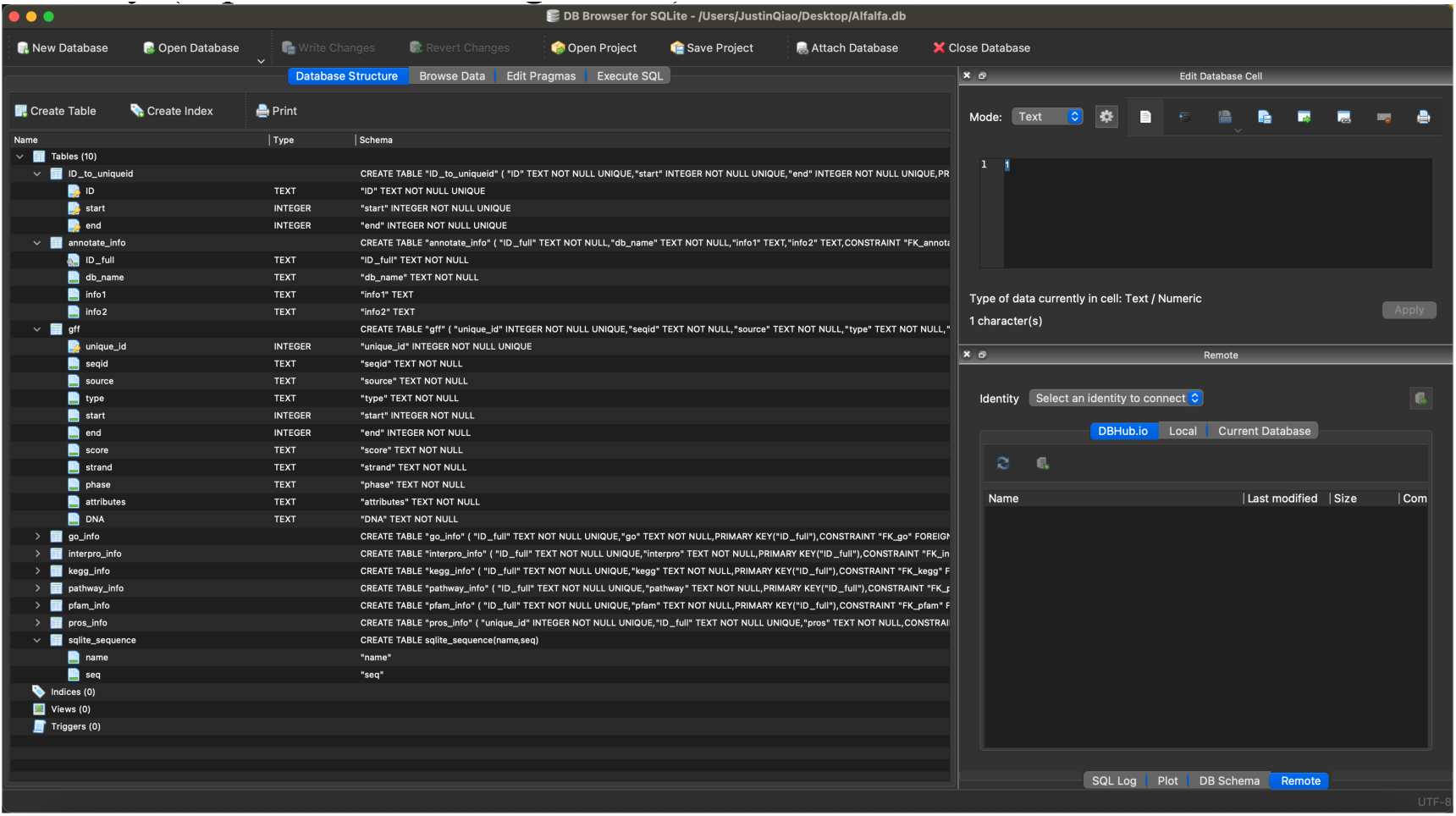
DB Browser for SQLite GUI.

### 3.2 Visual Studio Code and Programming Languages

According to Guo (2014), Python and Java are the top two popular programming languages in US undergraduate education. We decided to use them collaboratively in our case study to take advantage of both languages’ strong points. Additionally, HTML, CSS, and JavaScript were used for the web application. As a result, we chose Visual Studio Code as our main code editor since it is capable to manage all of these languages cooperatively (Microsoft, 2016). Note that PyCharm by JetBrains S.R.O. and Eclipse by the Eclipse Foundation are more centralized Integrated Development Environments (IDEs) that were also helpful.

Specifically, Python 3.8 by Python Org., JavaSE-11 and JavaSE-1.8 by Oracle Corporation, HTML5 and CSS3 by the World Wide Web Consortium were utilized in our study. Moreover, to connect our Java applications to the SQLite database, a Java Database Connectivity (JDBC) interface was required. Maven Repository provides free SQLite JDBC on their official website, and the latest version, 3.36.0.1 was used in our development process (MvnRepository, n.d.).

### 3.3 The Flask Framework

The Flask Framework is a web application framework developed by Armin Ronacher. It manages Python web applications. Based on this framework, our HTML page and the Python application could communicate with each other, pass data inputs and query results back and forth, and broadcast the final online database website through an IP address conveniently.

### 3.4 Microsoft Excel and/or Numbers

As mentioned in the Introduction section, the annotations are stored in text-based files with different extensions. However, by observation, the annotation files are essentially Tab-Separated Values (TSV) data. Each row in an annotation file contains information about one gene ID, and that information could be divided into different “columns”.

With these characteristics, spreadsheets applications such as Microsoft Excel and Numbers are perfect choices for researchers to view and manipulate the annotations to a format that is easy to be inserted into a SQL database graphically.

In our case study, Microsoft Excel was used to convert TSV files into Comma-Separated Values (CSV) files, which enabled us to deal with annotations stored in different files with the same Python algorithm. In addition, the sorting and concatenation functions in Excel also saved a lot of time for us to generate SQL scripts.

## 4. Methods: Backend SQLite Database Development

In this section, the development procedure of our SQLite backend database was explained. There were multiple input raw data files, Python and Java scripts, intermediate data files, and SQL scripts involved during the process, but there was only one database output file (Alfalfa.db). The integration of these fragmented data files into one portable file was one of the main purposes of developing this SQL database.

### 4.1 Database Structure and Table Creations

The database structure was designed to make the three types of queries, as mentioned in the Introduction section, faster and simpler. By observing the datafiles, we found that the gene ID was the main connection among the datafiles shown in **Figure 2**. Specifically, a gene ID number could contain 13, 16, or 20 characters. For our dataset, the first 13 characters of any gene ID correspond to a specific gene of Alfalfa, so we named this part the “Primary ID”. Further, if there existed such characters, the 14th to 16th characters could only be “.01”, the 17th to 18th characters could only be “.T”, which stood for Transcript, and then followed by two digits. Thus, we named character #14 to character #20 the “extension”. Since the majority of gene IDs in the raw dataset had 20 characters, we name such gene ID as “ID_full”.

Next, we classified the datafiles into five categories. Firstly, MuXu_Last.gff contains 580,707 rows of 9-column GFF entries. Using data in the seqname, start, and end columns, we could chop out the DNA sequence for each GFF record from Ms_genome.fasta and add it to a “DNA” column following the existing “attribute” column (The Ensembl Project, n.d.). We then give each row a uninque_id as Primary Key (PK), and we could obtain Table gff as shown in Section 10.1 on Line 58 through 71. We also added a SQL Table called “ID_to_uniqueid” to map each primary ID to its contiguous block of unique_ids, and a Table called sqlite_sequence was created by our environment automatically at this point. Secondly, the go, kegg_pathway, pathway, pfam, interpro, and pros.fasta datafiles were classified as “info” datafiles. Specifically, each entry in these files started with a 20-character gene ID and then followed by a various number of data cells. Thus, we used ID_full as PKs and created Table go_info, interpro_info, kegg_info, pathway_info, pfam_info, and pros_info as shown in Section 10.1 on Line 73 through 114. In addition, transcriptome data stored in alltpm.txt were planned to be transferred into a 48,807 row by 38 column SQL Table directly, as shown on Line 9 through 48. Moreover, we had MuXu_Last.annotate storing gene annotations from other online databases of *Medicago Truncatula*. This file was opened in Microsoft Excel as Tab Separated Values (TSV), as shown in **Figure 5**, and we took out column A (ID_full), D (db_name), F (info1), and M (info2) to create Table annotate_info for keywords queries, as shown in Section 10.1 on Line 50 through 56. Finally, we observed the 500 MB VCF sample and found that it was structurally a table with 167 columns. Thus, we placed a Primary Key column called “vcf_ID” in front of the original columns and created Table vcf as shown on Line 118 through 288.

**Figure 5.**
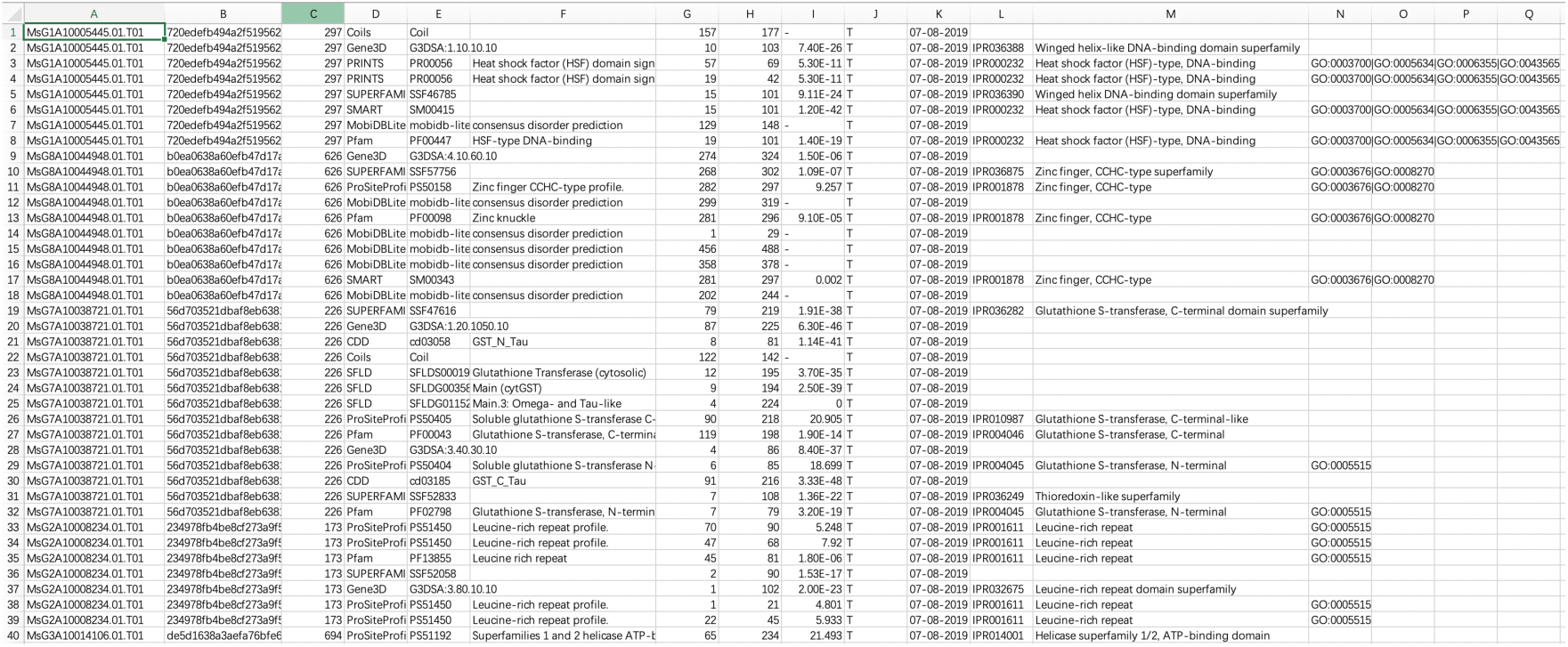
Part of MuXu_Last.annotate shown in Excel.

### 4.2 Pre-processing Annotation and FASTA Datafiles with Python

After table creations, the raw data needed to be pre-processed before insertions into our database through SQL scripts. In this section, methods of pre-processing FASTA and annotation files using Python would be elaborated.

We started with the go, kegg_pathway, pathway, pfam, and interpro files because they could be managed with very similar scripts. Observing the interpro annotation file, we noticed that no matter there were how many interpro IDs for each gene ID, they were all concatenated together using “|” as the separator. This could easily solve the problem that, in the other annotation files, the number of annotations attached to an ID_full was not unified. Thus, for the other four files, the following Python script was used to concatenate the annotations for each ID_full using semicolons as the separator:

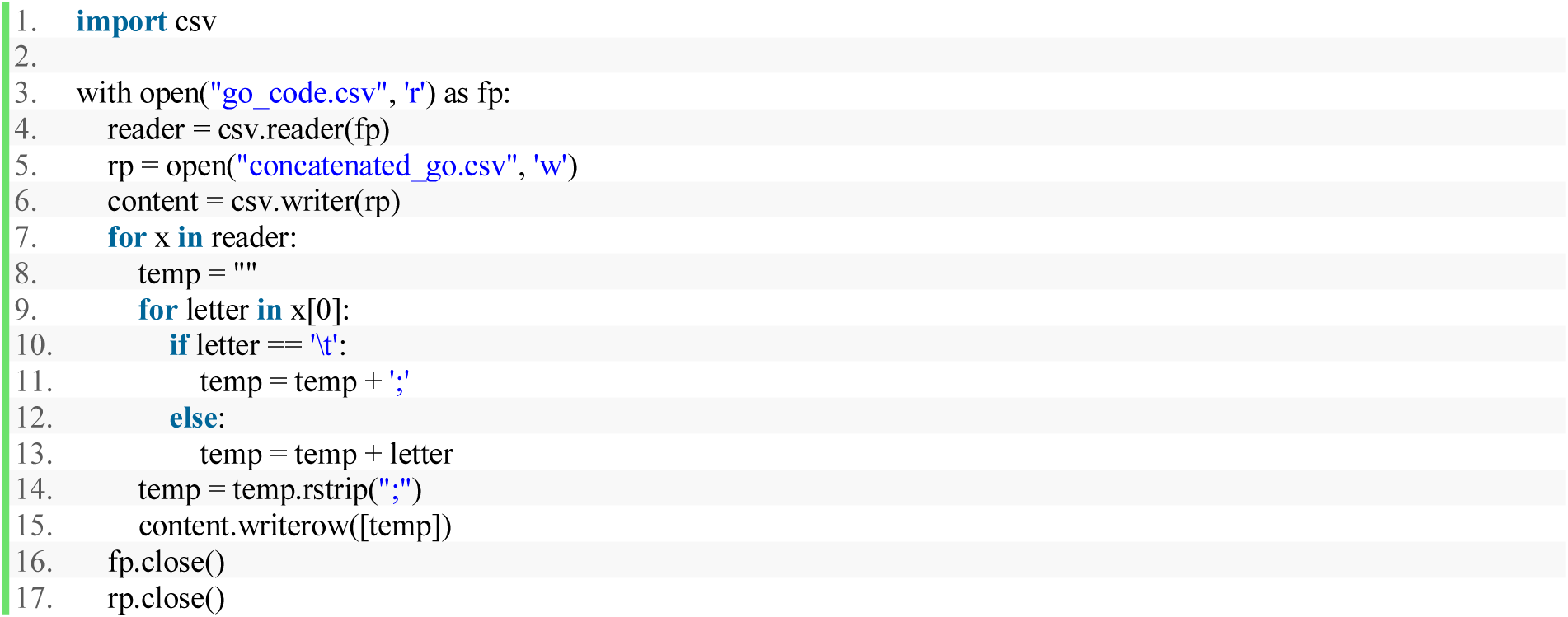

In addition, MuXu_Last.pros.fasta contained data with similar properties as well. Each FASTA entries started with a “>” followed by an ID_full, and then a protein sequence was on its following line. Thus, this file could be converted into a 2-column CSV table of key-value pairs, similar to the five files pre-processed above, by running the following Python script:

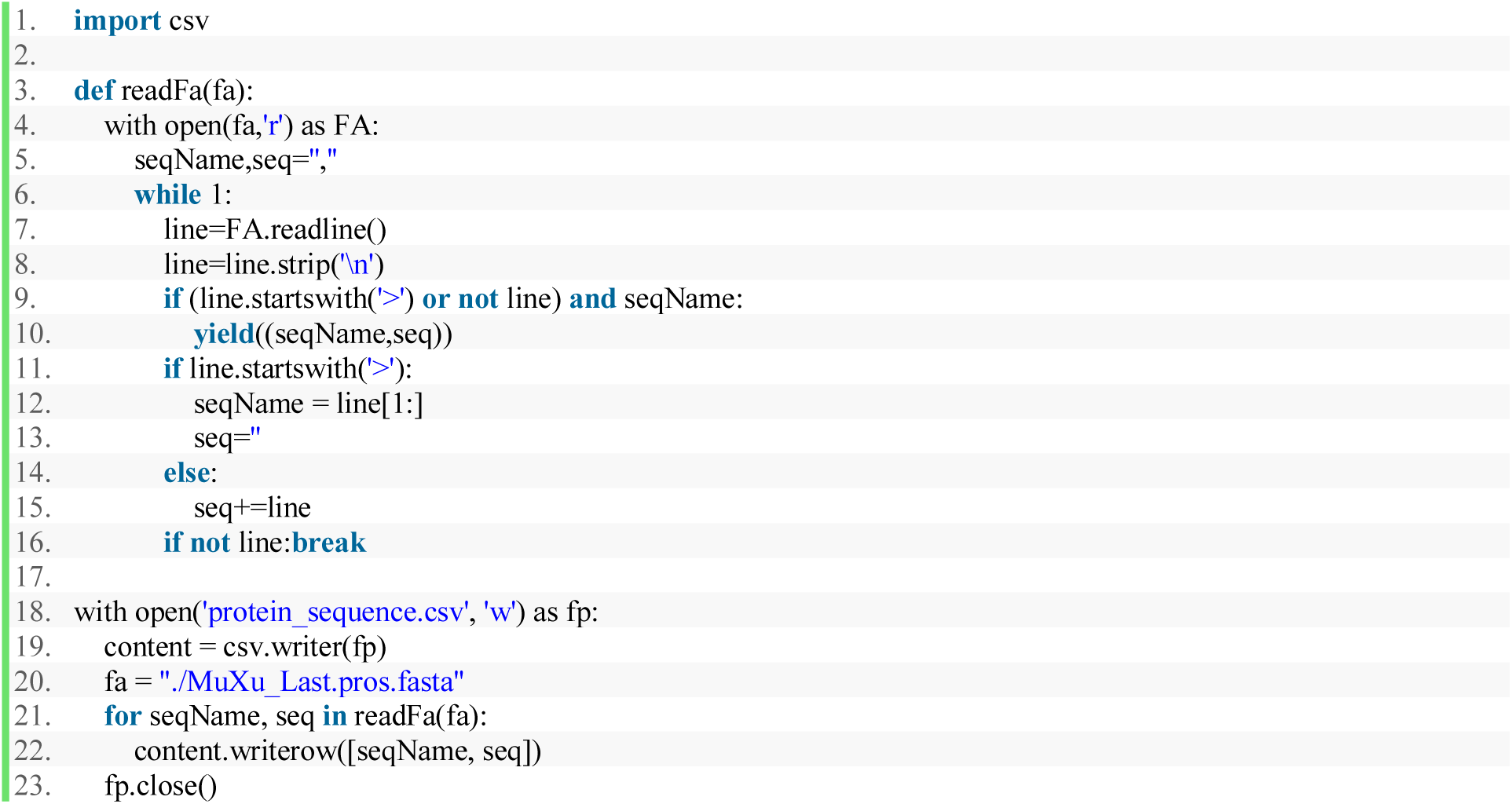

### 4.3 INSERT Pre-processed Data to SQLite Tables

After pre-processing, all files in our original dataset shown in **Figure 2** except Ms_genome.fasta and the VCF file were ready to be converted into SQL INSERT statements. Those treated CSVs and TSVs could be opened in Microsoft Excel as spreadsheets formats and used to generate SQL scripts.

For instance, when we were processing the GFF datafile, we changed the “.gff” extension to “.txt”, and then opened it with Excel using Tab as the delimiter. Next, the data were sorted by the ASCII order of the attribute section, which made sure that GFF entries belong to the same gene ID were clustered. Finally, the “unique_id” column was inserted into the spreadsheet, giving each row of GFF entry a unique tag as we planned (**Figure 6**).

**Figure 6.**
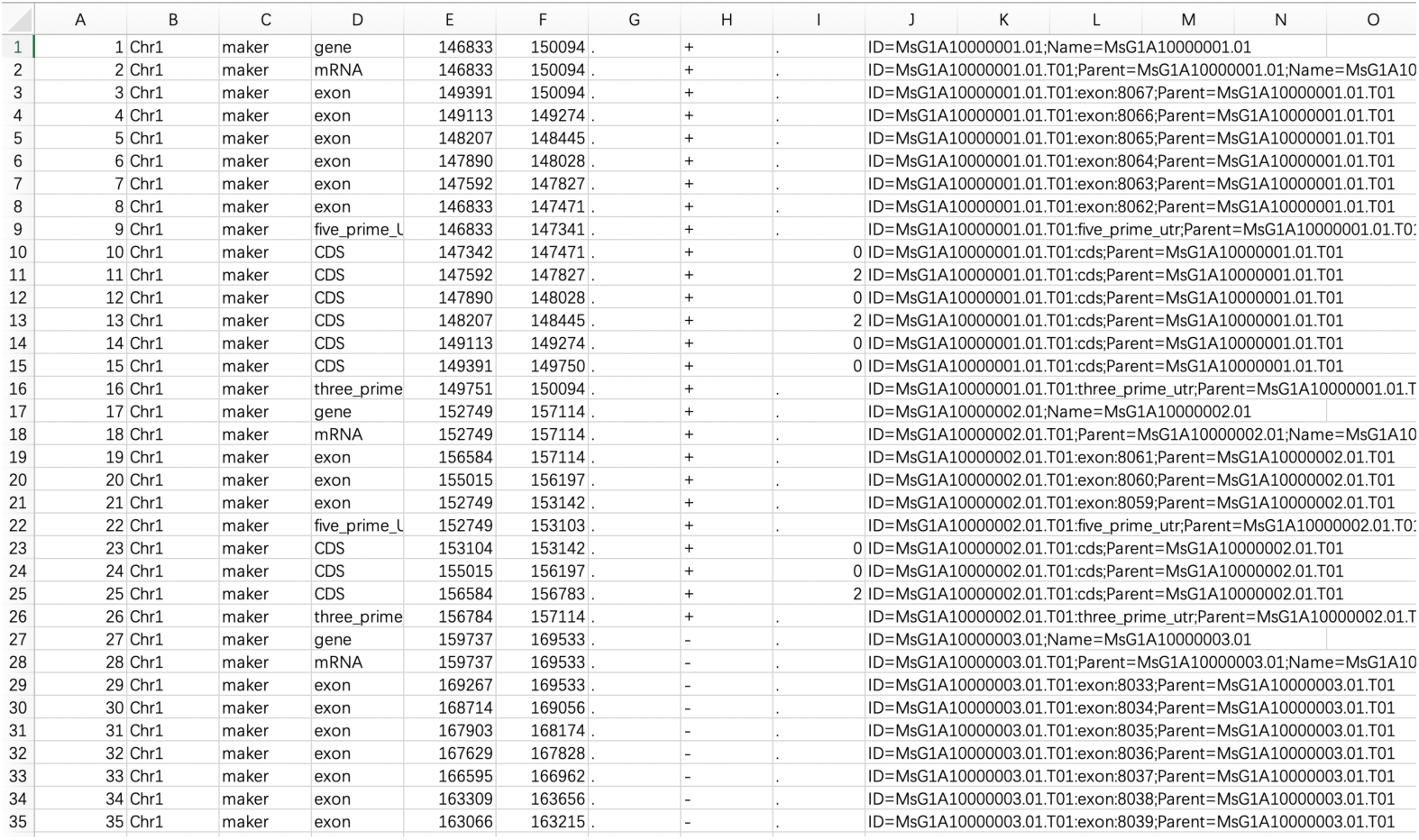
Part of MuXu_Last.gff opened in Excel with unique_id added.

At this point, all data in the GFF file had been prepared in a table format for insertion into the database. A build-in function of Excel called “CONCATENATE” was used at the K1 data cell of the spreadsheet shown in **Figure 6**:

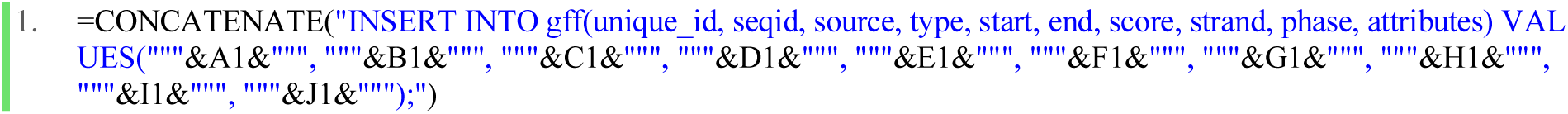

An example of the resulting SQL command is:

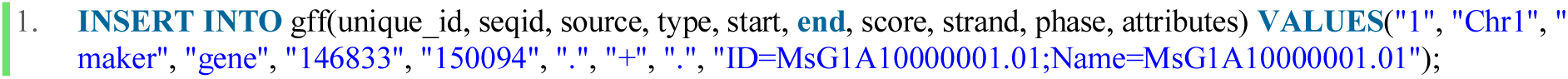

Later, the whole K column was selected, and a shortcut key, Control plus Enter, was used to implement the equation to each row and generate the complete insertion SQL script automatically. Copying the entire column K to a text file with the extension “.sql” and executing it in DB Browser for SQLite gave us Table gff as shown in **Figure 7**.

**Figure 7.**
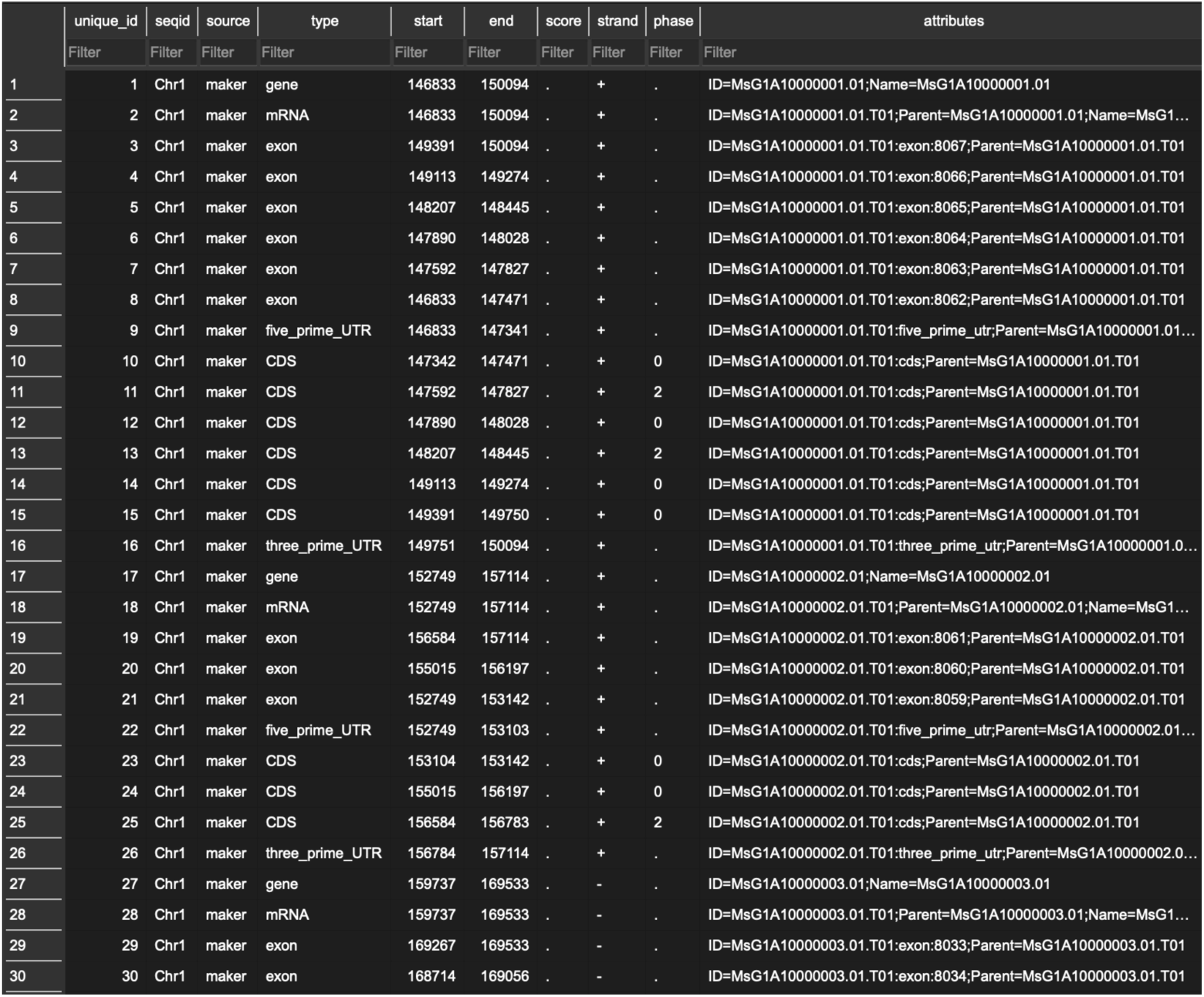
Part of Table gff in Alfalfa.db shown in DB Browser for SQLite.

For the pre-processed CSV files, they could be opened in Excel with comma as the delimiter, and then similar CONCATENATE functions could be applied. Notice that not all characters could be handled directly using this procedure. For example, while creating Table annotate_info, we found that 15 Excel-generated SQL commands had syntax errors because the character “-” was not supported in INSERT instructions. One of these invalid commands was:

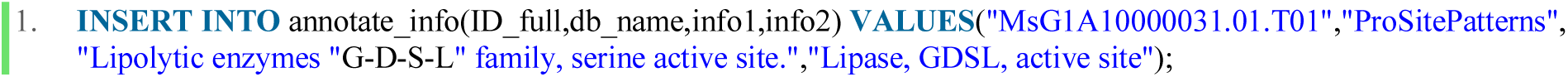

Since there were only 15 exceptions, we added these entries manually to Table annotate_info via DB Browser for SQLite. If there were much more exceptions, replacing these characters might be necessary.

Moreover, when the source datafile contained too many columns, the convenience of using Excel would be largely reduced. In such cases, using Python scripts to convert the text-based file directly to SQL INSERT statements could be a better option. For example, alltmp.txt was processed by the following Python program:

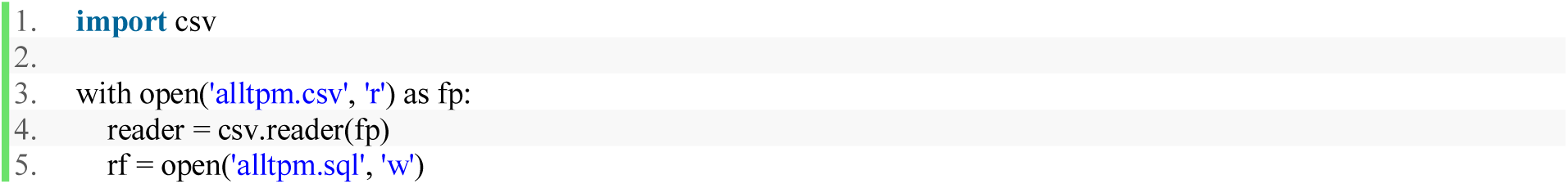
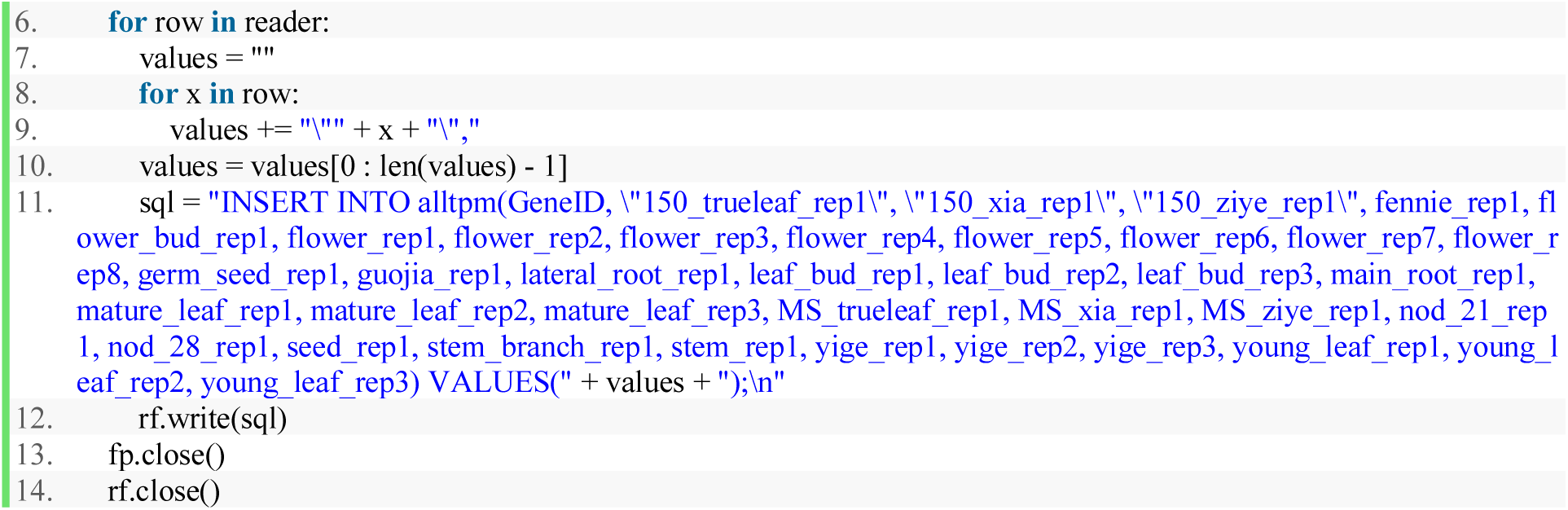

### 4.4 Improving Table gff

As mentioned in Section 4.1, a linkage table called “ID_to_uniqueid” was created to optimize gene ID queries from Table gff. The first 13 characters after “ID=” in the attribute column in Table gff would be extracted out and stored in the ID column in this helper Table. The extraction was done by the Convert Text to Columns Wizard in Excel, using “=” and “.” as delimiters. Then, the unique_id column and the extracted primary ID column were saved as a separate CSV file named “uniqueid_ID_primary.csv” (**Figure 8**).

**Figure 8.**
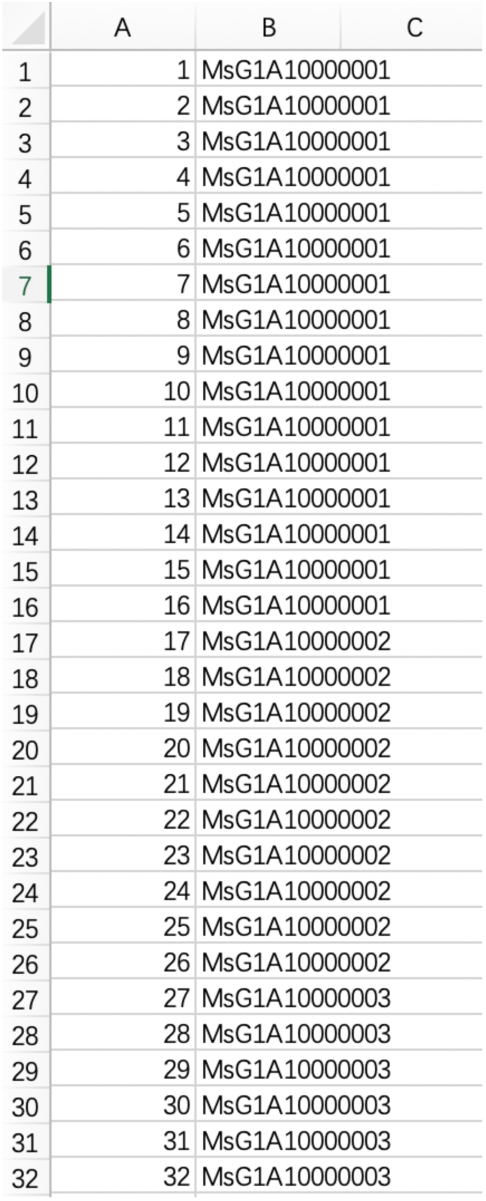
Part of uniqueid_ID_primary.csv shown in Excel.

Next, the following Python script was executed on this CSV file to find out the starting and ending unique_id matched to different primary IDs:

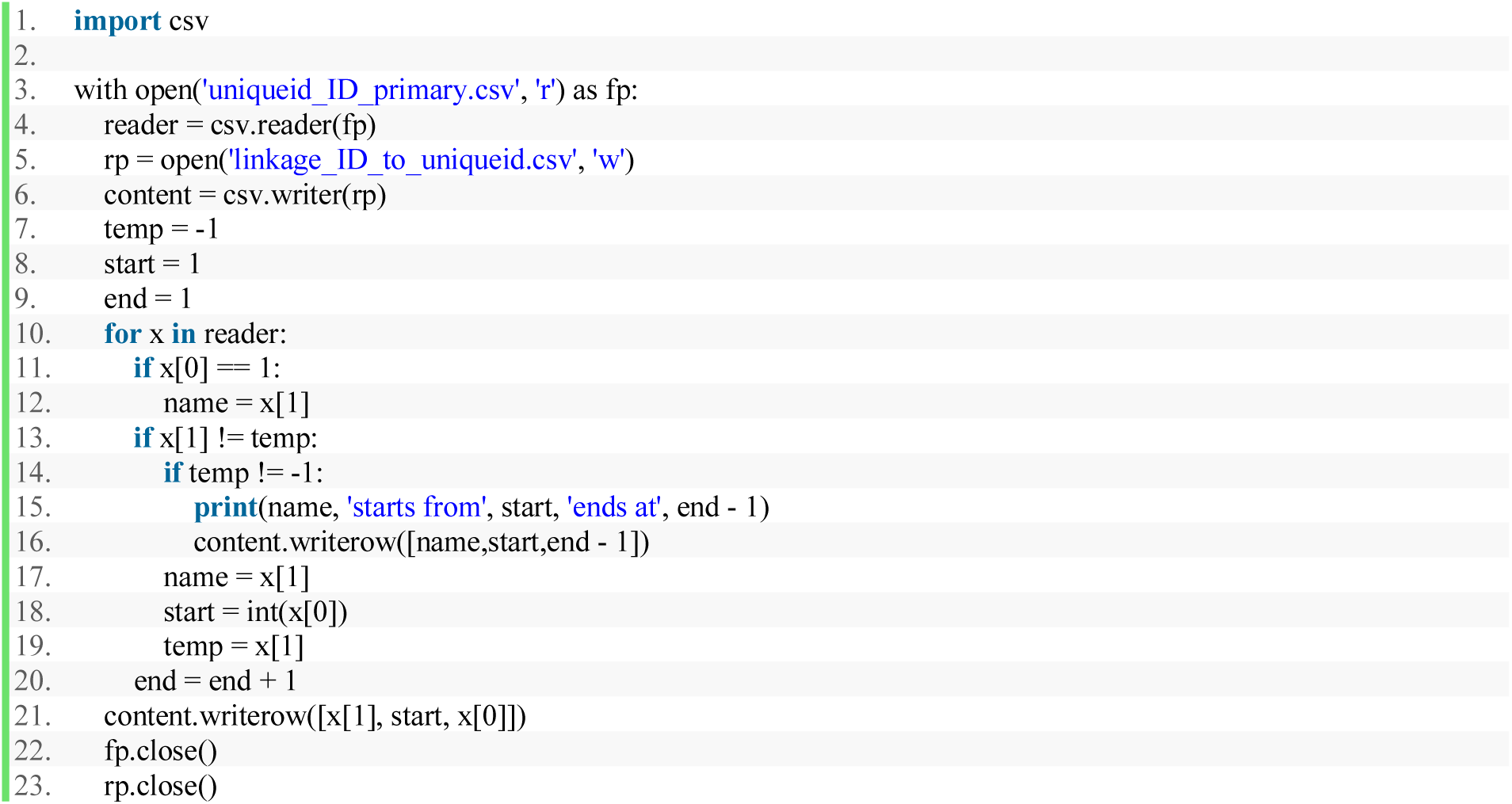

The for-statement traverses through each row of the CSV file, and if the name (primary ID) changes between two consecutive rows, the starting and ending unique_ids would be recorded in the output CSV file.

Using a similar formula, these data were inserted into Table ID_to_uniqueid by executing an Excel-generated SQL script using the following function:

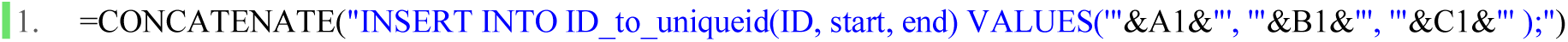

Notice that when the data being inserted does not contain double quotation marks or apostrophes, using ‘“&A1&”’ and “““&A1&””” in the function above would generate the same SQL commands.

Further, for each GFF entry regardless of its type, the substring obtained from the corresponding FASTA entry stored in Ms_genome.fasta between (closed interval) the indexes specified in the “start” and “end” columns in Table gff should be the detailed DNA sequence matched to that GFF row. Thus, Ms_genome.fasta was chopped 580,707 times to obtain the SQL scripts to fill the “DNA” column of Table gff.

However, keeping the entire FASTA file open for these many operations was not efficient. Thus, the following Python script was used to divide Ms_genome.fasta into 657 text files. Each file was named as a seqid followed by “.txt”, and the content was the complete DNA sequence attached to that seqid (the readFa function was defined in Section 4.2).

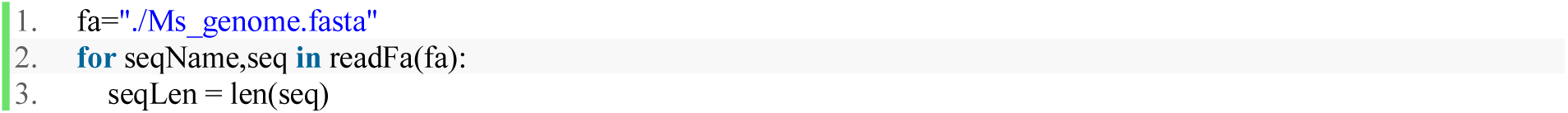
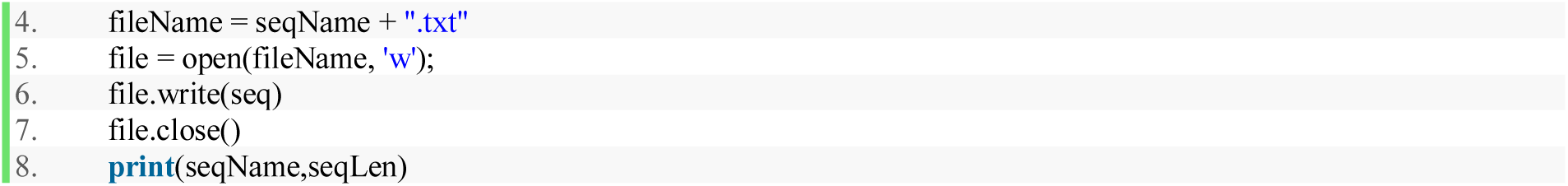

Next, the “unique_id”, “seqid”, “start”, and “end” columns in Table gff was exported as a CSV file called “start_end_raw.csv” from Table gff, and the following Python script was used to truncate the sequences as described above and generated 580,707 SQL UPDATE commands:

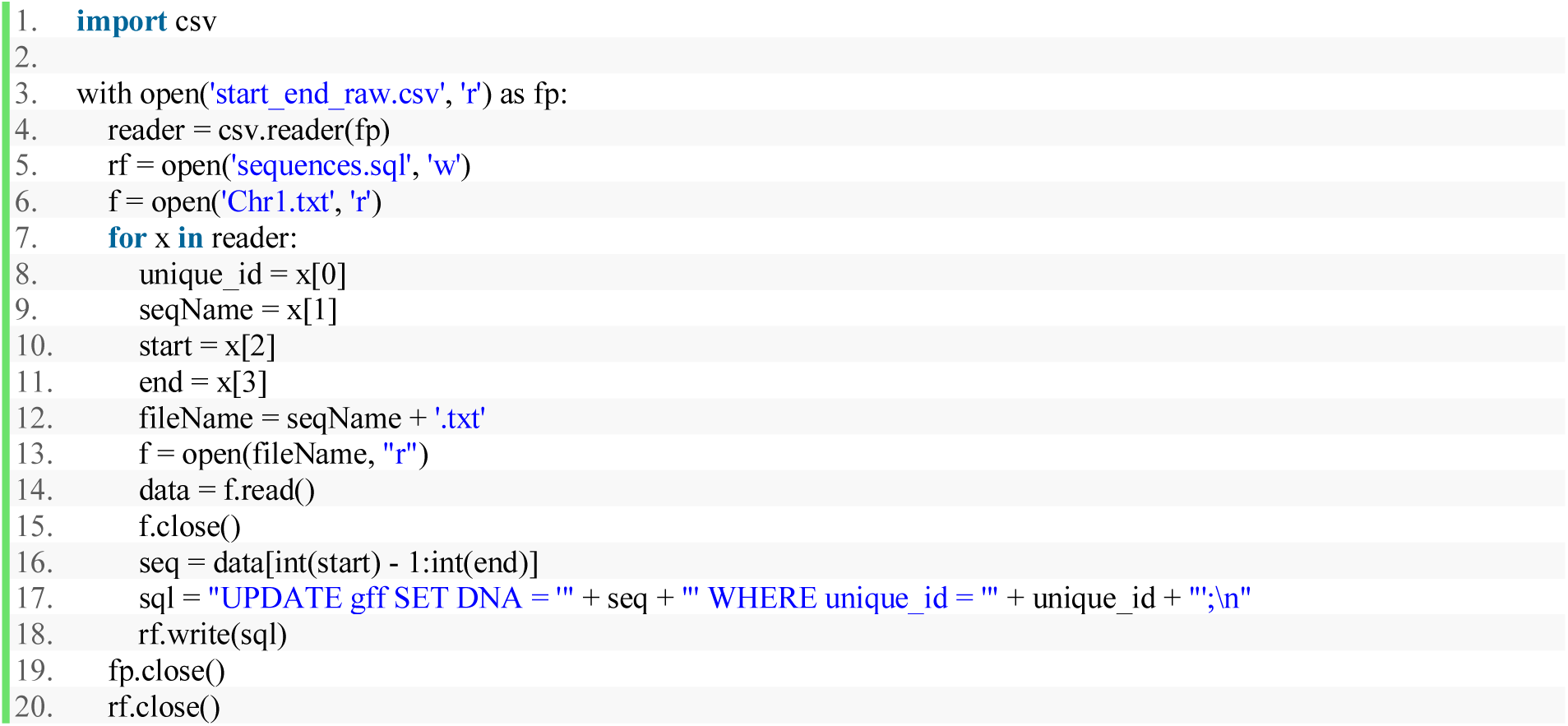

After executing sequences.sql, Table gff was finalized (**Figure 9**).

**Figure 9.**
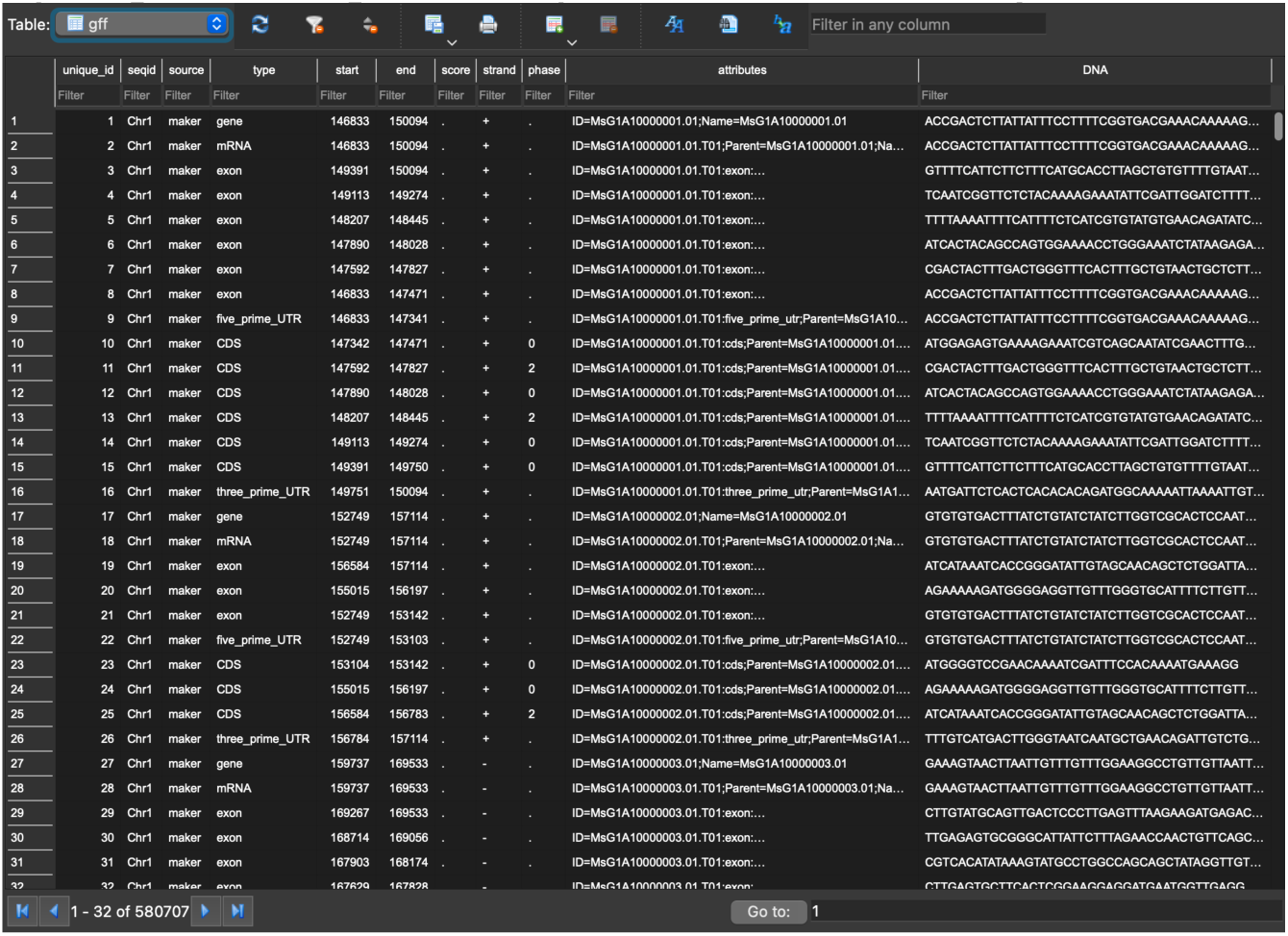
Finalized Table gff with DNA sequences.

### 4.5 Improving Table annotate_info

So far, all data except the VCF file had been added into the SQLite database. However, 81,266 duplicated rows were found in the Table annotate_info. This was caused by the replications of annotations on the same gene in the raw “.annotate” file. To eliminate the redundant copies, only the row with the smallest row number was kept for each cluster of rows with the same contents after running this SQL command:

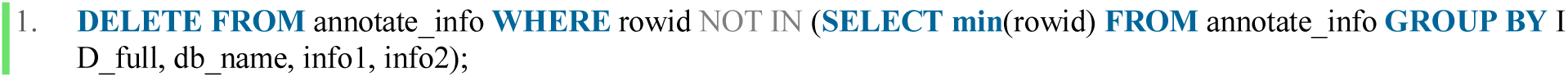

Further, to make frontend development easier, all empty data entries (shown as “” in INSERT statements) was replaced by the NULL character by the following SQL commands:

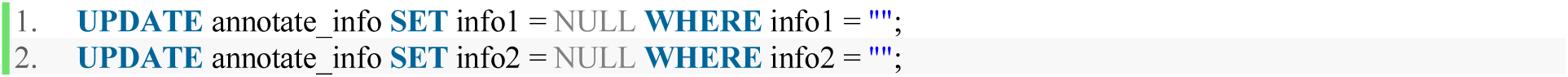

### 4.6 Building Table vcf

The last type of data was the gene variation records stored in the massive VCF files. In this section, the approaches we had taken to transfer VCF data into our SQLite database were discussed.

The first step was analyzing how VCF data were stored. By standards, there is a single row in the VCF file that begins with only one “#”, and all column names are listed on that row. The rows above are file headers that begin with “##” on each row, and each row below describes gene variations at a specific position. That row with column names was taken out, put into a file editor, and stored as a TSV file called “column_names.txt” (**Figure 10**).

**Figure 10.**
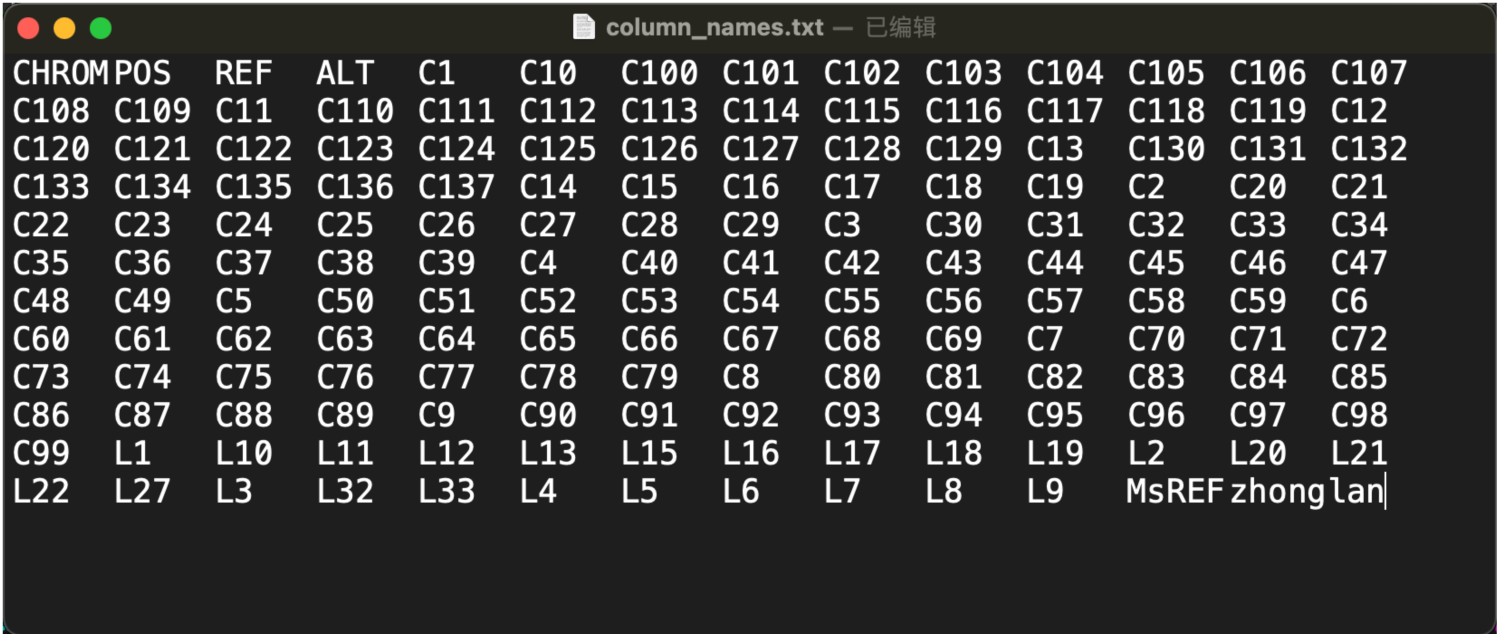
column_names.txt.

Next, we decided to give each variation record a unique number, because a Primary Key was needed to improve searching efficiency. Thus, a similar variable like we used for Table gff was utilized again. To be more specific, a column of consecutive natural numbers, called “vcf_ID”, was added to each row of Table vcf. The beginning and ending vcf_ID of each chromosome would be recorded by the frontend applications to distinguish different chromosomes.

With all column names and properties decided, the following Python script was executed on this text file to generate the CREATE statement in SQL:

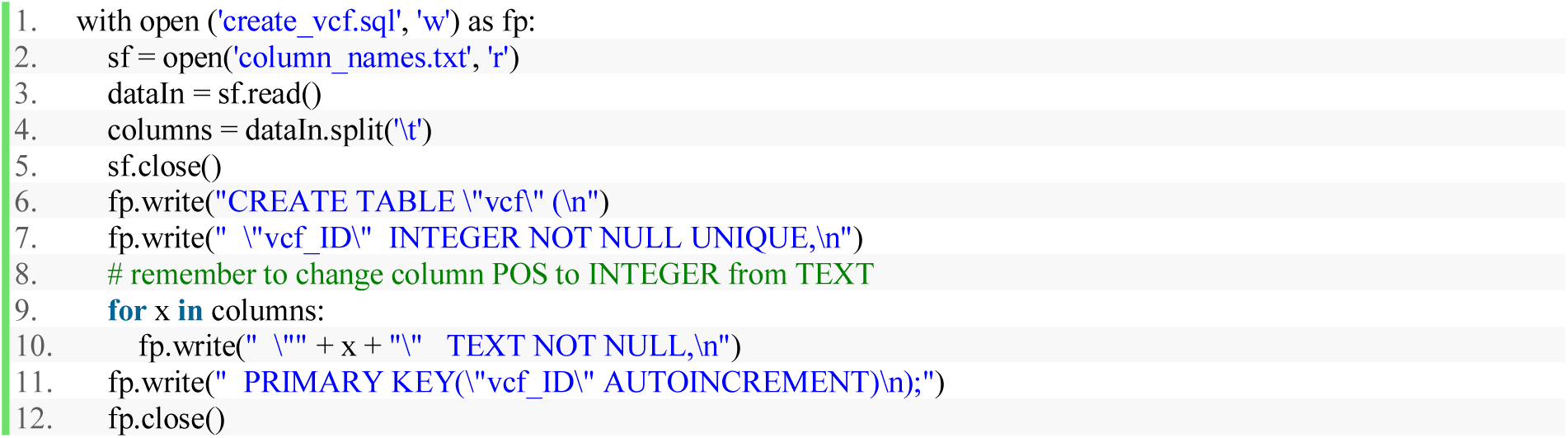

The scripts above unified all columns stored in column_names.txt to be TEXT data for simplicity, but it was necessary to make the “POS” column contain INTEGERs. The next step was filtering out the gene variation data in the VCF file (as shown in **Figure 11**), and converting it into a more human-readable way. By our design requirements, we were only interested in the actual nucleotides at each position (POS). In the GFF file, “./.” means the variation data at the corresponding position in a specific breed of Alfalfa is not applicable. If for a certain breed, gene variation data is available, that data would be in a format like “0/1:1,2:3:28:67,0,28”. The number right before and right after the “/” character is most important to our database. If a number was 0, it means that the gene at that specific position was the same as the “REF” value on the same row; if a number was 1, it means the gene at that position is a variant same as the first variation listed in column “ALT”; if the number was 2, it matches to the second variation in column “ALT”; etc.

**Figure 11.**
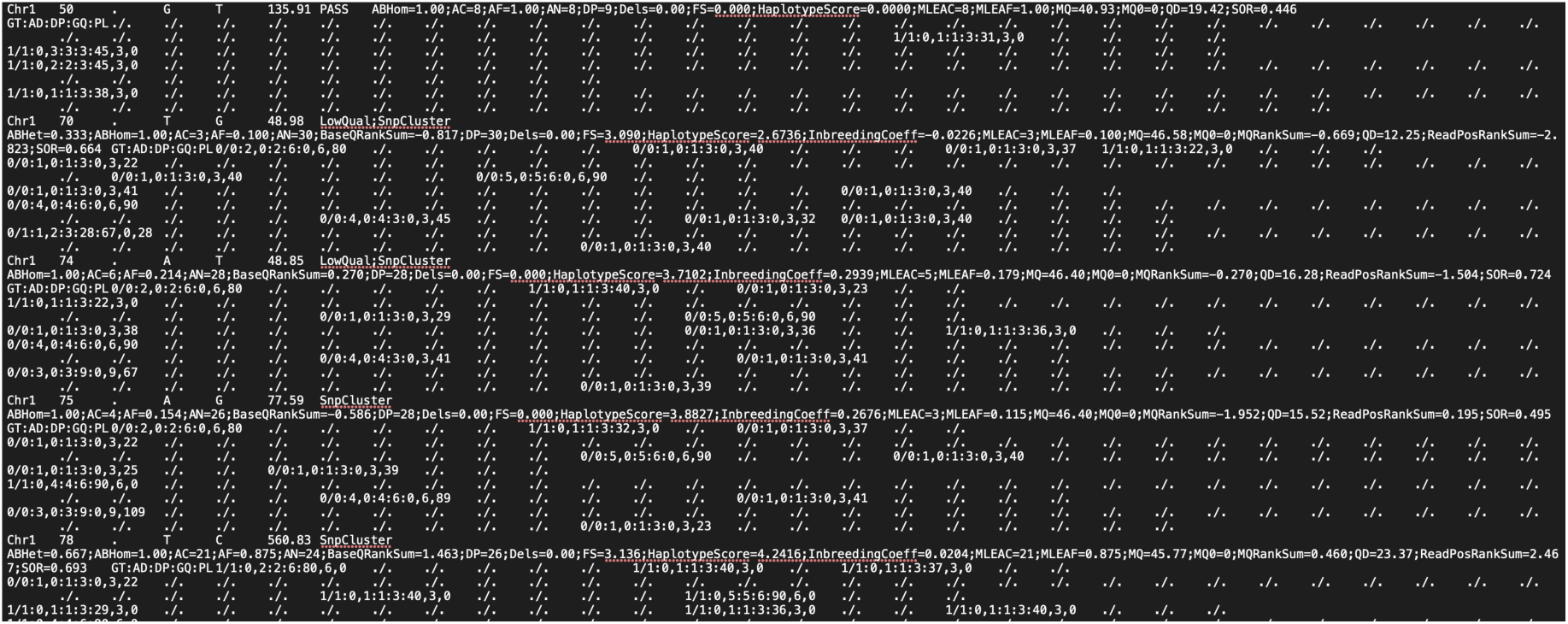
Part of raw VCF data.

Manually looking back and forth between the numbers and the “REF” column was not efficient. Thus, we replaced all numbers with the actual nucleotides they represent, and generated SQL INSERT statements for the VCF data by running the Java application shown in Section 10.2. Finally, executing the generated SQL script gave us Table vcf in our database file (**Figure 12**).

**Figure 12.**
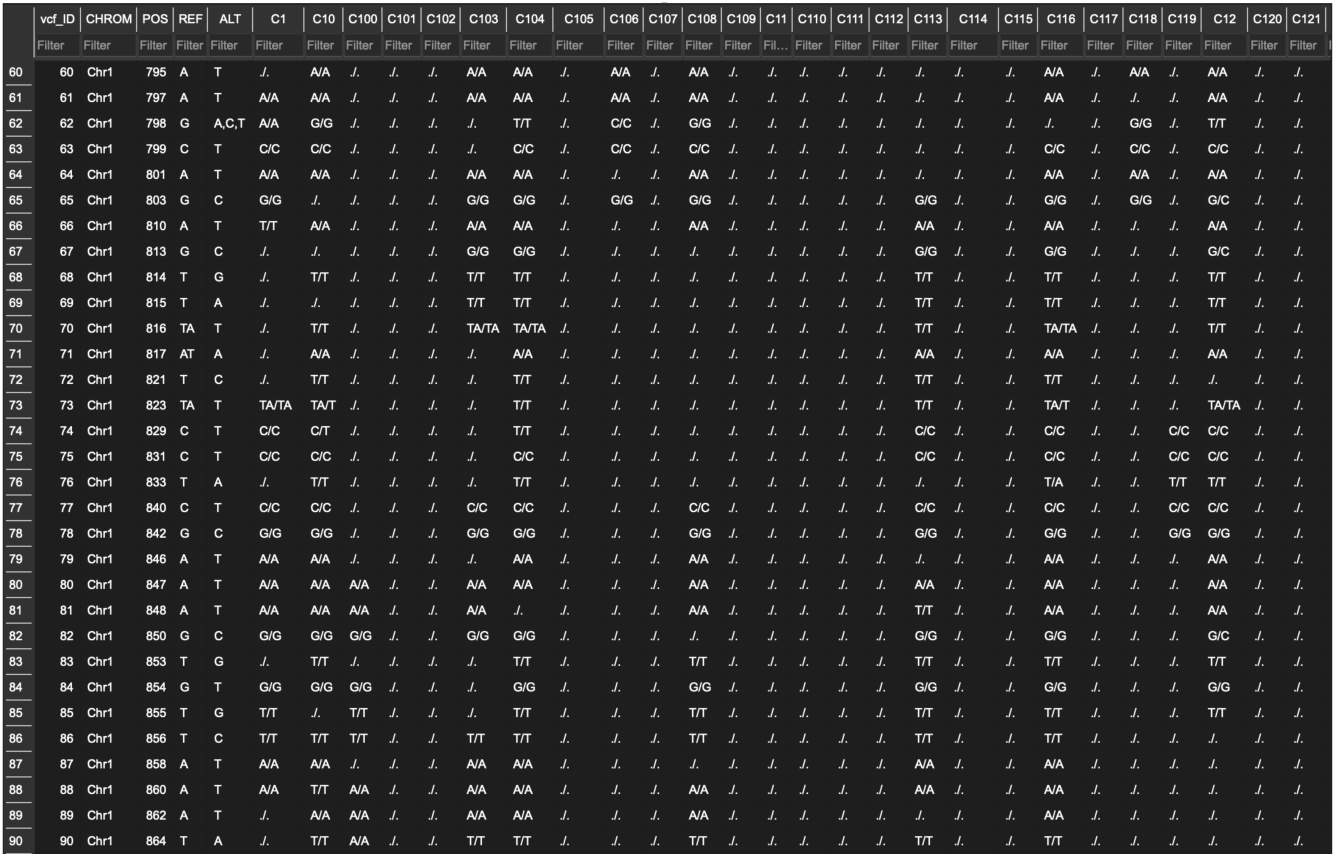
Part of Table vcf shown in DB Browser for SQLite.

Note that the same process was applied to the concatenation of all 8 original VCF files, which was about 592 GB, on our CentOS server to get the complete Table VCF.

### 4.7 Constructing vcf_jump_table for Faster Access

In our case, there were about 150 million rows of VCF data in the complete dataset from Alfalfa. However, our query inputs were supposed to be seqid and POS values, instead of vcf_ID, the Primary Key. This could largely harm the query efficiency because SQLite would go through all POS values within the Table for every VCF query. Generally speaking, for integer data columns like POS, adding an index to that column could improve query efficiency. Yet, in our case, although the POS values were monotonically increasing for each chromosome, the same POS value could appear multiple times at a different or even the same chromosome. For instance, within the first 500 MB of VCF data from Chromosome #1, there were 5,660 rows with POS values appeared more than once (as shown in **Figure 13**). This made indexing very complicated.

**Figure 13.**
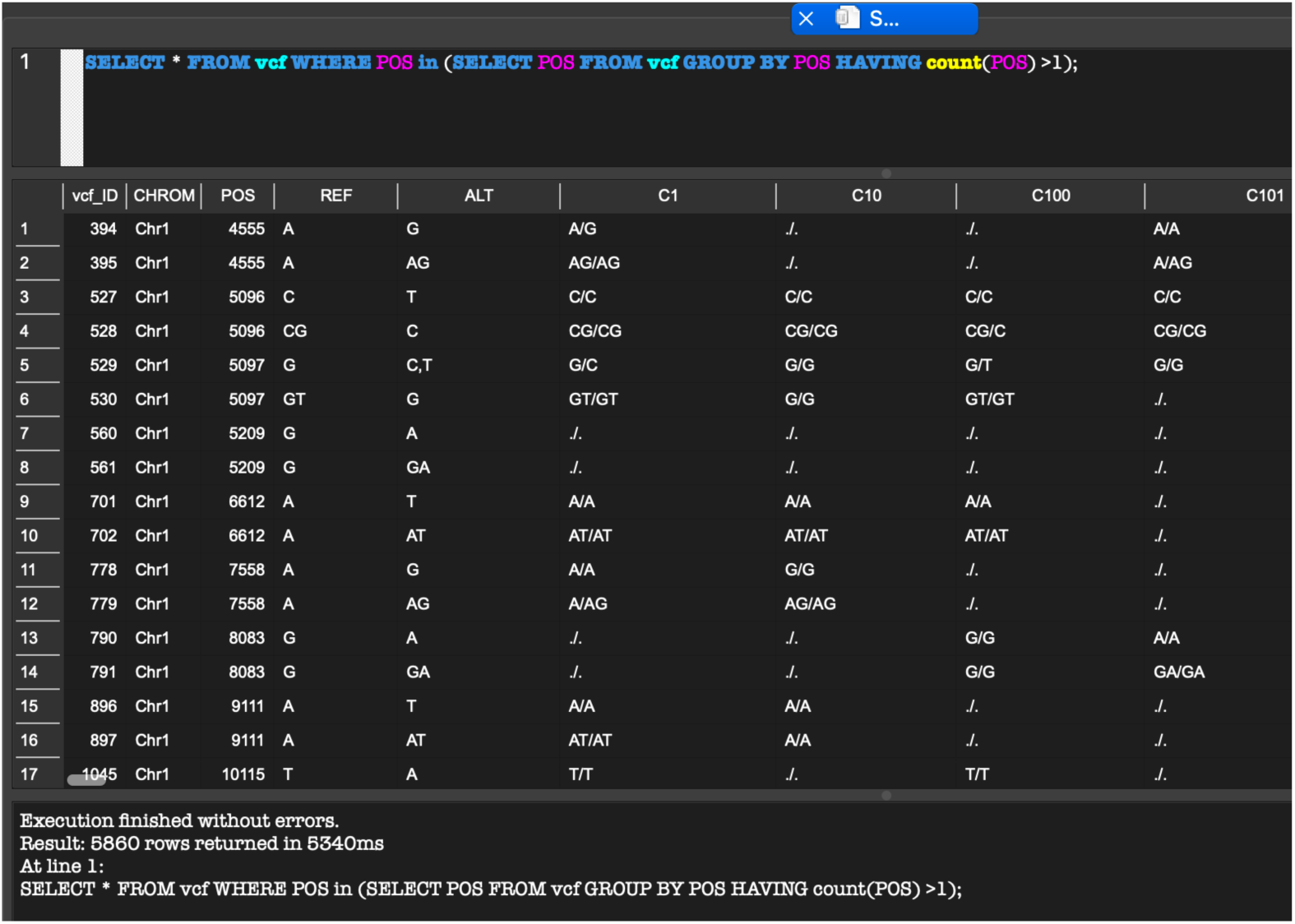
POS problem.

Thus, a jump table was created as follows:

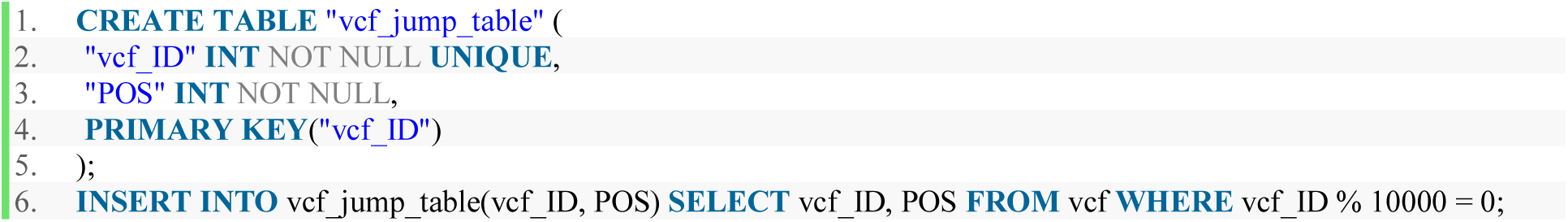

This way, for every 10,000 vcf_IDs, the first POS value would be stored in Table vcf_jump_table as shown in **Figure 14**. Searching the POS value being queried in this jump table could help the frontend application to limit the query range of vcf_ID from an entire chromosome to a small segment of it. The detailed query commands would be elaborated in Section 5.3.

**Figure 14.**
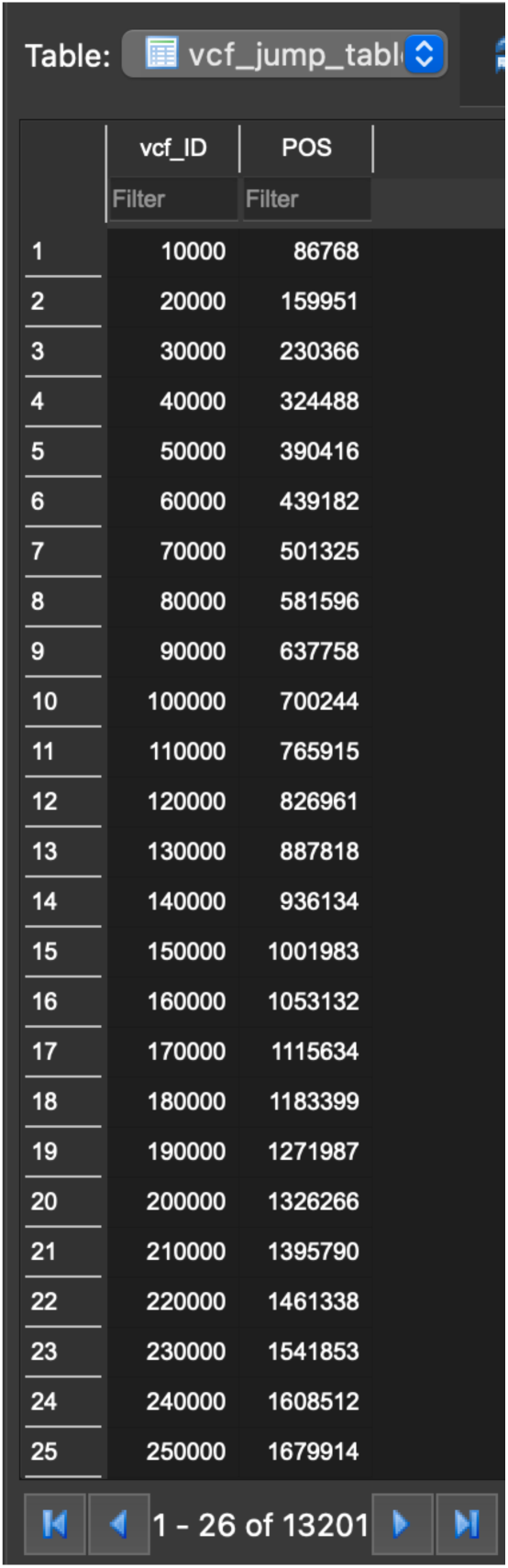
Part of Table vcf_jump_table shown in DB Browser for SQLite.

## 5. Methods: Frontend Flask Web Application

With the database file developed so far, we could move on to frontend design. In this section, we would refer to our three query modes as ID Query, Keywords Query, and VCF Query. The frontend web application should be capable to execute searches with user inputs from the website, giving correct and visualized query results, and handle improper inputs with detailed instructions. These functions were achieved under an isolated Python environment created by a Python package called “virtualenv”. The web application itself was developed under the Flask web framework, which was based on Jinja Templet Engine. Jinja enabled the Python program to get and pass data to the HTML page and render proper query results on the web pages shown to the users (Ronacher, n.d.).

### 5.1 Overall Look on the Web Project

As shown in **Figure 15**, the final project folder of the frontend web application contains a README document, the main Python program called “app.py”, a virtual environment folder, a static resource folder, and an HTML templates folder. Looking deeper into our folder levels, there were 8 txt files in the DNA folder containing the complete sequences of the 8 chromosomes of Alfalfa, which was copied from the intermediate datafiles created in Section 4.4. The backend database, Alfalfa.db, and two SQL scripts were stored in the folder “sql”. Those two scripts were used to get the marginal vcf_ID values for each chromosome’s VCF records. Moreover, the CSS file contained the format settings of our website and the three HTML templates were used interactively with the Python program to render the GUI of our database on a website. The complete scripts in the Python, CSS, and HTML files were shared in the Appendix section, and on our GitHub repository.

**Figure 15.**
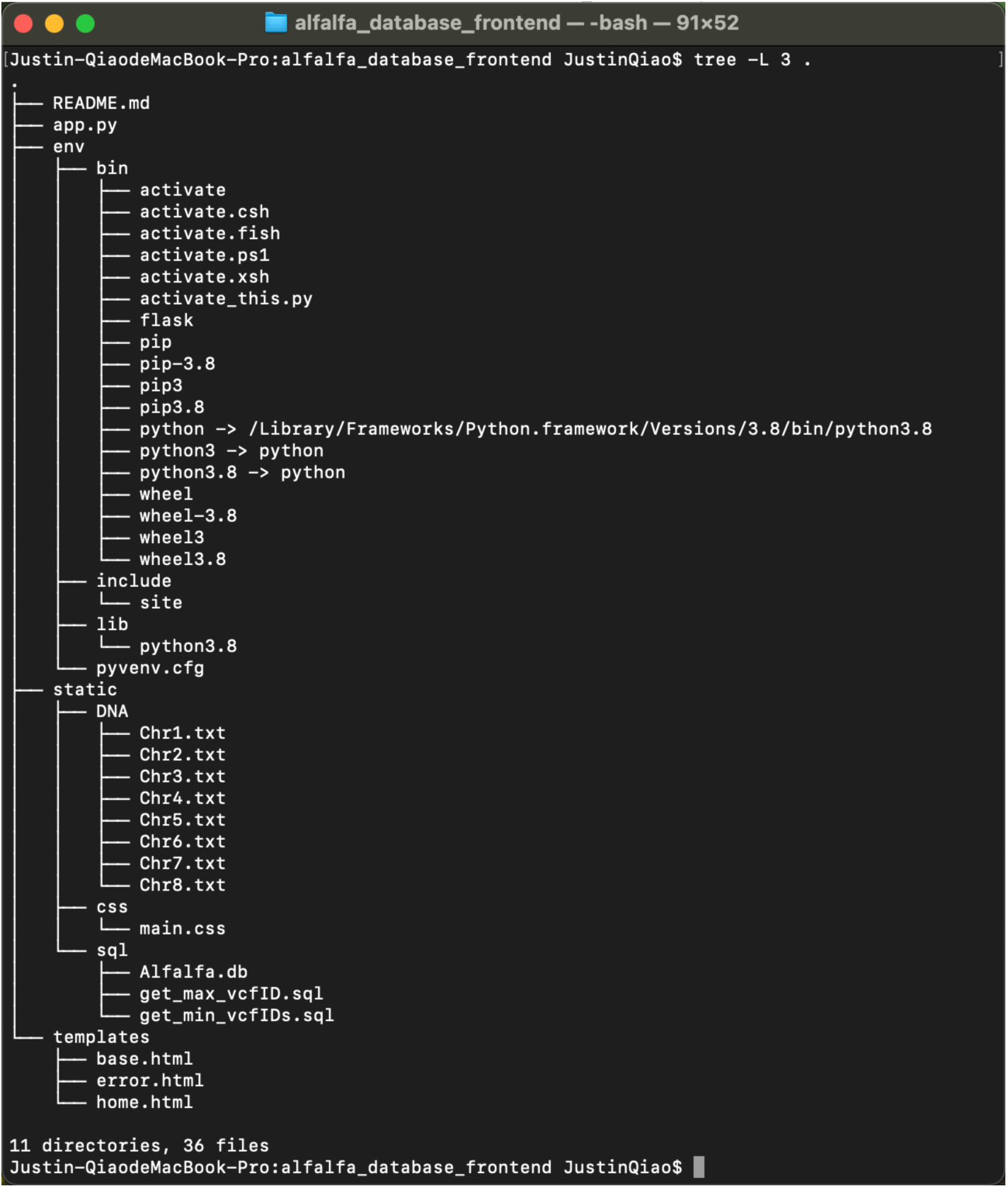
Final project folder of the frontend web application.

### 5.2 SQL Query Statements

For ID Queries, the frontend application was required to SELECT all information about a gene ID from Table annotate_info, pros_info, gff, alltpm, and the other five Tables containing annotations from other databases.

Retrieving annotate_info, pros_info, and alltpm data were straightforward. The following SQL SELECT commands could be used to take out all rows in those three Tables associated with the given ID (using MsG1A10000001 as an example):

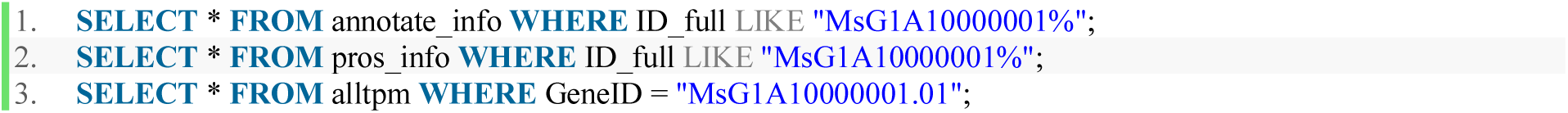

To make querying GFF information more efficient, an implicit JOIN instruction was utilized:

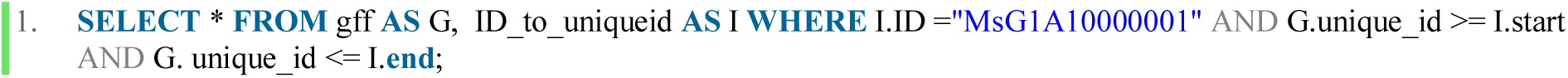

Noticed that the “ID” column in Table ID_to_uniqueid only contained the 13- character primary ID, so that we used “I.ID =” here without the “%” wildcard. This way, the query speed could be improved.

Moreover, explicit LEFT OUTER JOIN was used to find annotations from other databases:

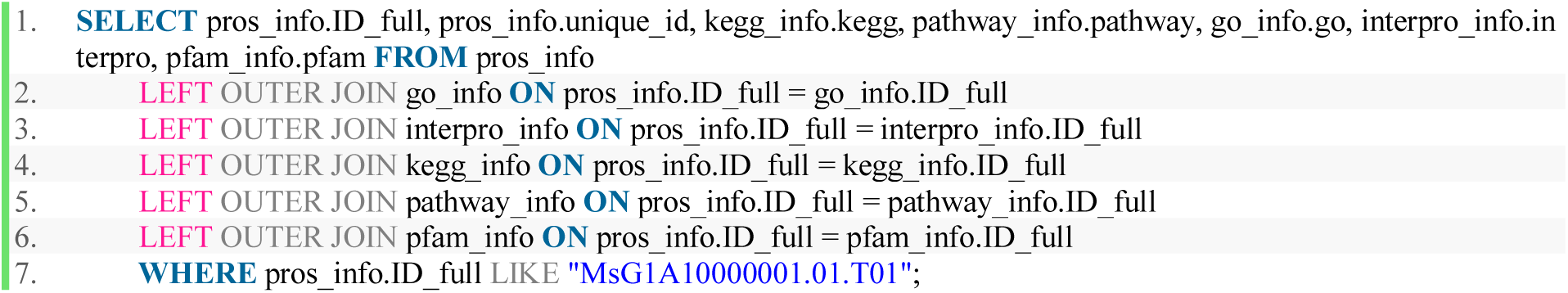

This SELECT statement could give us the correct result because all possible ID_full was stored in Table pros_info, and if some of the Tables did not contain the provided gene ID, the NULL character would be used as a placeholder because of the property of OUTER JOINs. If the gene ID was not included in any of these five Table, the frontend applications could detect those NULL characters and hide them respectively.

The Keywords Query mode could be restated as follows: if a row of data in Table annotate_info contains the entered keywords as a sub-string of the data string stored in column “info1” and/or column “info2”, the entire row should be returned. Fuzzy searches like this could be managed by the wildcard symbol, %, in SQL. The following SELECT statement was created to achieve these goals (using keywords “HSF” as an example):

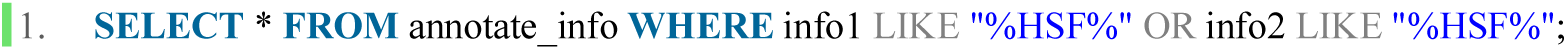

Finally, as we planned in Section 4.7, we use two cascading searches for VCF Queries. Since HTML rendering could be time-consuming, we restricted the query range of POS to be between 1 base pair (bp) to 3,000bp. This could be easily changed by modifying the if statement on Line 257 of app.py. Additionally, we were required to know the first and last vcf_ID of each different chromosome. These values were obtained by running get_max_vcfID.sql:

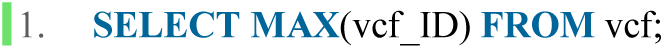

and get_min_vcfIDs.sql:

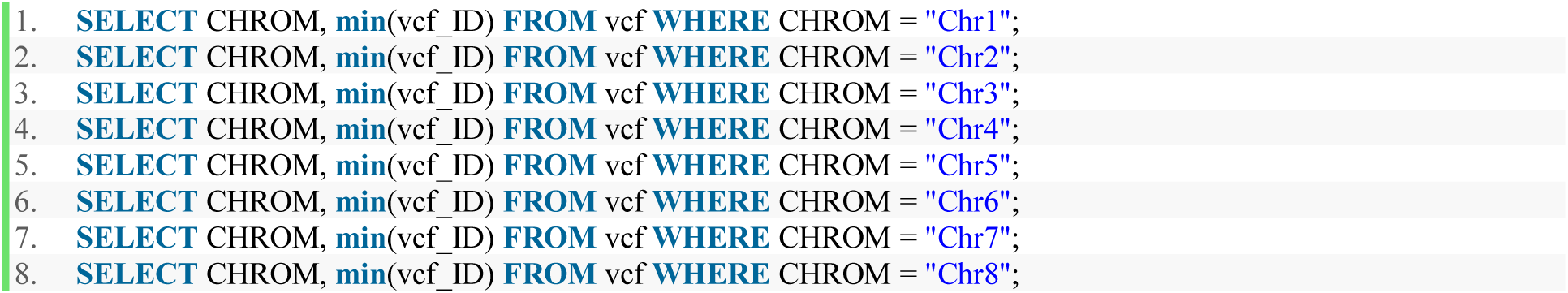

over the complete database file with VCF data from all of the 8 chromosomes on our CentOS server. These SQL query results were stored as pre-defined values in our frontend code on Line 264 and 265 in app.py.

When an actual VCF Query happens, our frontend application first executes the following two SQL queries on Table vcf_jump_table:

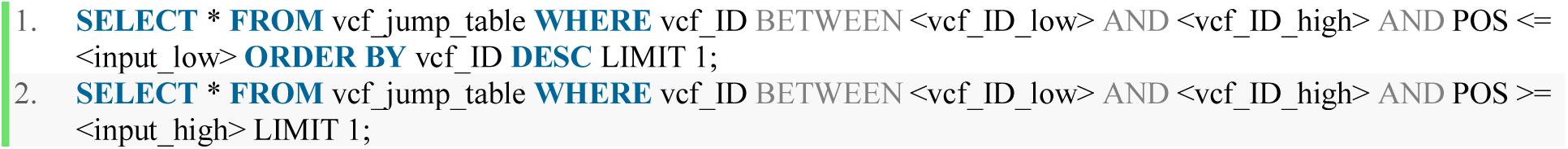

<VCF_ID_LOW> and <VCF_ID_HIGH> were obtained from the pre-defined array, matching to the specified seqid selected by the user, and <INPUT_LOW> and <INPUT_HIGH> were the specified POS range. Since we had limited the POS range to be less than or equals to 3,000bp, these two queries were likely to return a query range of vcf_ID less than 20,000 rows. This narrowed a VCF Query task from one search over about 20-millions of rows down to two queries over tens of thousands of rows per query. It also stops the application from going deeper on unrelated VCF records to look for POS values within the specified range.

Finally, we used the value returned above, vcf_ID_lower_bound and vcf_ID_upper_bound to pull out the section of rows on Table vcf whose POS value were between <INPUT_LOW> and <INPUT_HIGH>:

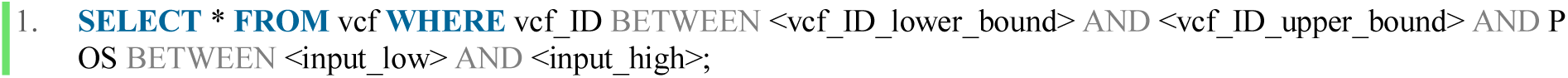

### 5.3 Frontend Python Application

The Python application, app.py, was responsible to connect to the backend database, execute queries according to the HTML inputs, and render HTML templates with the visualized query results. There are four URL routes controlled by this program: “/”, “/ID/”, “/annotation/”, and “/VCF/”.

To begin with, the home route, “/”, renders the default home page with user inputs placeholders, as defined in Section 10.5 on Line 9 through 57. Whenever the user refreshes the page or clicks the reset button, the application should be redirected to this route and render this page as shown in **Figure 16**.

**Figure 16.**
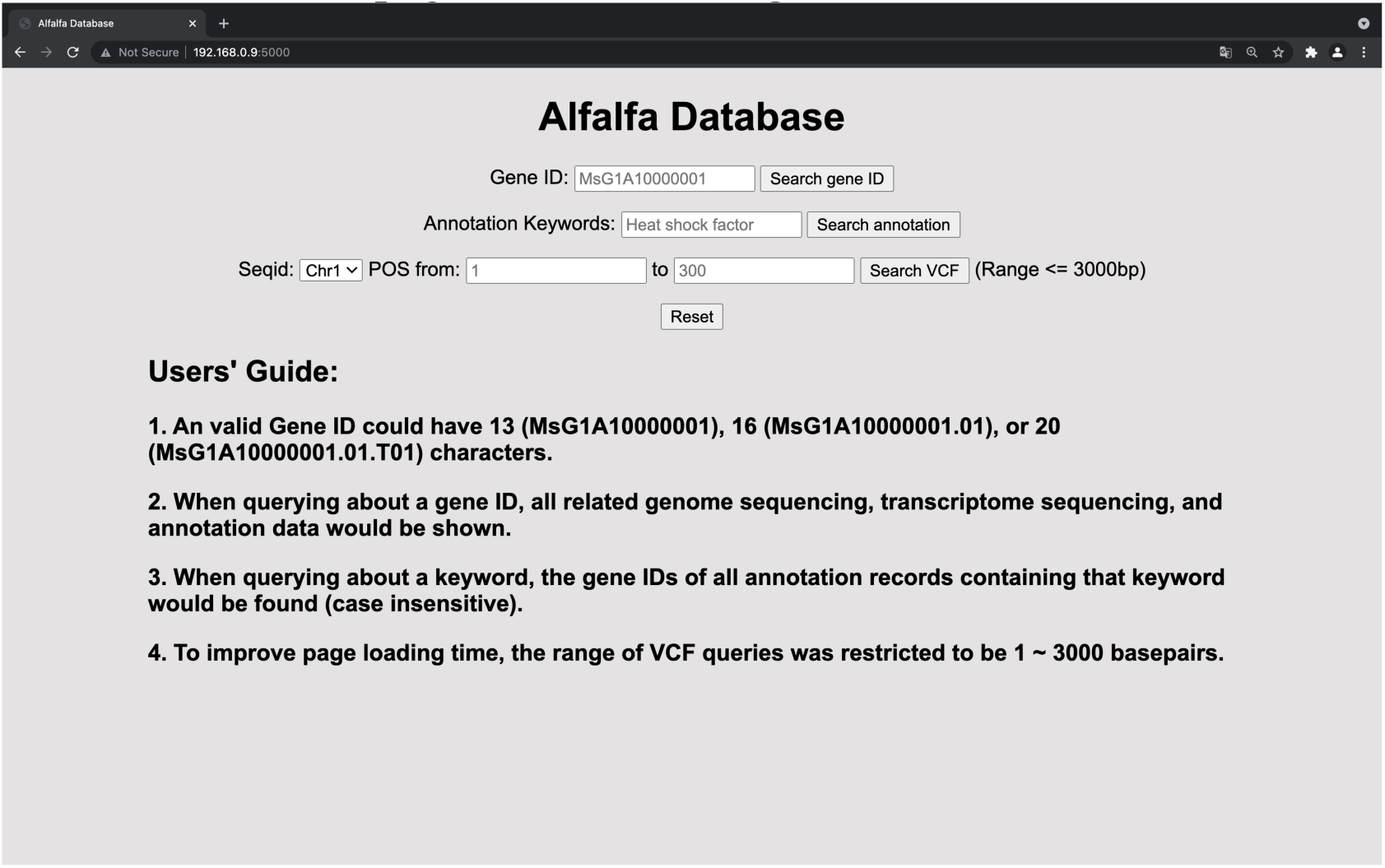
Web application home page.

Secondly, route “/ID/” manages ID Queries. The only input is obtained from the text box on the top of the home page. As mentioned before, a valid gene ID could only have 13, 16, or 20 characters for our dataset. Thus, any input string with a different length is rejected and the error.html page, fully defined in Section 10.5, should be rendered with a proper error message. If the route is not returned at Line 33 of app.py as defined in Section 10.3, it would be connected to Alfalfa.db using the Python package called “sqlite3”, and then executes queries over all SQL Tables except Table vcf as planned in Section 5.2. There are flag variables such as show_annotate and have_info for the HTML templates to hide “None” results for different SQL queries. Note that Table pros_info is searched first on Line 38 through 44 because all valid ID_full were stored in this Table. If Table pros_info does not contain the gene ID from the HTML user input, the error page should be rendered and inform the user that there is no information found about the given gene ID. Further, while querying from Table go_info, interpro_info, kegg_info, pathway_info, and pfam_info, the concatenated data are divided to list objects, so that each annotation entry could be correctly hyperlinked to their source pages from other databases’ websites. Moreover, while querying GFF data, many resulting GFF entries’ “strand” value is “-”. In such cases, the reverse complement DNA sequence in the “DNA” column should be printed to the web page. On the other hand, if the “strand” value is “+”, directly printing the DNA sequence stored in the backend database would be sufficient. To achieve this, a helper method, get_conjugate, was defined on Line 6 through 19 and utilized on Line 147 through 152. Next, all CDS sequences in the GFF results would be concatenated together on Line 154 through 166 for better user references. Finally, expression quantity data about the queried Primary ID stored in Table alltpm are stored in a Python list object, as shown on Line 168 through 174. The third route, “/annotation/”, executes queries over Table annotate_info. Note that there could be more than one annotation entry attached to the same 20-character gene ID because those annotations could come from different source databases of *Medicago Truncatula*. Thus, a rowspan attribute is calculated for each row in the query results, and added to the end of each tuple object in ans_list, as shown on Line 208 through 236. For each ID_full in the result set, the top row has a rowspan value equals to the number of all rows attached to that ID_full, and the rest of rows’ rowspan values were set to 0. This way, the HTML page could only print each unique ID_full only once.

Finally, route “/VCF/” returns the proper section of Table vcf on the seqid specified in the drop-down menu, as shown in **Figure 16**, between the specified POS range. The complete DNA sequence being queried would be extracted from the proper txt file stored in the folder “DNA”, using a similar method as discussed in Section 4.4.

### 5.4 Frontend HTML Templates and Cascading Style Sheets (CSS)

The HTML templates and the CSS file were responsible to interact with users via the website, collect user inputs, format and visualize query results passed back from the Python program, and add hyperlinks both internally and externally to other existing online databases. Since the Jinja template engine is extensible, we could inherit HTML codes and settings from other parental HTML templates. In our case study, both the home.html and error.html templates inherited codes from base.html, as shown in Section 10.4 through 10.6. Since error.html renders the same user interface, with only an additional error message area, as home.html does in its reset state as shown in **Figure 16**, only the design of home.html with main.css is elaborated in this section. Screenshots of sample queries would be shared below in Section 6.

Line 9 through Line 108 of home.html are responsible to render the home page, shown placeholders at reset, and echo the most recent query inputs. Note that echoing text inputs with empty space characters and remembering dropdown menu selection might require help from JavaScript to process the input value passed back from the main Python application, as shown on Line 68 through 79 and Line 86 through 94.

Further, for query mode A, ID Query, we visualized our query results into four separate tables and a bar chart. Those tables were put into a CSS class called “container” to force the query result window to stay at an acceptable height. Specifically, Line 114 through 171 in home.html could create a table of hyperlinks. Any KEGG, Pathway, GO, Interpro, and Pfam annotation on the gene being queried would be shown with clickable links to their source databases’ websites. A second resulting table would contain annotation keywords from those outside databases. Below the second table, a bar chart, powered by Apache Echarts (n.d.), should be rendered to visualize the expression quantity as defined on Line 199 through 243 in home.html. Moreover, a third table would print the complete protein sequence and CDS sequence for each queried ID_full, and the last table should contain the complete GFF records related to the given gene ID. Note that if any of these tables have no meaningful content, the table headers would be hidden accordingly to the flag variables passed from app.py. To make the tables resulting from ID Queries more human-readable, all DNA and protein sequences were defined to be in a “seq” class. The CSS file had regulated objects in this class to shown its text content completely, as defined on Line 36 through 40 so that users could have a better impression of the length and characteristics of the sequences they found. Additionally, we had taken advantage of the “scroll” mode for “x-overflow” and “y- overflow” in main.css to prevent long text data from occupying nearby data cells. In addition, in the Keywords Query mode, Line 298 through 323 in home.html was used to format the resulting table according to the rowspan attribute passed from app.py. For each unique ID_full in the result set, internal links to its ID Query result were managed by hidden input clauses on Line 312.

Finally, VCF Queries were taken care of by Line 329 through 578. The input parameters and their corresponding complete DNA segment were designed to be shown in a separated table defined on Line 332 through 354. The resulting VCF table would contain all VCF data within the specified range, and data cell coloring was implemented to improve data visualization. Specifically, the reference (REF) gene type was colored green, the heterozygous gene type with a REF strand was colored orange, the heterozygous gene type with no REF strand was colored yellow, and the homozygous mutation type was colored pink. This way, researchers could quickly detect the genotypes and patterns they are looking for. Note that the resulting VCF table was given a style “user-select: all” on Line 359. This is critical to make copying the entire resulting set and pasting to Excel spreadsheets more convenient.

## 6. Results

Overall, the entire project flowed as shown in **Figure 17**. The resulting web application perfectly satisfied our design requirements. The three query modes, ID Query, Keywords Query, and VCF Query were all managed properly with good preciseness, intuitive visualization, and detailed exception handling. The complete project folder was shown in **Figure 15**, and the default home page was shown in **Figure 16** in the previous section. In this section, sample queries are shown as the result of our study on Alfalfa. A demo video of our web application could also be found at: https://youtu.be/Djr81po206A.

**Figure 17.**
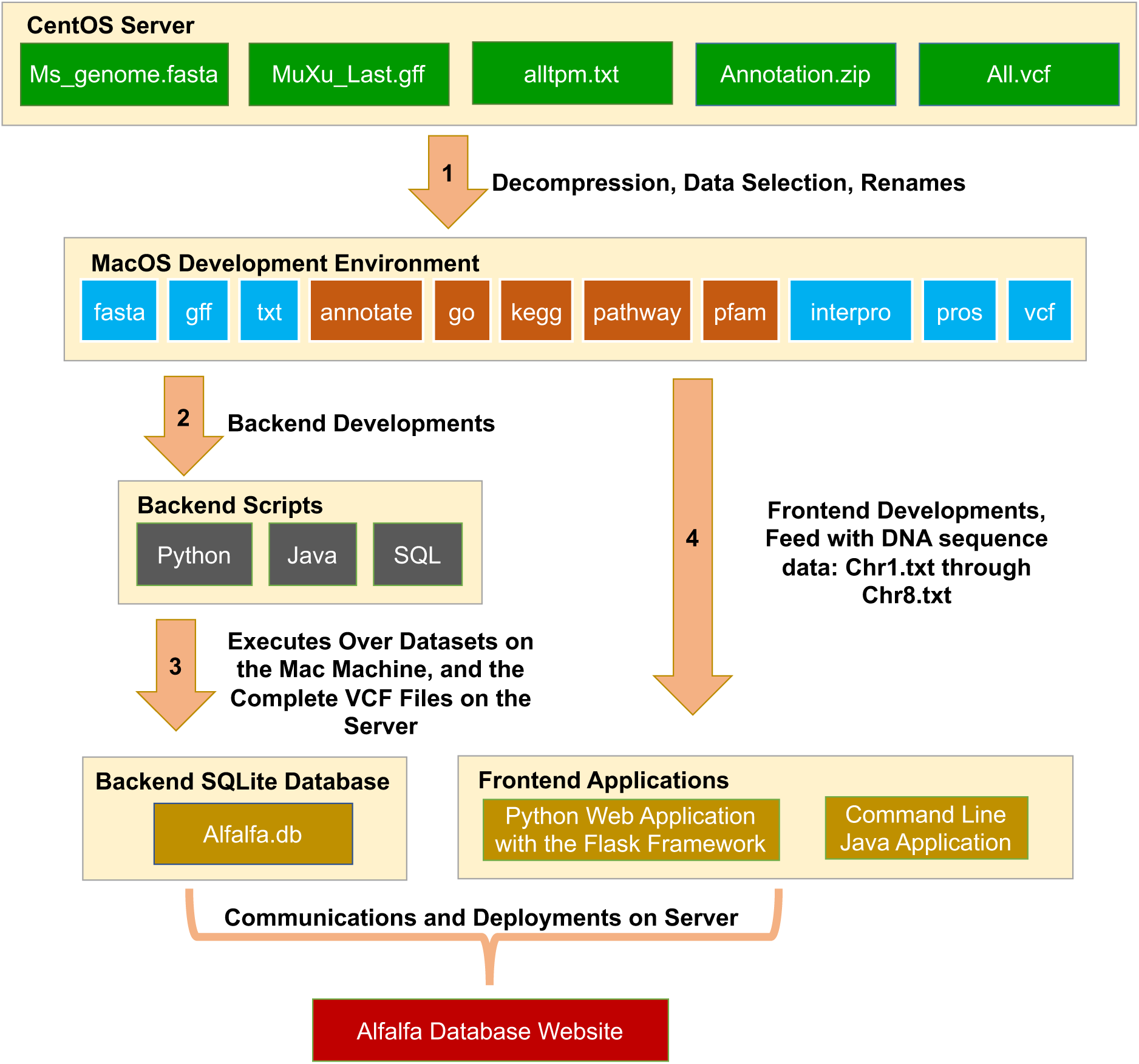
Flow Chart of the entire project.

To begin with, **Figure 18** through **Figure 21** present the result of a 16-character ID Query. All hyperlinks to external online databases work as excepted, and all information related to the given ID in our raw dataset are retrieved precisely.

**Figure 18.**
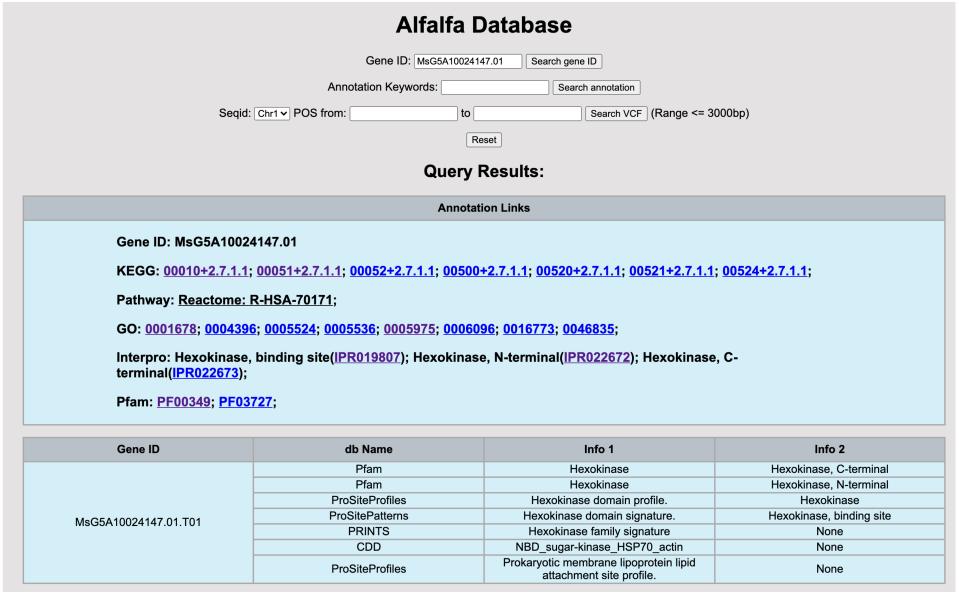
Sample 16-character ID Query, Part 1.

**Figure 19.**
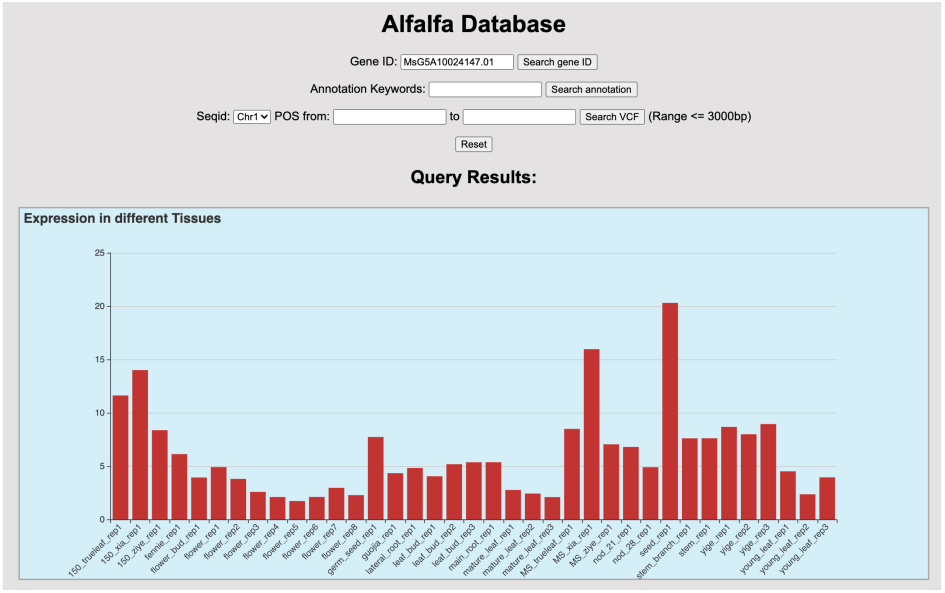
Sample 16-character ID Query, Part 2.

**Figure 20.**
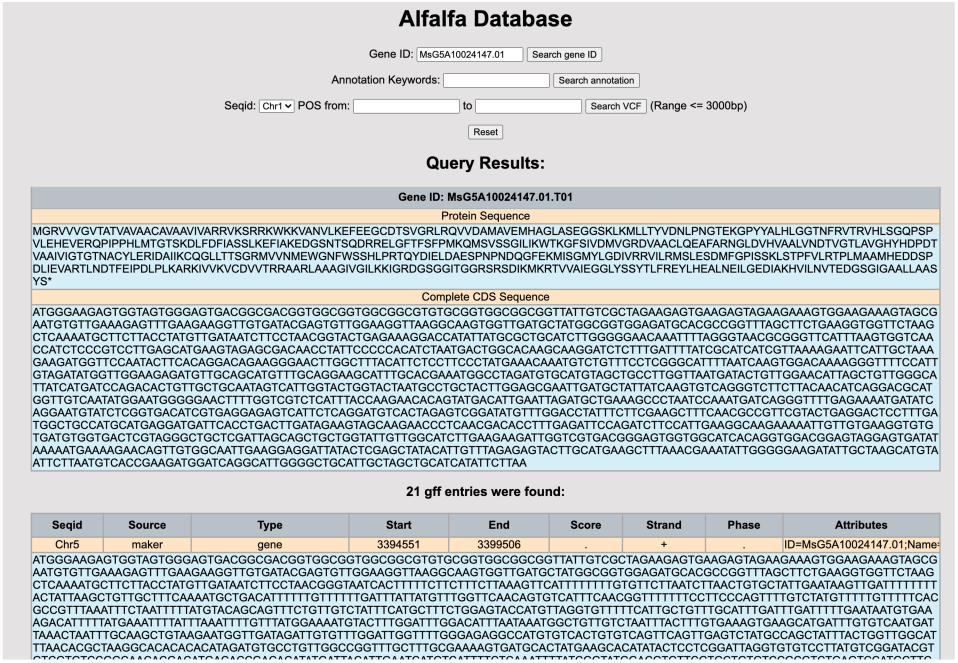
Sample 16-character ID Query, Part 3.

**Figure 21.**
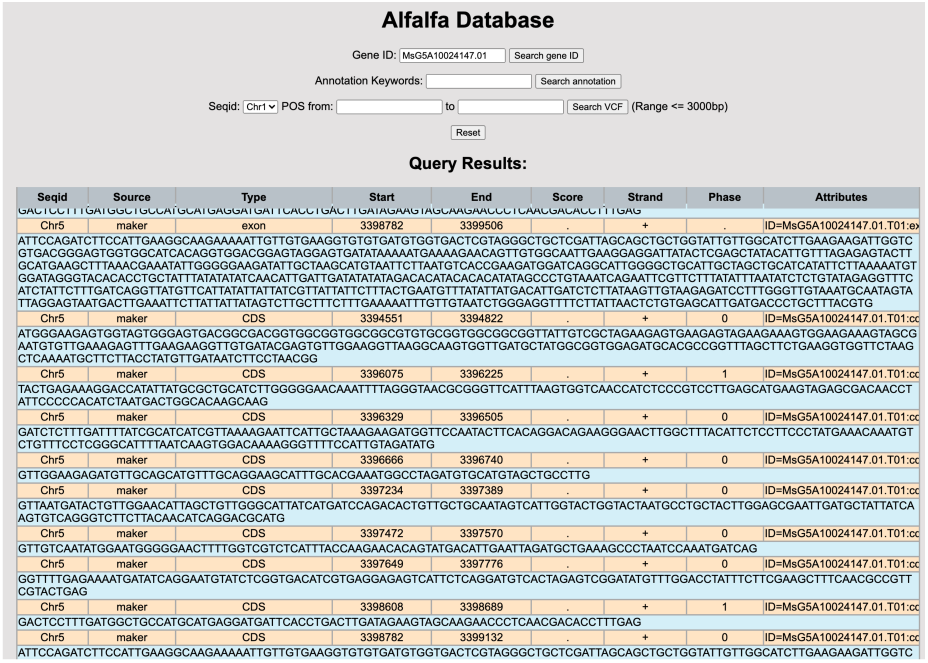
Sample 16-character ID Query, Part 4.

The resulting tables are all flexible. When there are more than one sequences related to a given gene ID with 13 or 16 characters, the complete CDS and protein sequences table would incorporate all of those records (**Figure 23**). On the other hand, any Null or None characters in the query result would be hidden as expected (**Figure 22**).

**Figure 22.**
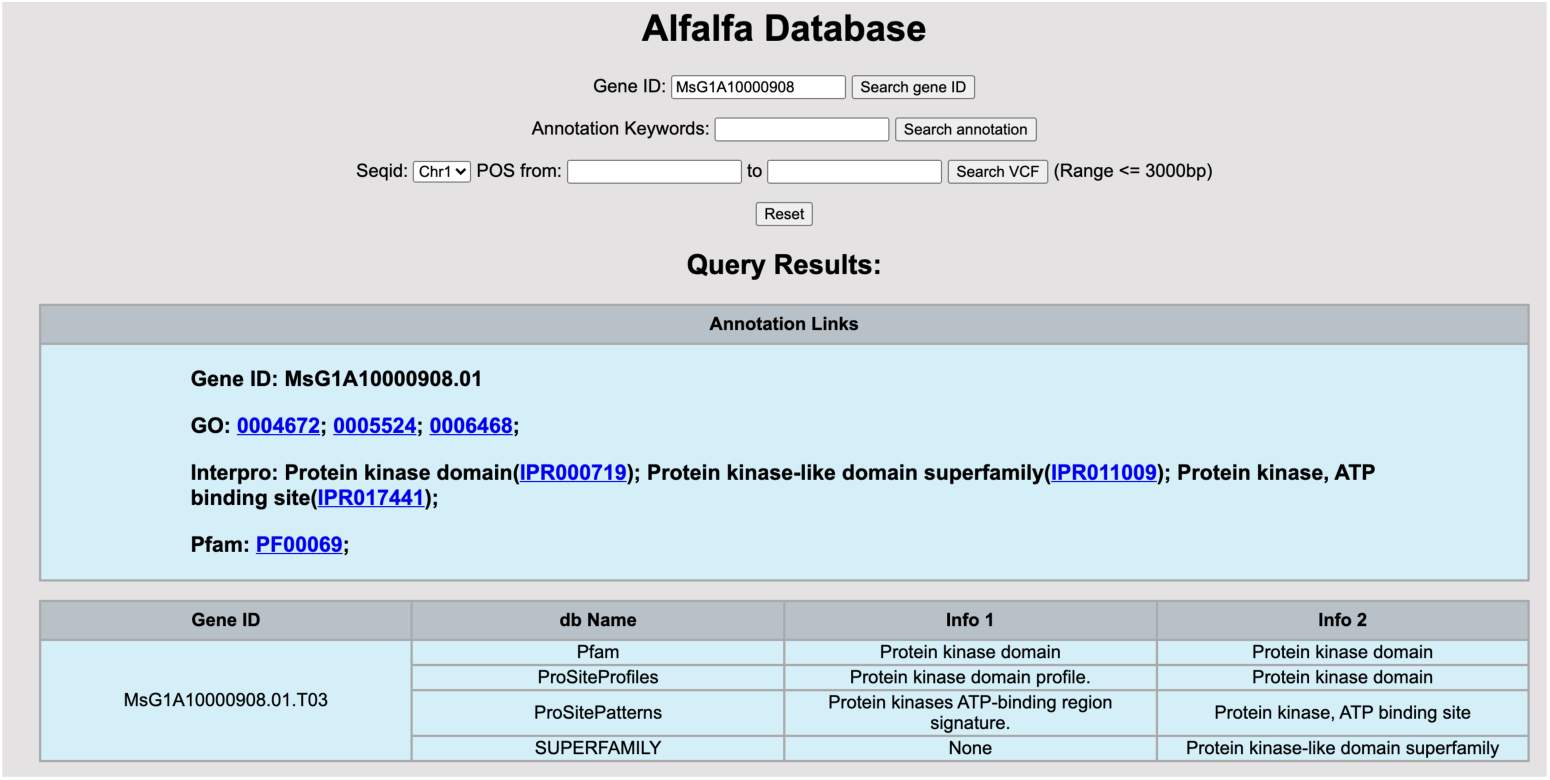
Sample 13-character ID Query, Part 1.

**Figure 23.**
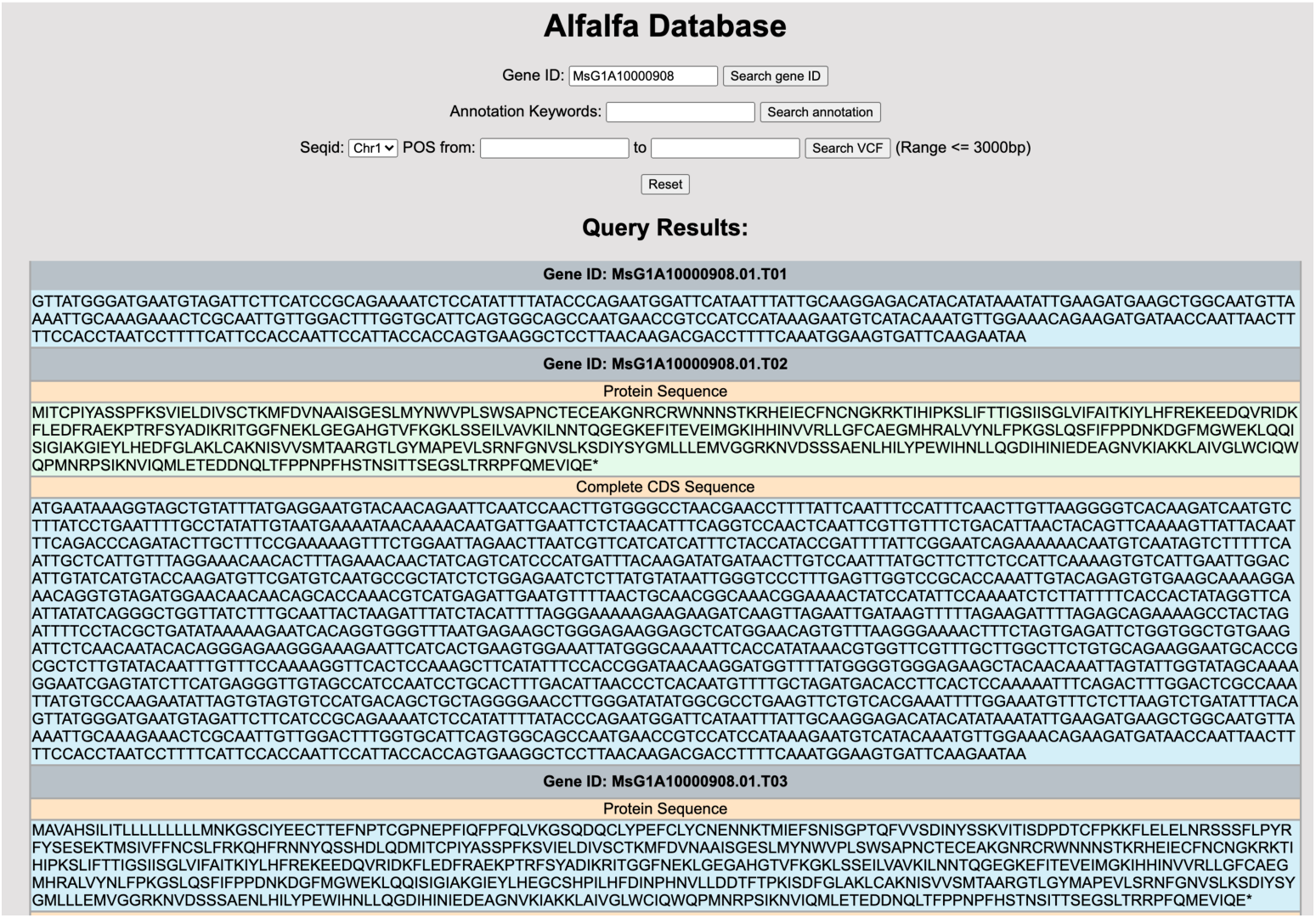
Sample 13-character ID Query, Part 2.

Moreover, if a 20-character gene ID is searched, only that specified sequence is shown in the resulting table. Any other sequences or GFF entries from the same primary gene ID are hidden, as shown in **Figure 24**.

**Figure 24.**
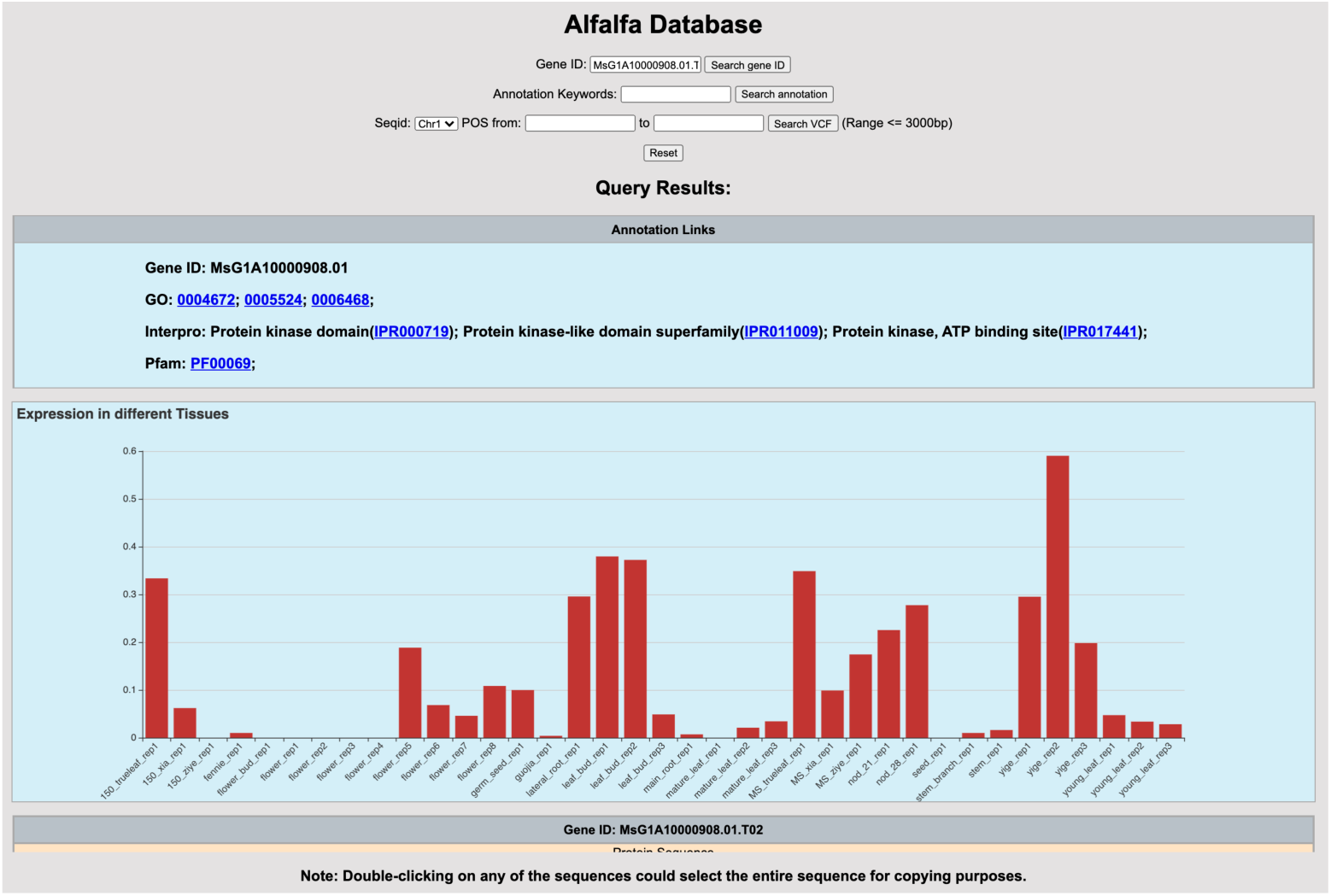
Sample 20-character ID Query.

Secondly, for Keywords Queries, the internal links to ID Query results and the combined data cell formats are rendered as we planned (**Figure 25**).

**Figure 25.**
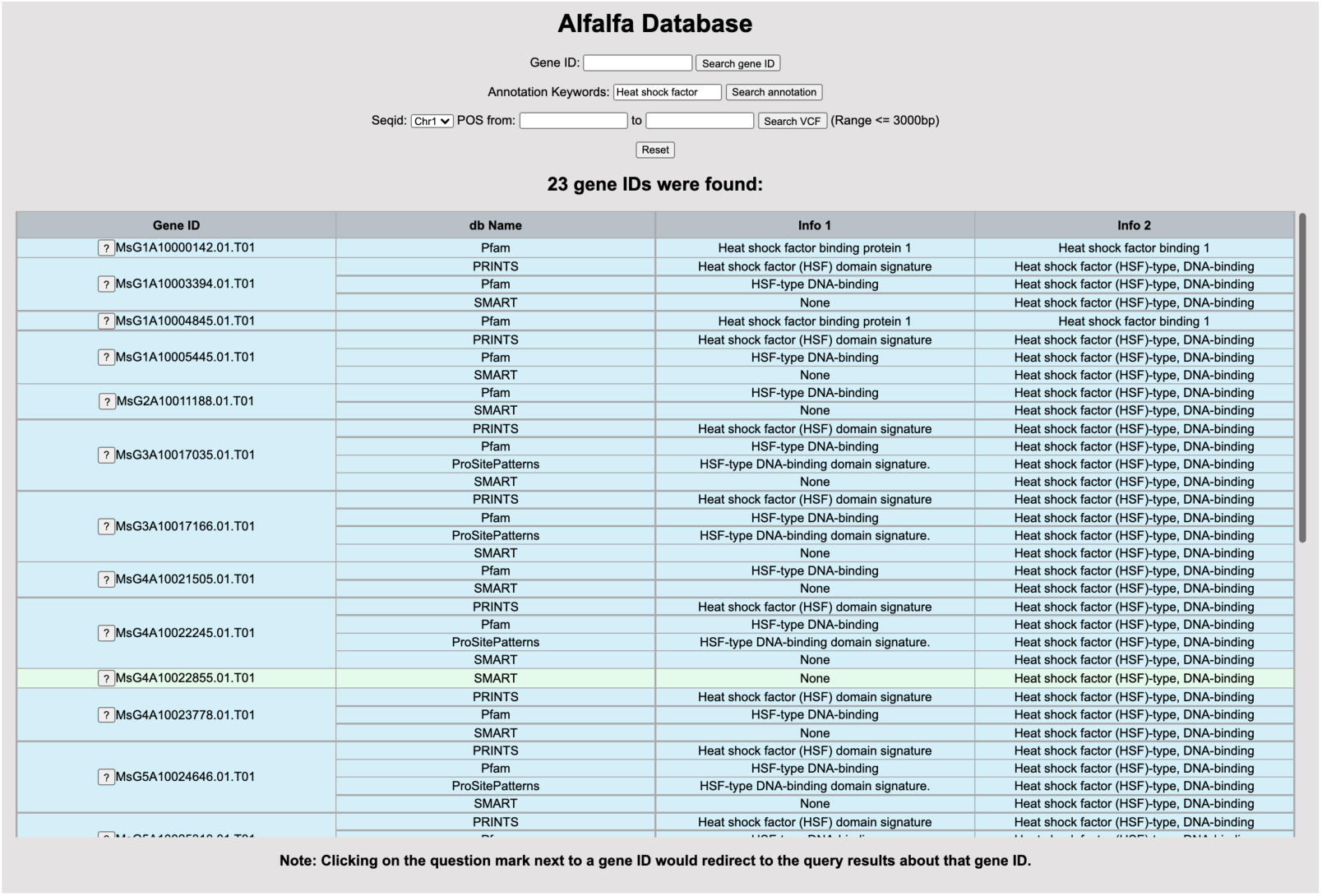
Sample Keywords Query.

Moreover, in VCF Query mode, the expected portion of the complete VCF table is returned (**Figure 26**). The nucleotides at each POS are visualized precisely, and one-taping copying from the resulting VCF table to Excel spreadsheets is supported.

**Figure 26.**
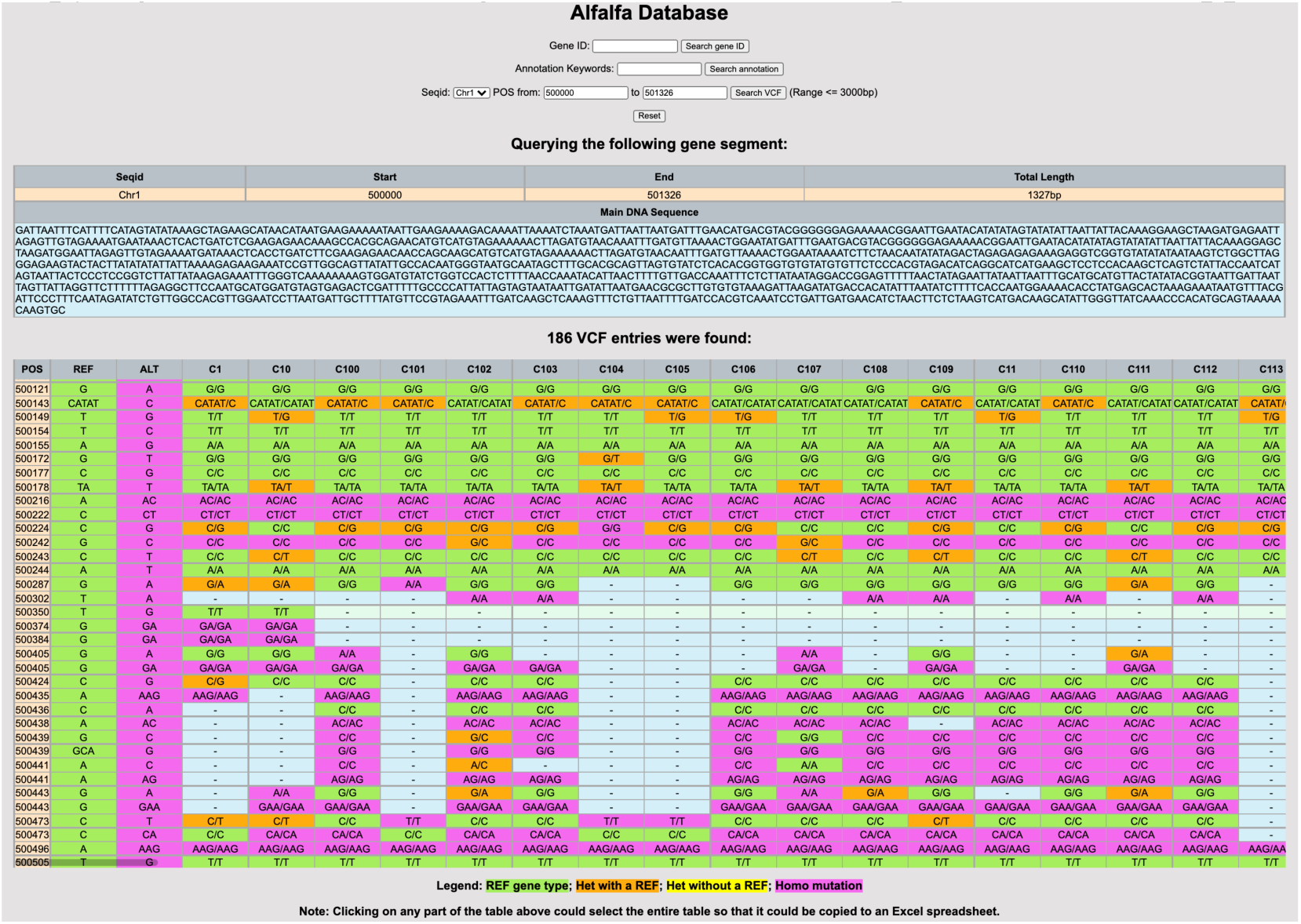
Sample VCF Query.

Finally, if any user inputs were invalid, or if there was no result found, error.html was rendered with the proper error message, as shown in **Figure 27** and **Figure 28**.

**Figure 27.**
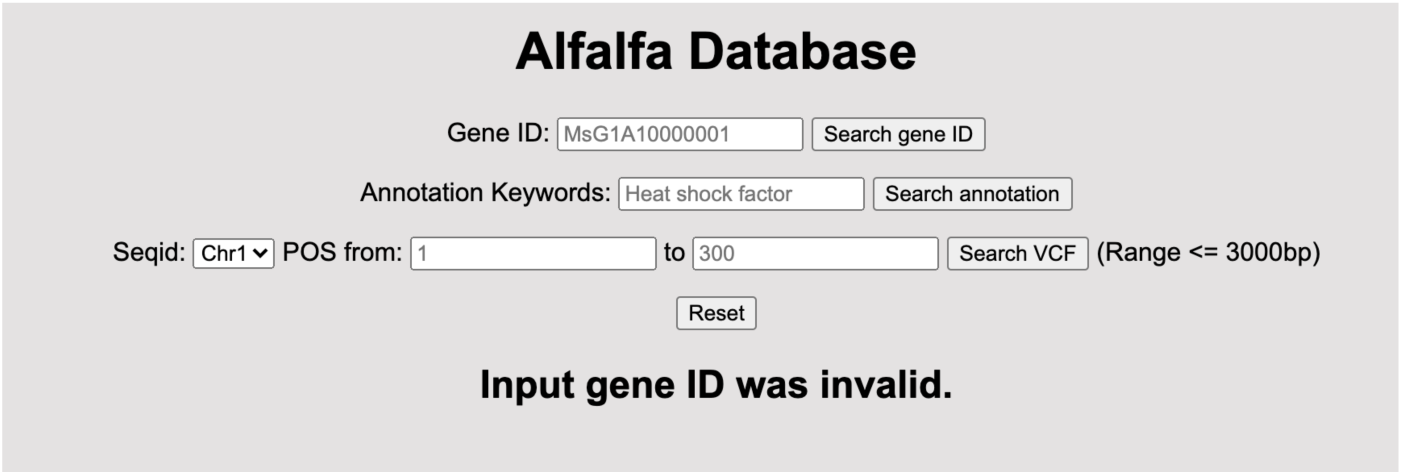
Sample ID Query with invalid ID input.

**Figure 28.**
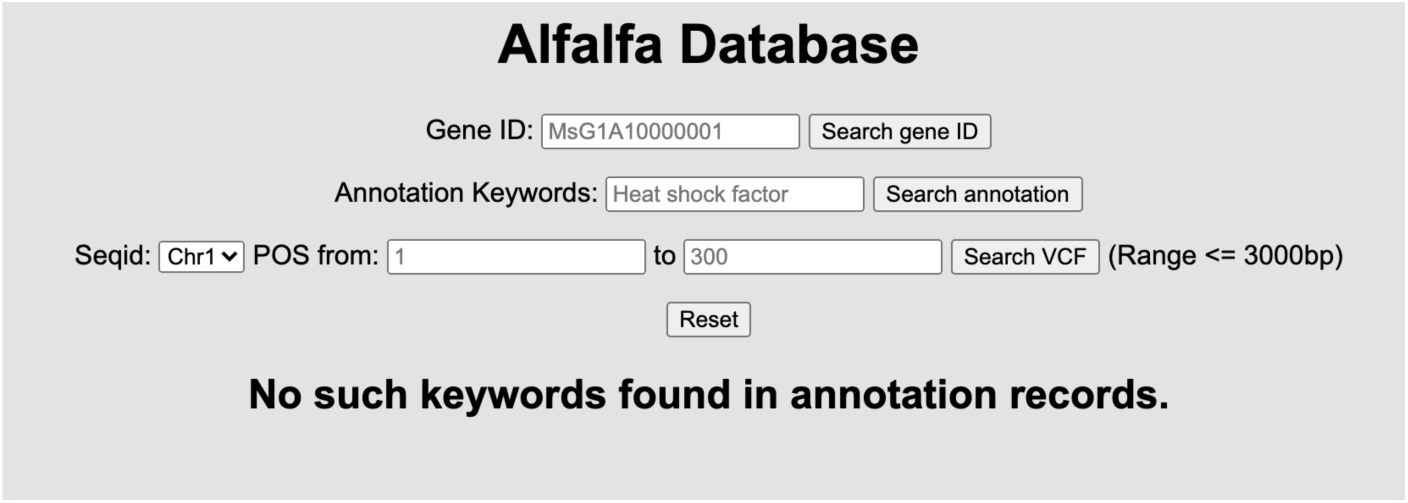
Sample Keyword Query with no matched result.

Lastly, the static resource folder of our final product is shown in **Figure 29**. Our backend SQLite database file not only made the original dataset more structured and portable, but also compressed the dataset from near 600 GB down to 114 GB.

**Figure 29.**
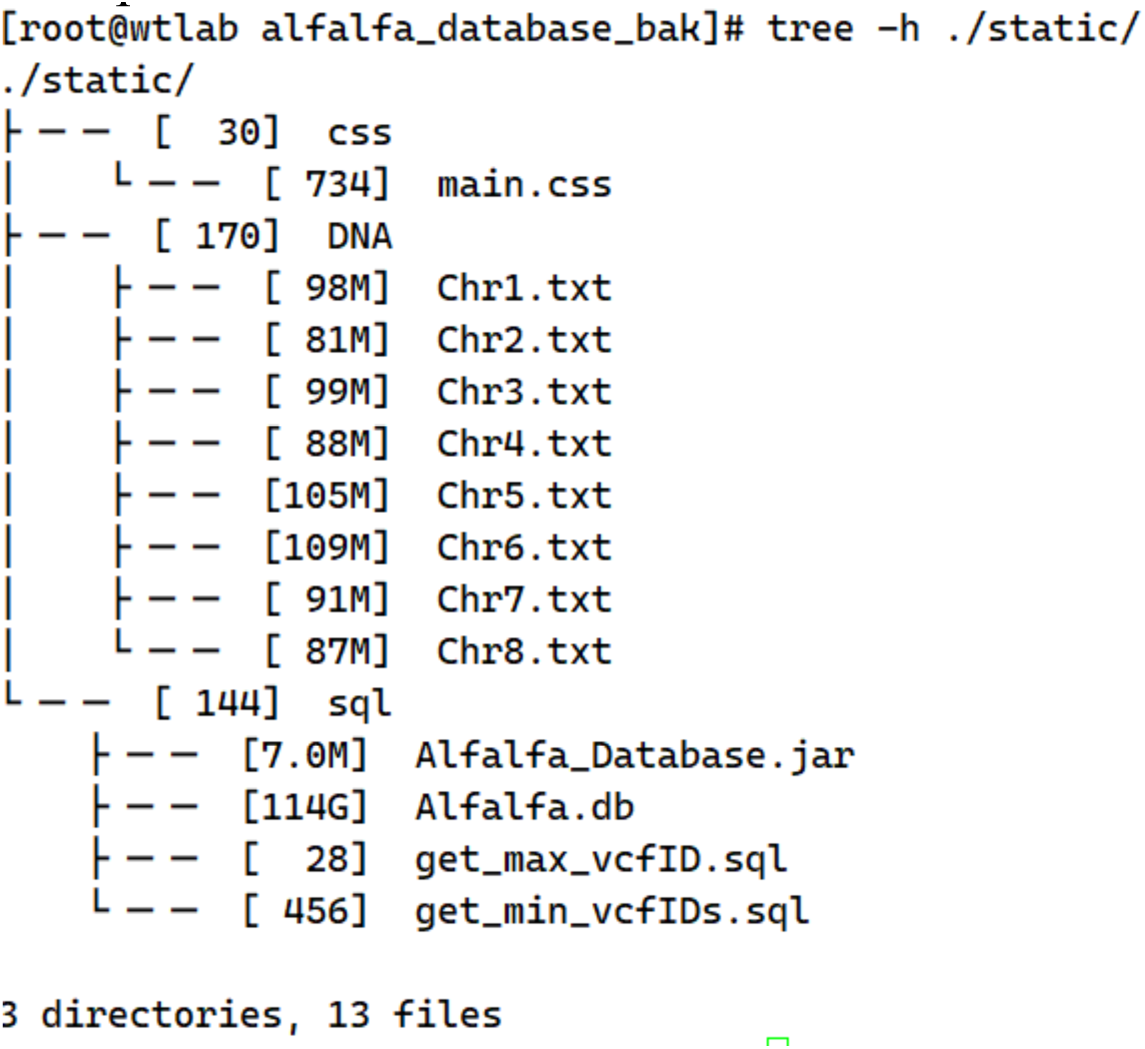
Resulting static resource folder.

In all, our Alfalfa Database web application is the first comprehensive, online, bioinformatical database of *Medicago sativa*. Although the product is still under testing on our local area network, the development process and scripts used along the procedure have referential significance to develop online databases of other species. from similar text-based datasets.

## 7. Discussions

Although all of our Python, SQL, and Java scripts worked smoothly while constructing the database’s backend and deploying the web frontend, we have also discovered some potential improvements.

### 7.1 Basic Improvements on Our Procedure

Firstly, the process of converting TSV annotations to SQL INSERT scripts could be improved. In our case study, we changed the raw data files’ extension to “.txt”, opened them in Excel spreadsheets, and used the CONCATENATE function in Excel to generates the SQL instructions. However, when the annotation file is large enough, Excel may crash when applying the formula to millions of data cells at the same time on a personal computer. In this case, using a Python script to process the TSVs and generate SQL commands directly could be a more stable way. Drabas (2016) has provided a convenient way to read and write TSV files with a Python package called “pandas”. With the help from that package, scripts similar to what we used to generate the UPDATE statements in Section 4.4 could be used to process the raw data.

In addition, sorting some of the raw data could improve the processing time. For instance, when we truncated the main FASTA file into DNA sequences, if Table gff was sorted by seqid, it would be no longer necessary to open and close text files for every GFF entry. Instead, the text files could be opened only once in a predefined order, such as an alphabetical one, and a file would only be closed once after processing the entire continuous block of GFF entries from that corresponding DNA sequence. This way, thousands of unnecessary file-open and file-close operations could be prevented.

### 7.2 R and RStudio

In addition to Python, R is another popular programming language for statistical analysis developed by the R Foundation, and RStudio is a widely used IDE for R (The R Foundation, n.d.; RStudio PBC, 2021). Some R packages, such as Biostrings (**Figure 30**) and vcfR (**Figure 31**) could give researchers a statistical view on the massive raw Genome Sequencing and Transcriptome data (GFF and FASTA files). In this study, these statistics were used to check the programs’ correctness. For future studies and developments, vcfR could be a powerful tool to manipulate, visualize, and increase the readability of the information stored in VCF files.

**Figure 30.**
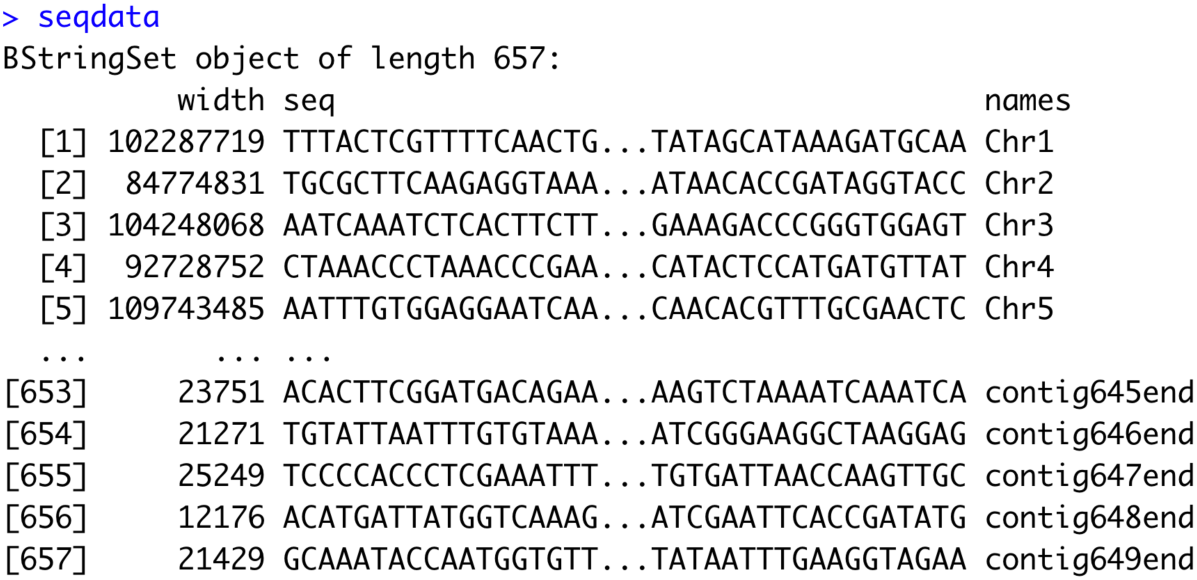
Viewing Ms_genome.fasta in RStudio using Biostrings package.

**Figure 31.**
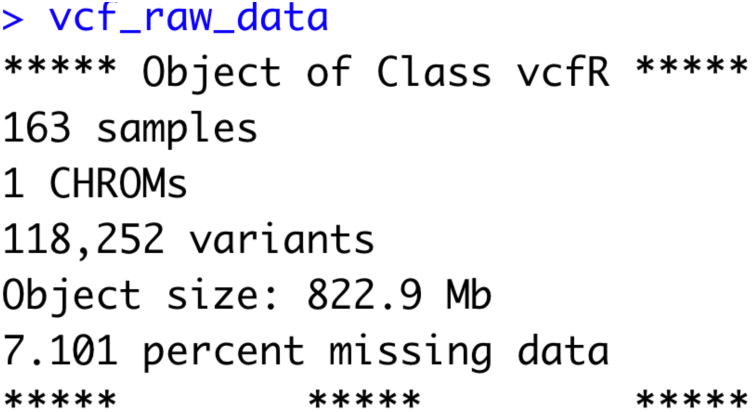
Statistics of Chr1.vcf.filtered.500M.vcf shown by vcfR.

### 7.3 Faster Access through Command-line-based Database Frontend

After we deployed the complete Alfalfa Database web application to our CentOS server for user testing, we noticed that the HTML rendering time was still a problem. For some users that prefer access speed to the genome or transcriptome data over good visualizations, waiting for the website to construct hyperlinks and format the tables could be unnecessary. Thus, a supplementary command-line-based Java frontend application has been developed for faster access to our database file, as shown in Section 10.8. Note that for VCF Queries in this application, the POS range was not restricted, so that users could work with the VCF data more productively. Some sample queries were recorded in this video: https://youtu.be/qs05TDjWvd0. This Java application was mainly based on a JDBC package, sqlite-jdbc-3.36.0.1.jar, and the SQL query statements were very similar to what was utilized in the web application. Those queries were executed via the stat.executeQuery(<SQL SELECT statements>) method on Line 95, 111, 141, 155, 196, and 272. After the query results were returned to a ResultSet object called rs, rs.next() would return true when there were more rows of unprocessed results. In this case, rs.getString(<COLUMN name>) was called to read out and print the proper searching result to the console window. To improve the readability of query results on the command line, variable cnt was used repeatedly throughout different queries to count how many meaningful rows of data had been returned. Many other formatting details and strategies to eliminate NULL results were also applied to improve user experience.

## 8. Availability

The Alfalfa Database Project, including the Python, Java, and SQL scripts used during the development stage, the Flask web application, and the Java command line application are all open-sourced and posted on our GitHub repository:

https://github.com/ShichenQiao/Alfalfa_Database.git

However, up to the point we post this paper, we are not authorized to reveal the original dataset, our intermediate datafiles, and the resulting SQLite backend database to the public.

The service of our web application is only available at the Local Area Network (LAN) of China Agriculture University at 202.112.170.236:5000.

## 9. Authors’ Contributions

SQ designed the backend SQLite database and the frontend applications, developed the Python, Java, and SQL scripts used during the development stage (except vcf.java as shown in Section 10.2), and wrote the frontend web application and the command line application. CS managed and filtered datafiles, wrote vcf.java, provided server support, processed the complete VCF files, and deployed our web application to the LAN. All of the authors have read and approved the final form of this paper.

## 10. Appendix: Complete Scripts

### 10.1 create_tables.sql

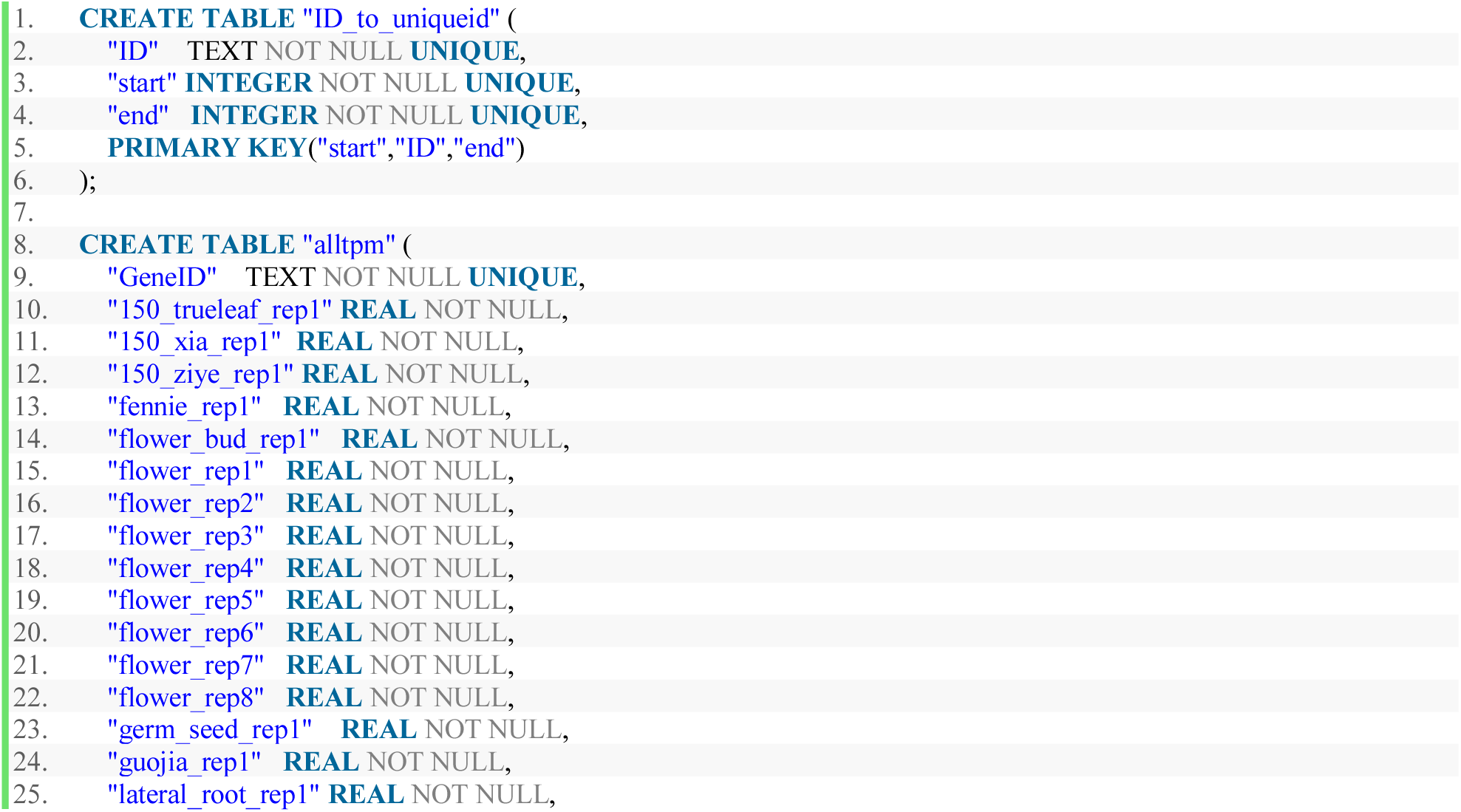
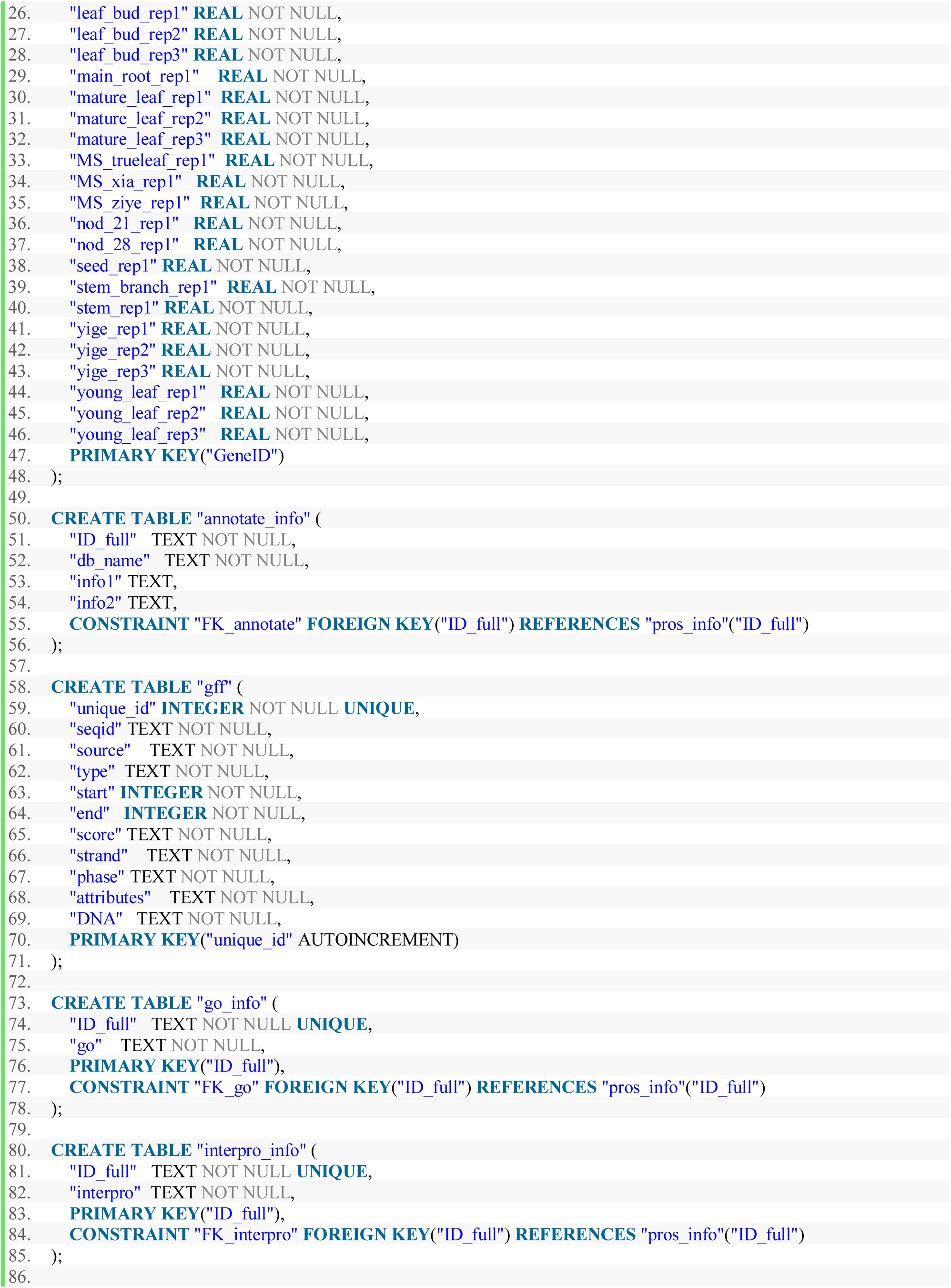
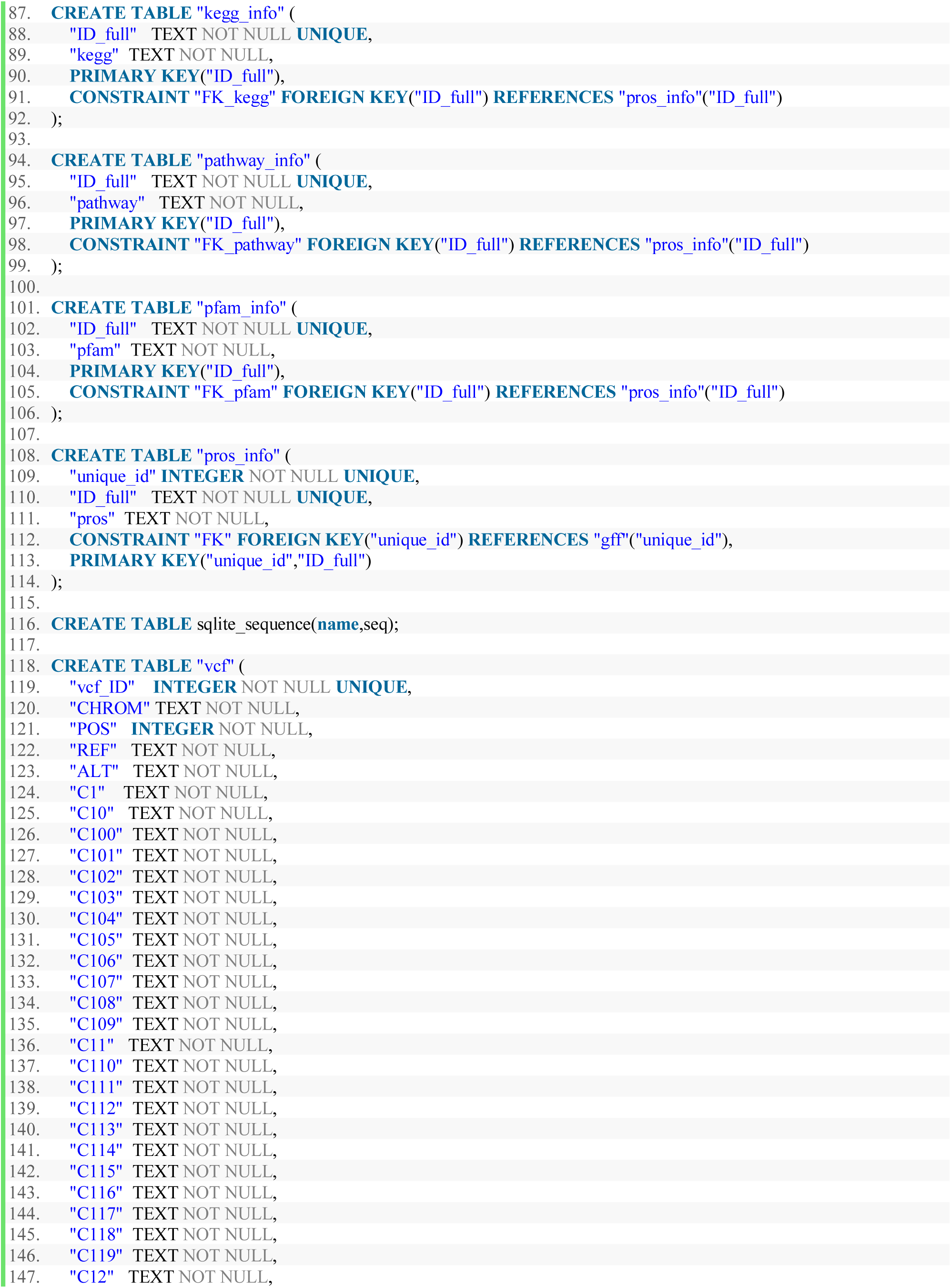
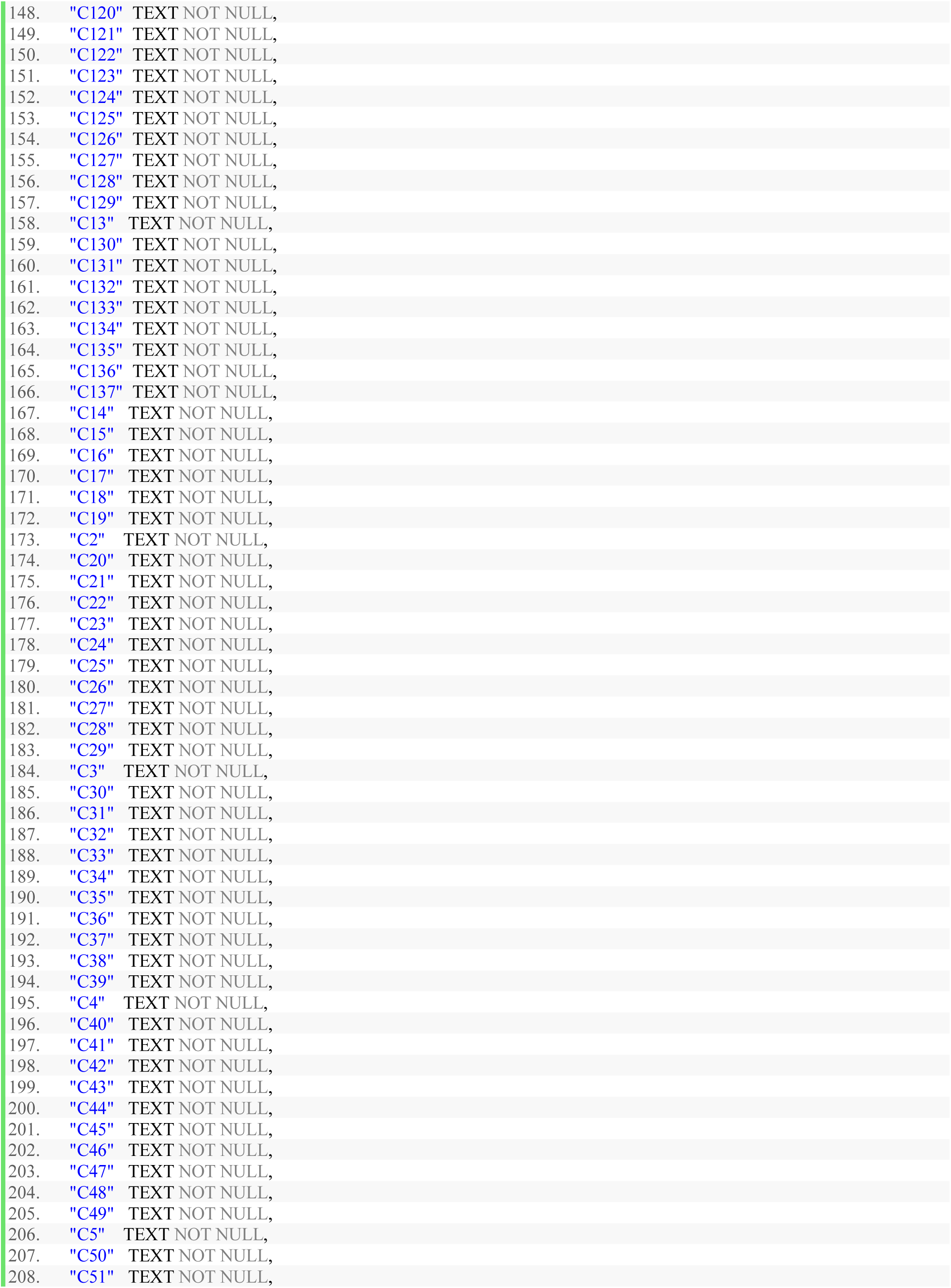
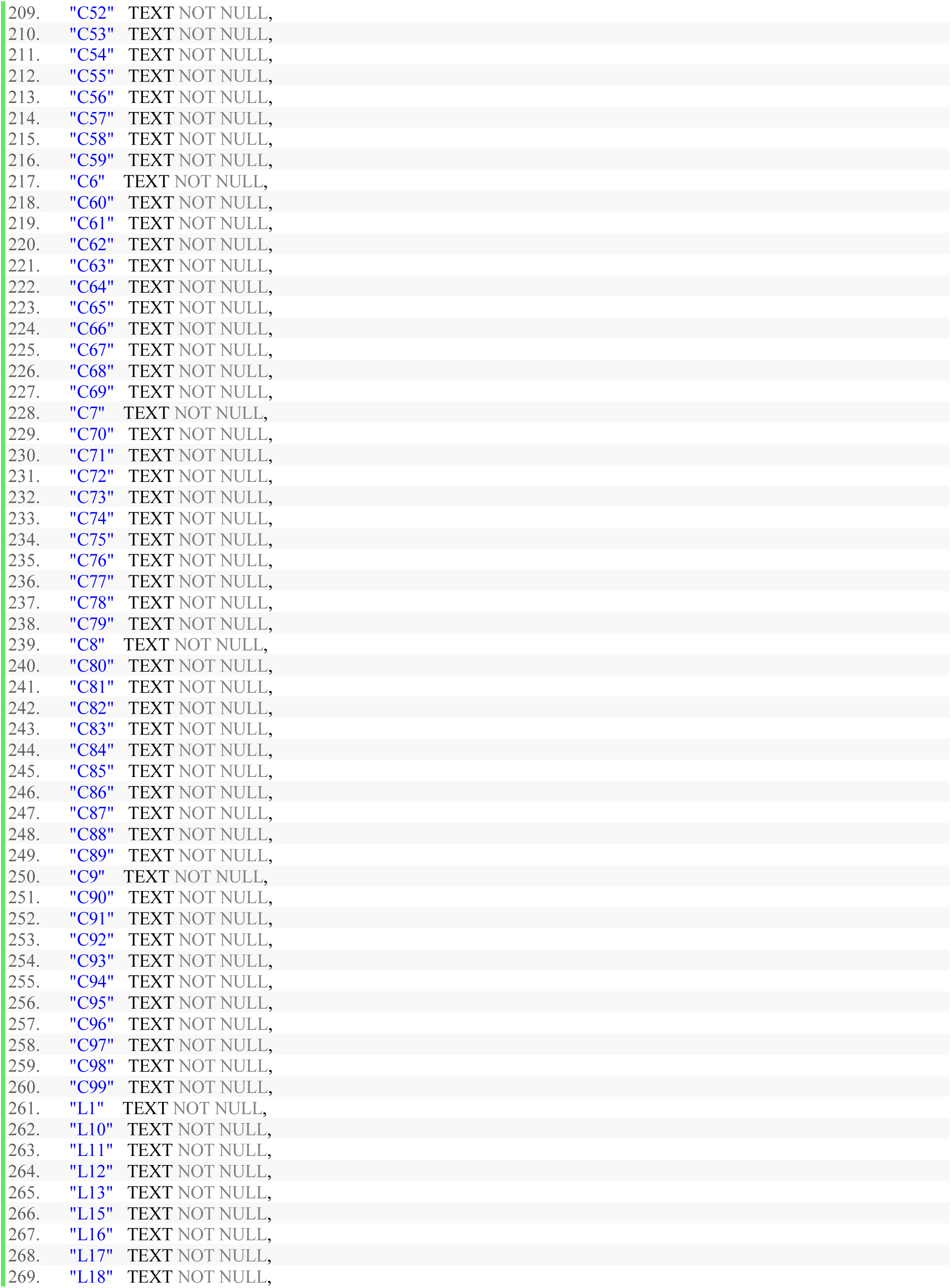
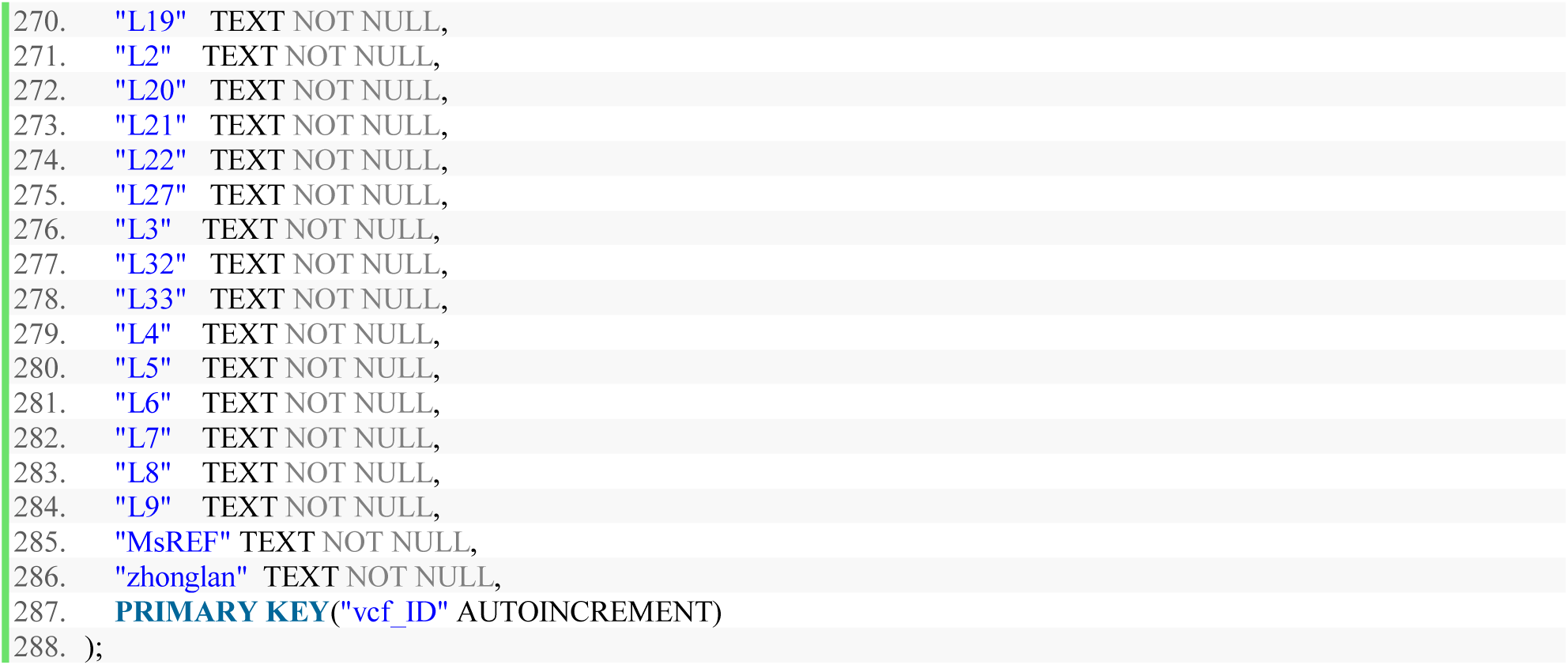

### 10.2 vcf.java

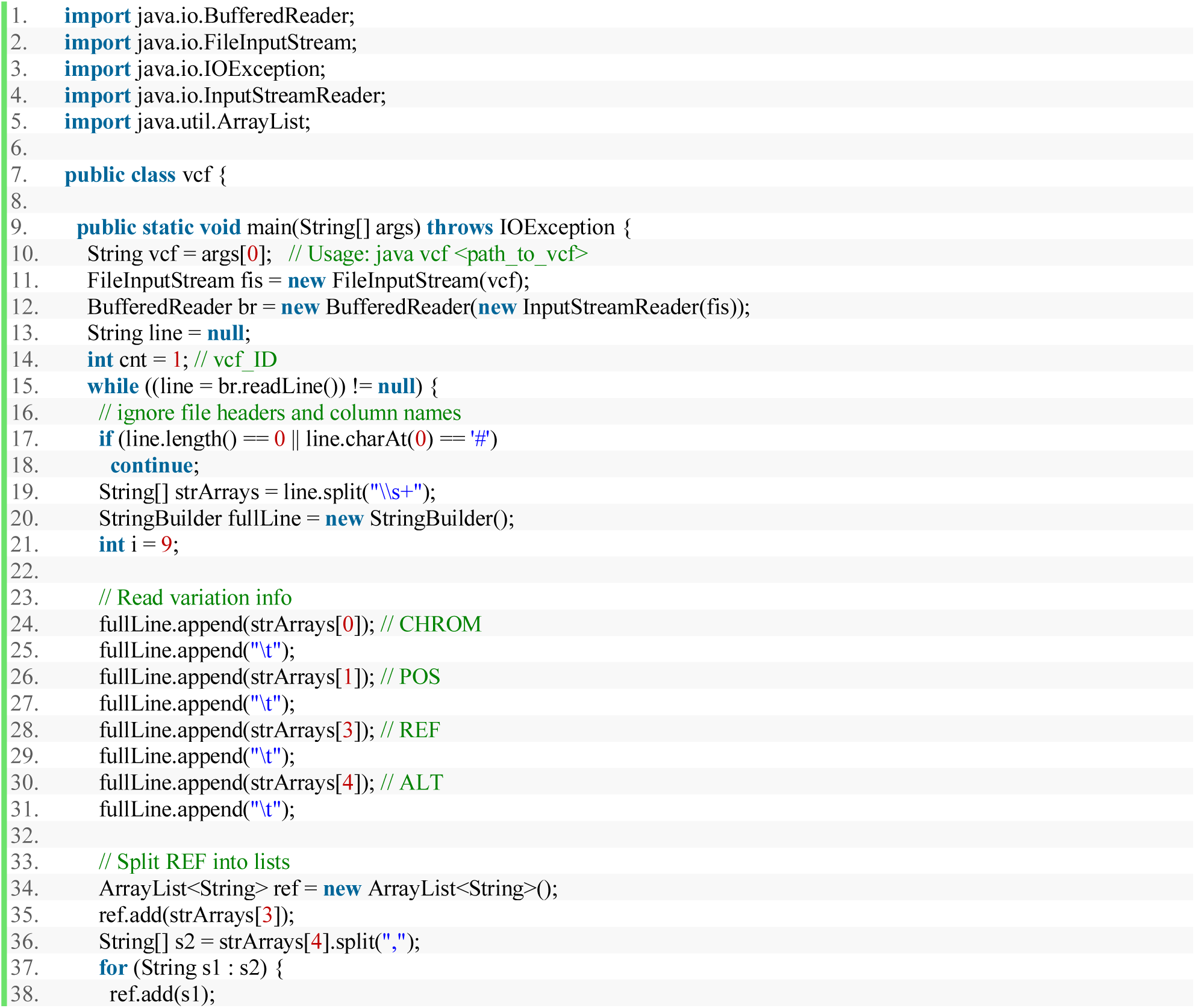
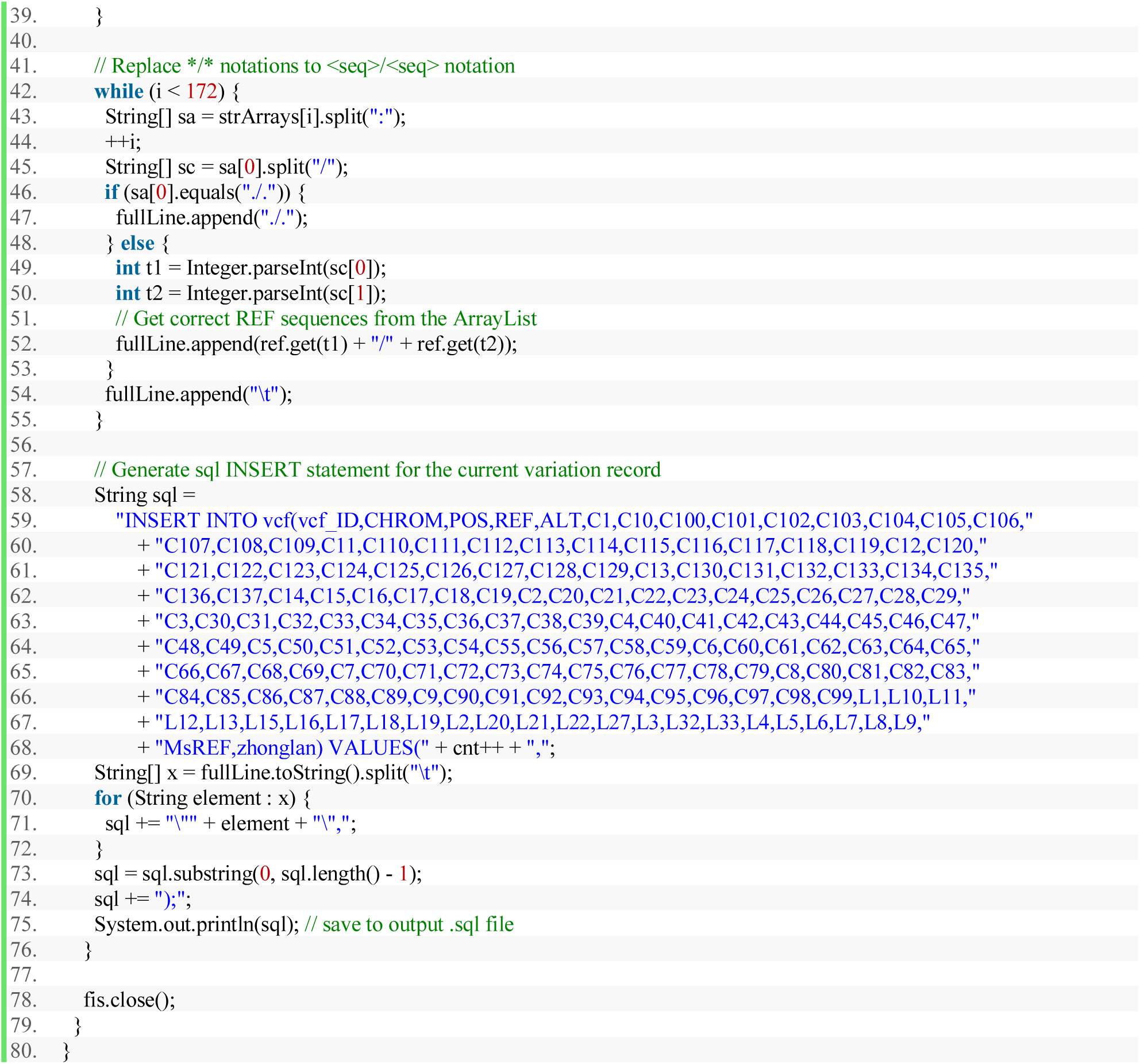

### 10.3 app.py

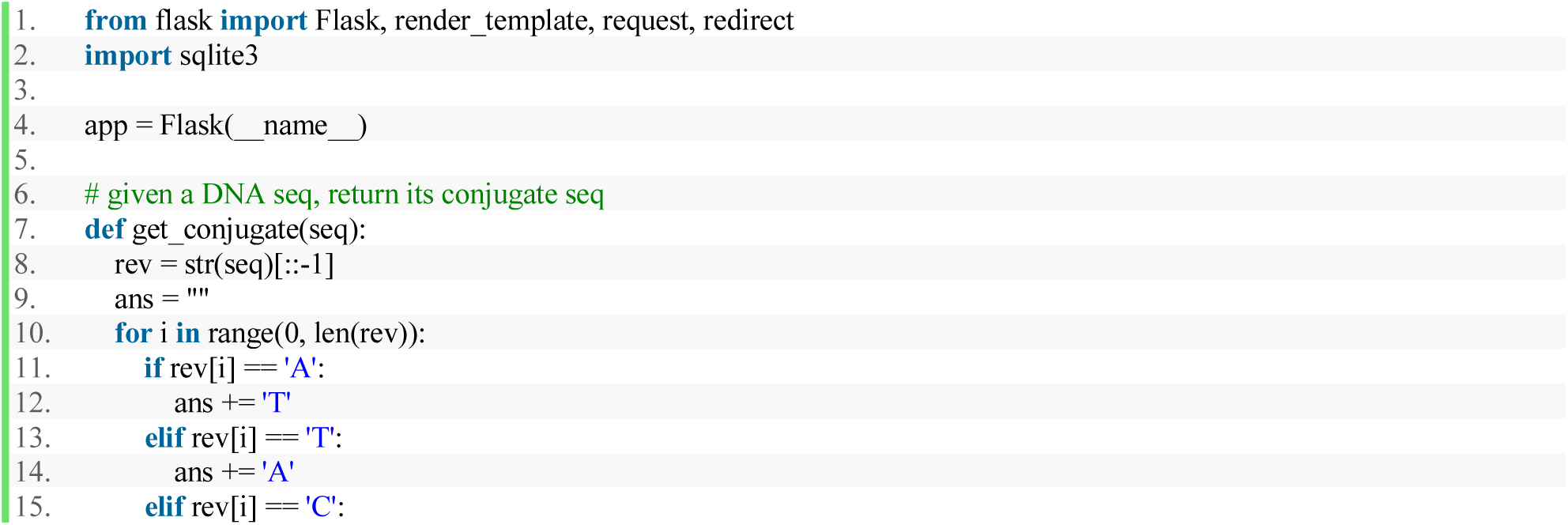
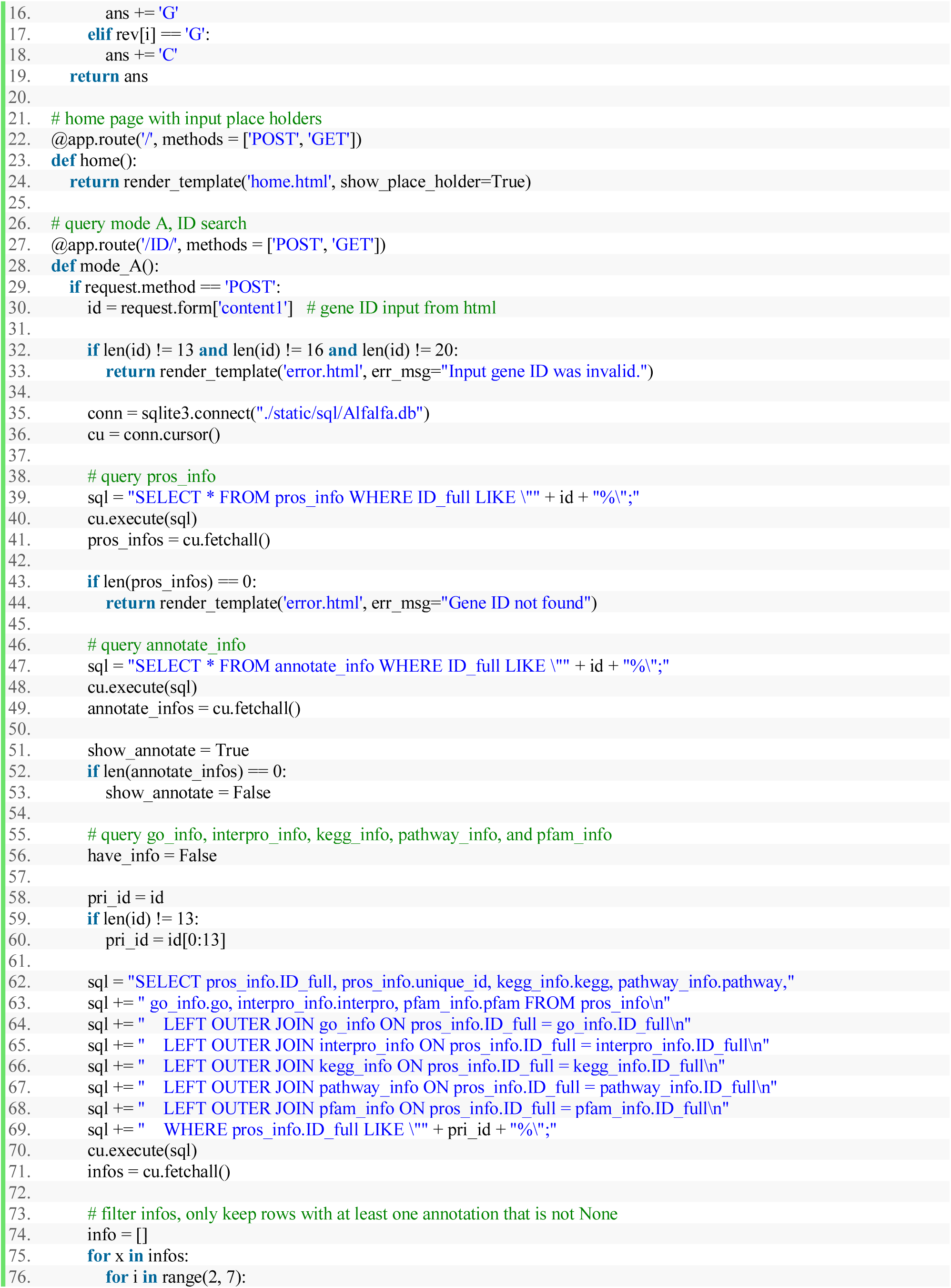
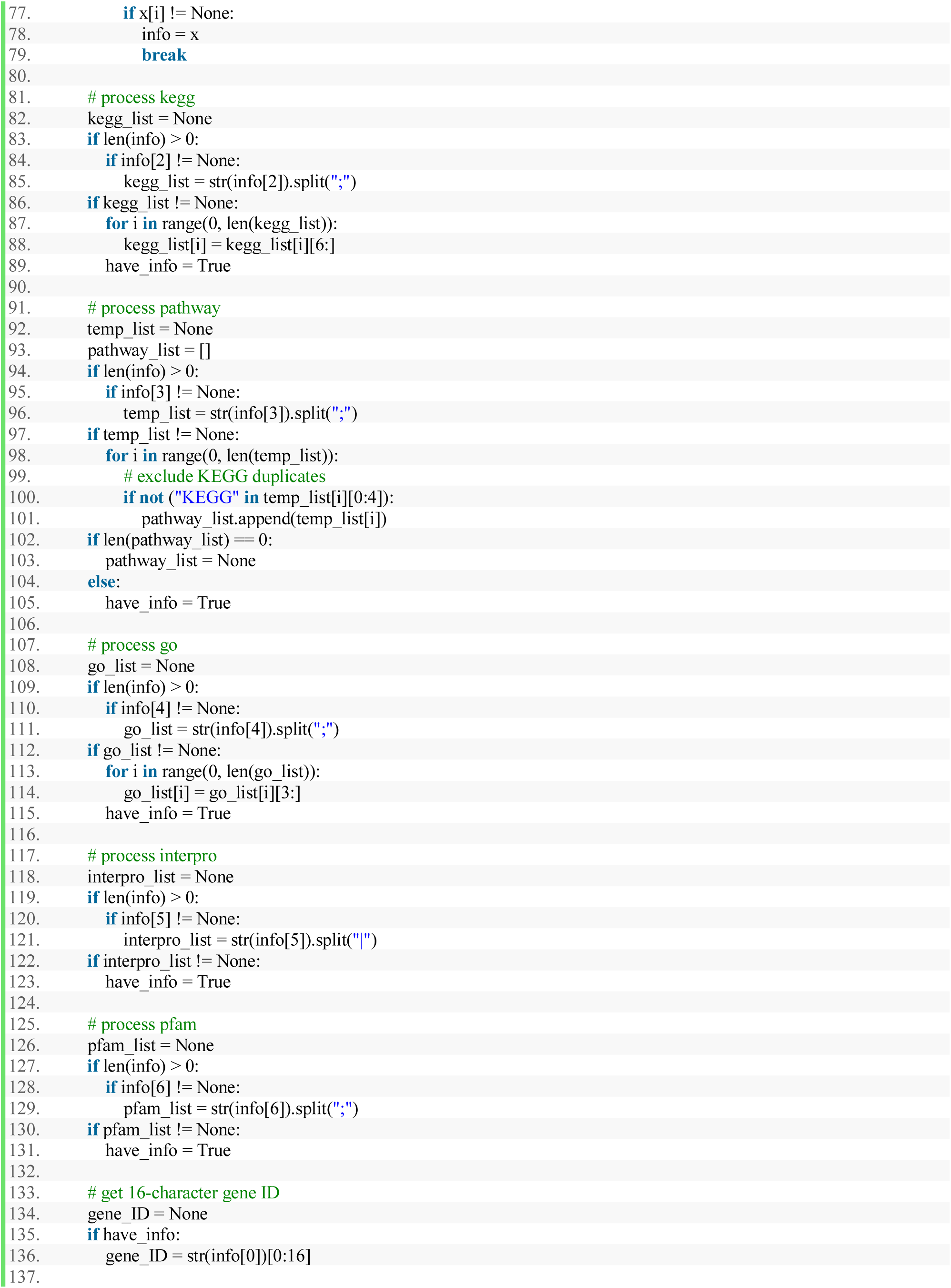
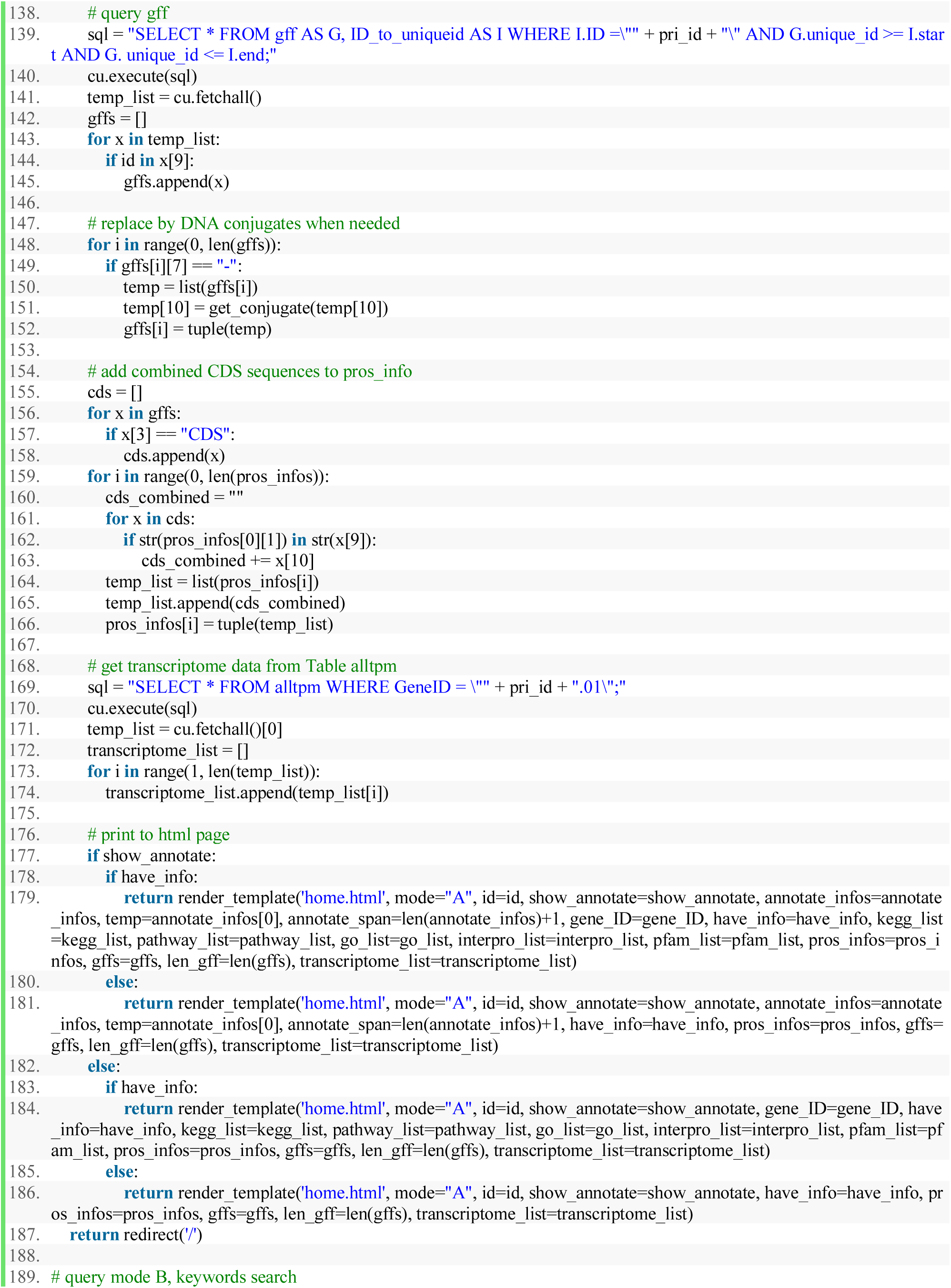
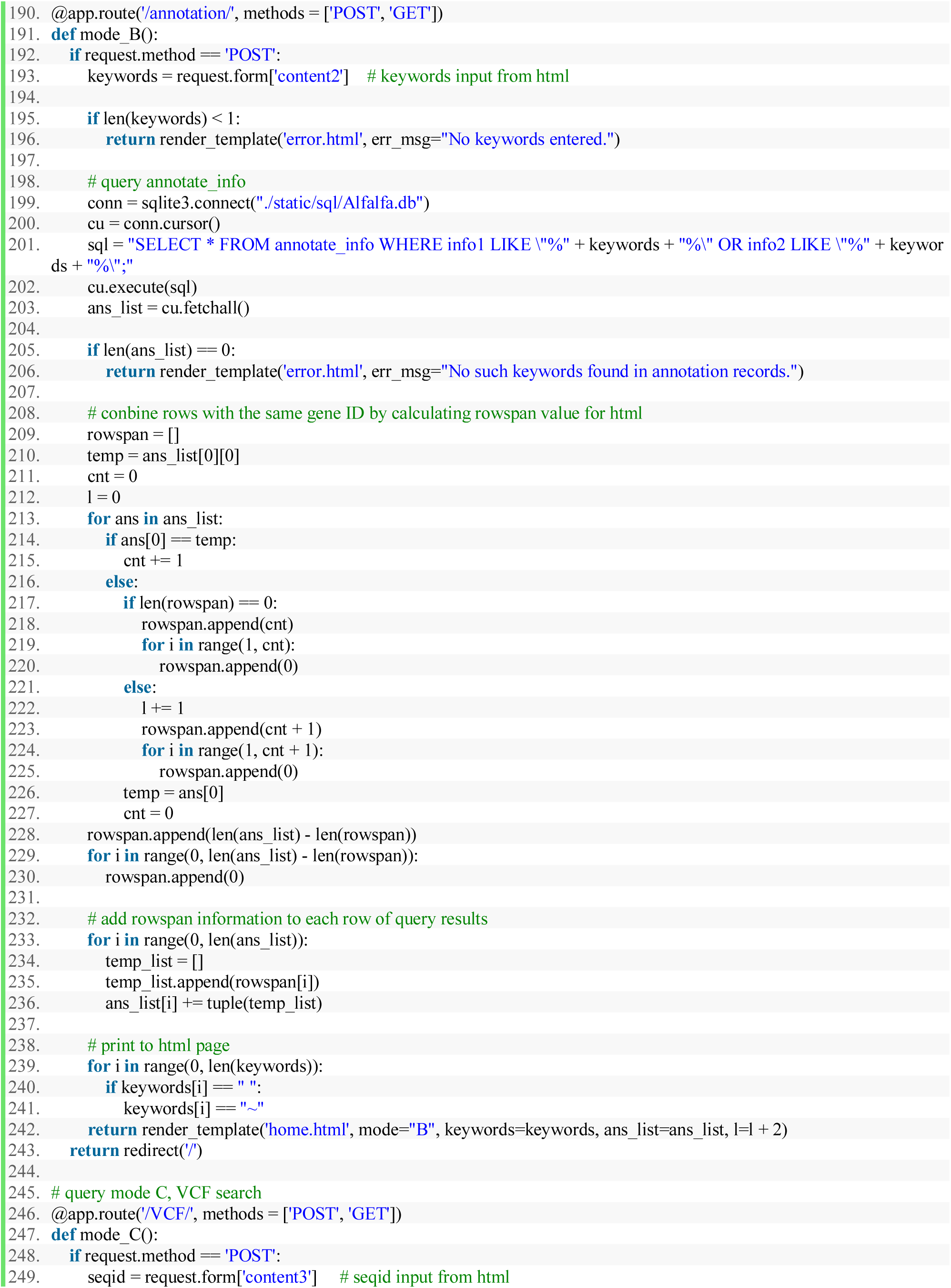
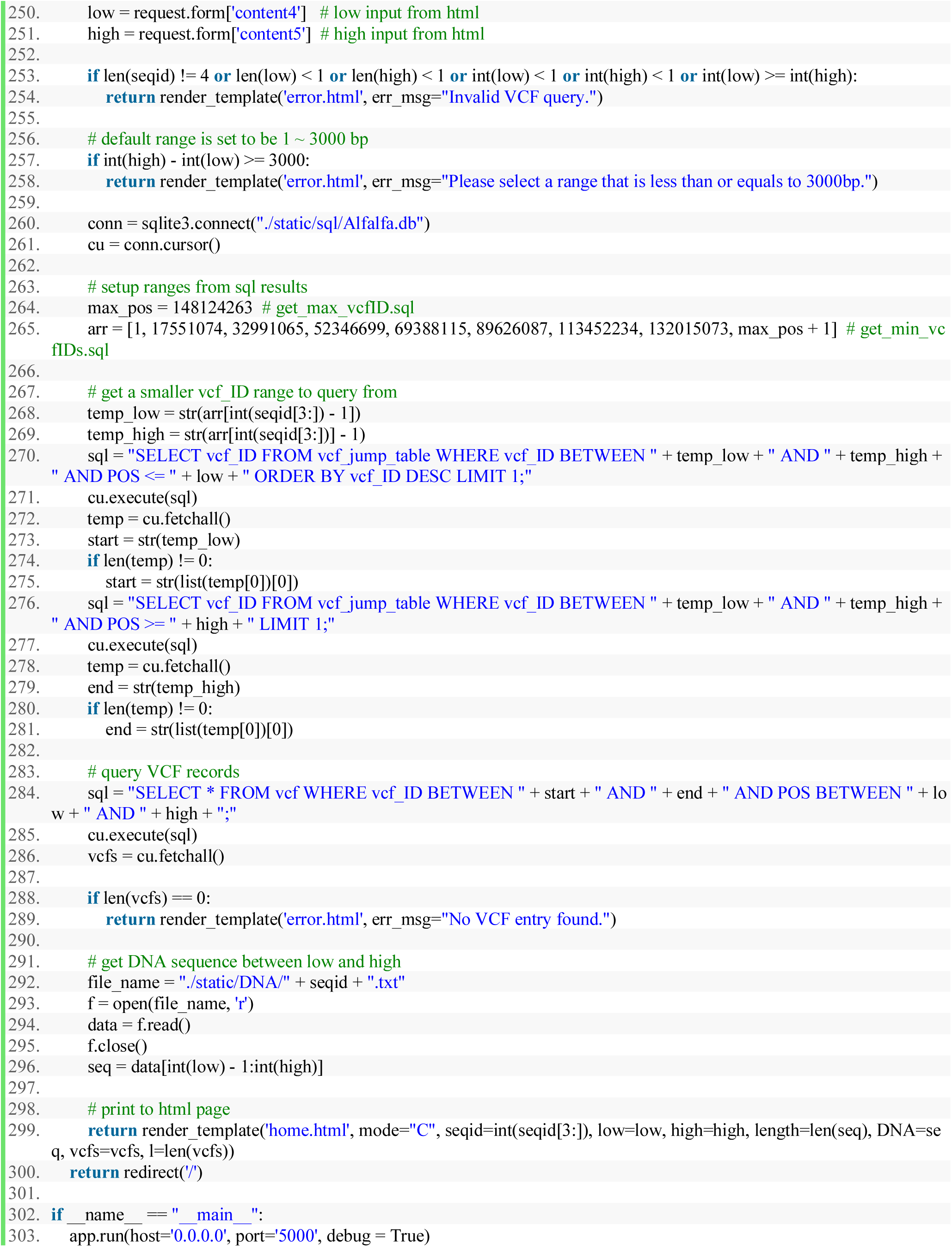

### 10.4 base.html

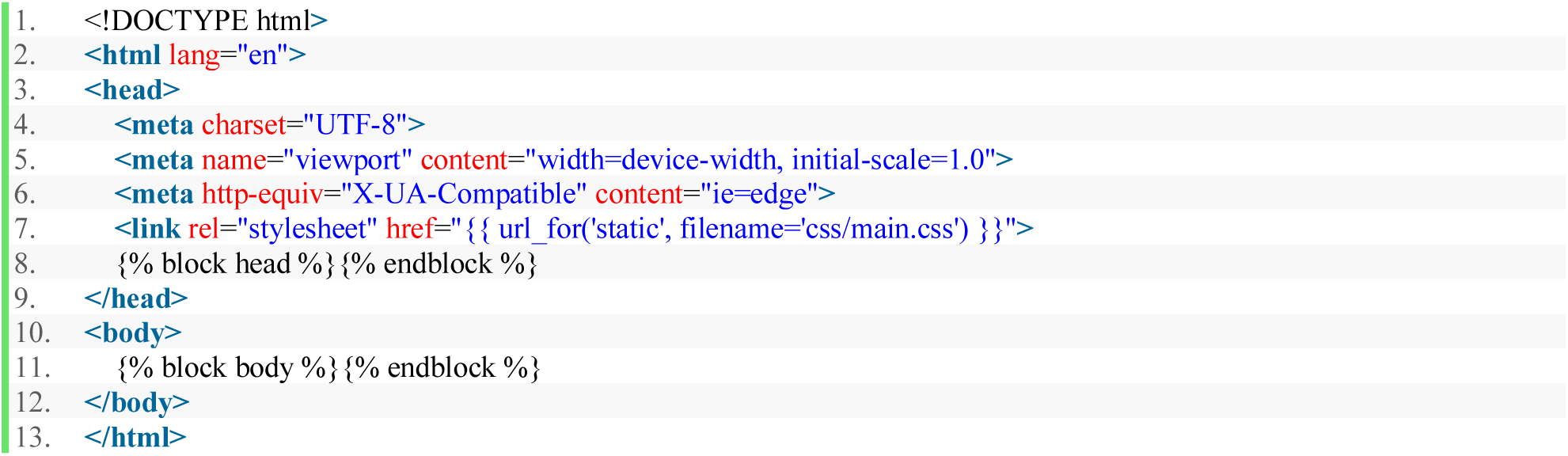

### 10.5 home.html

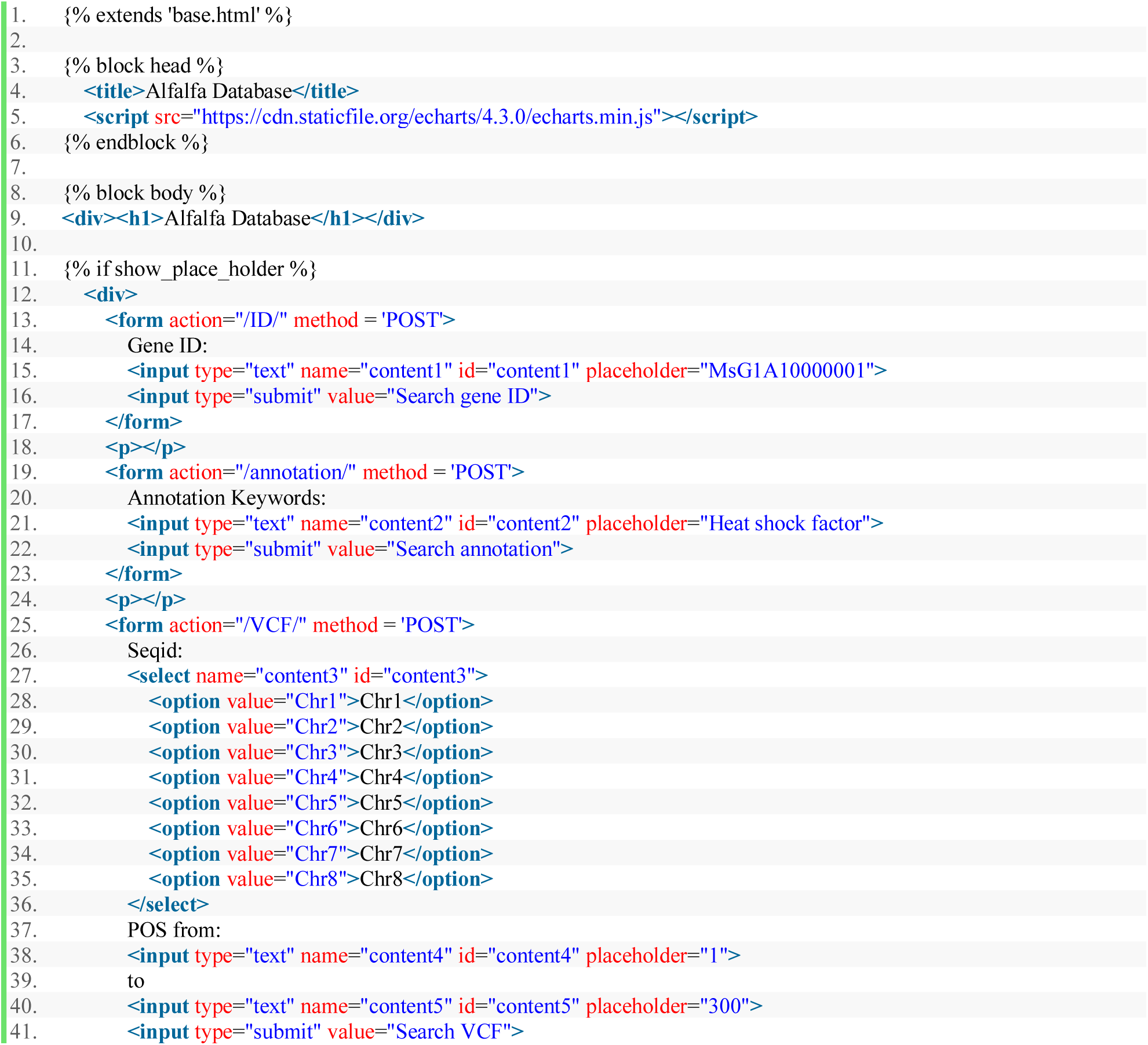
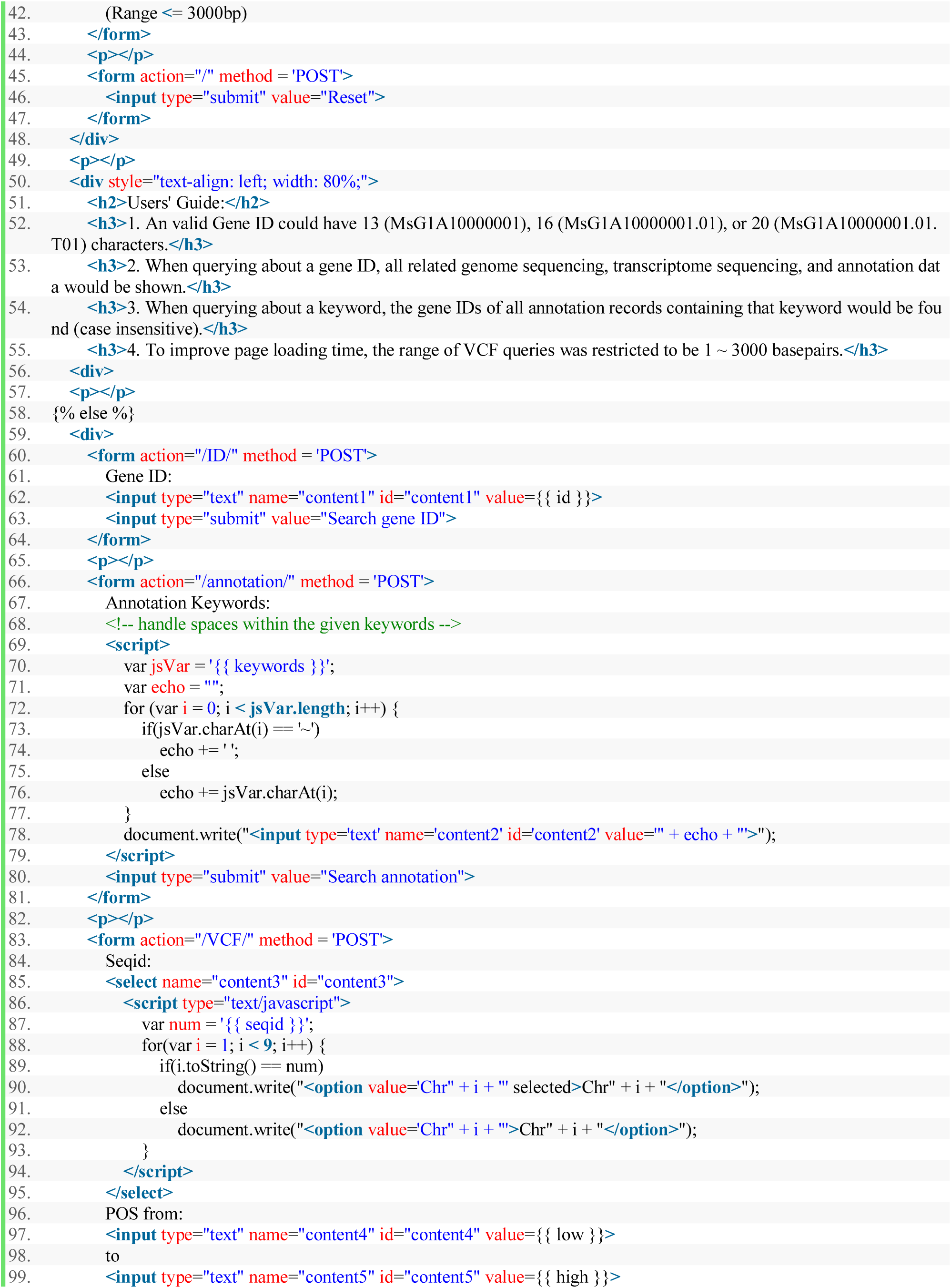
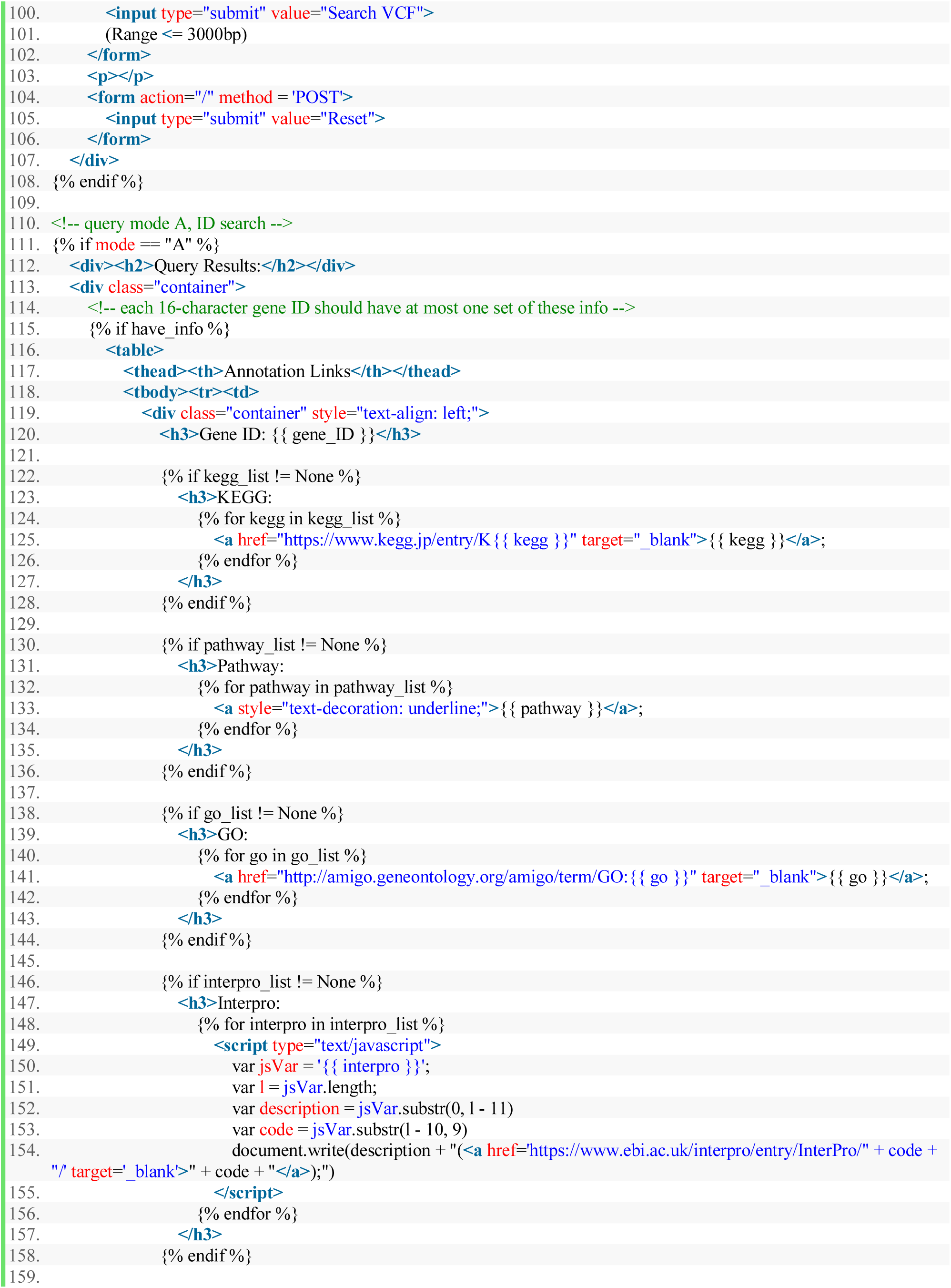
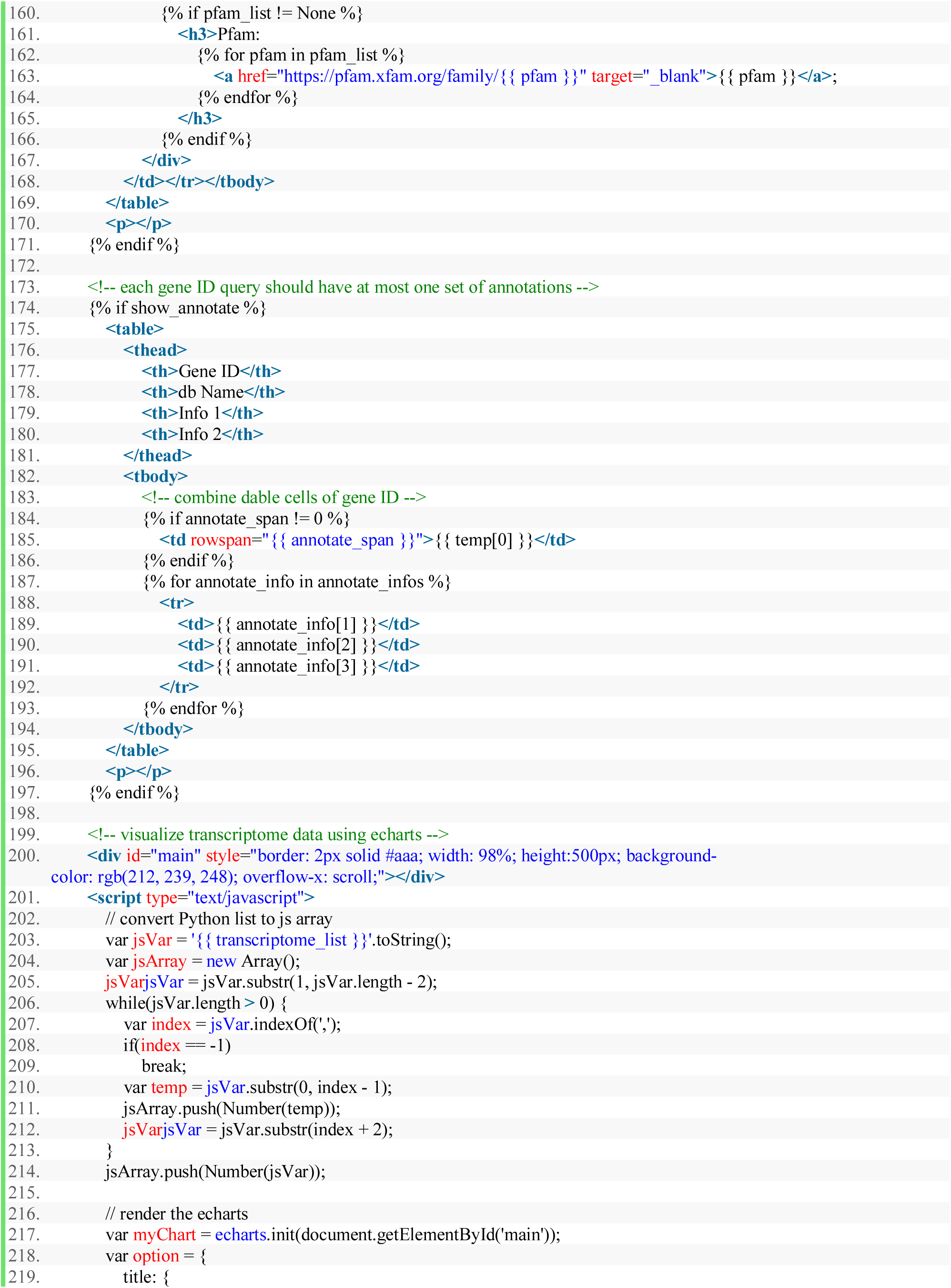
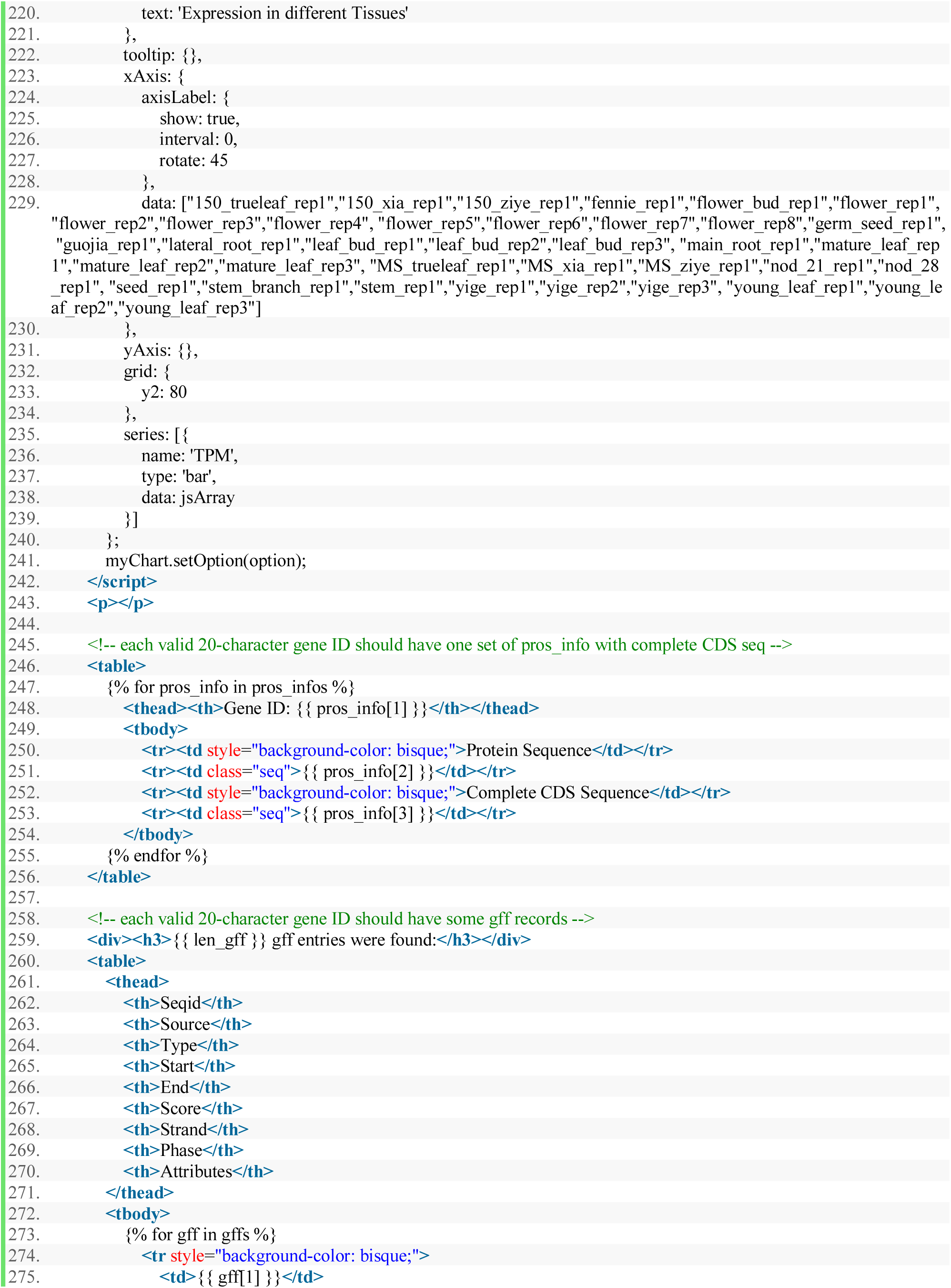
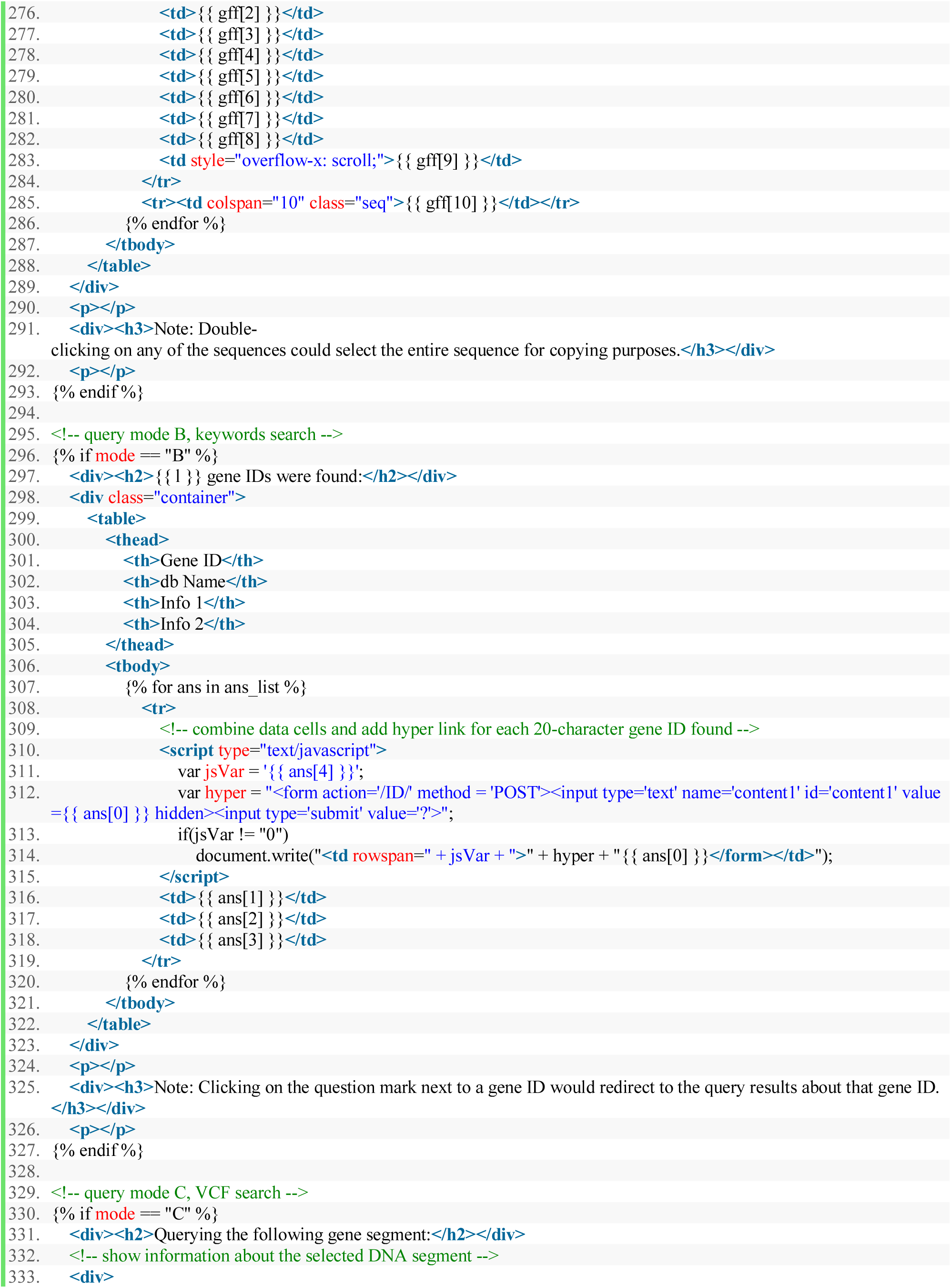
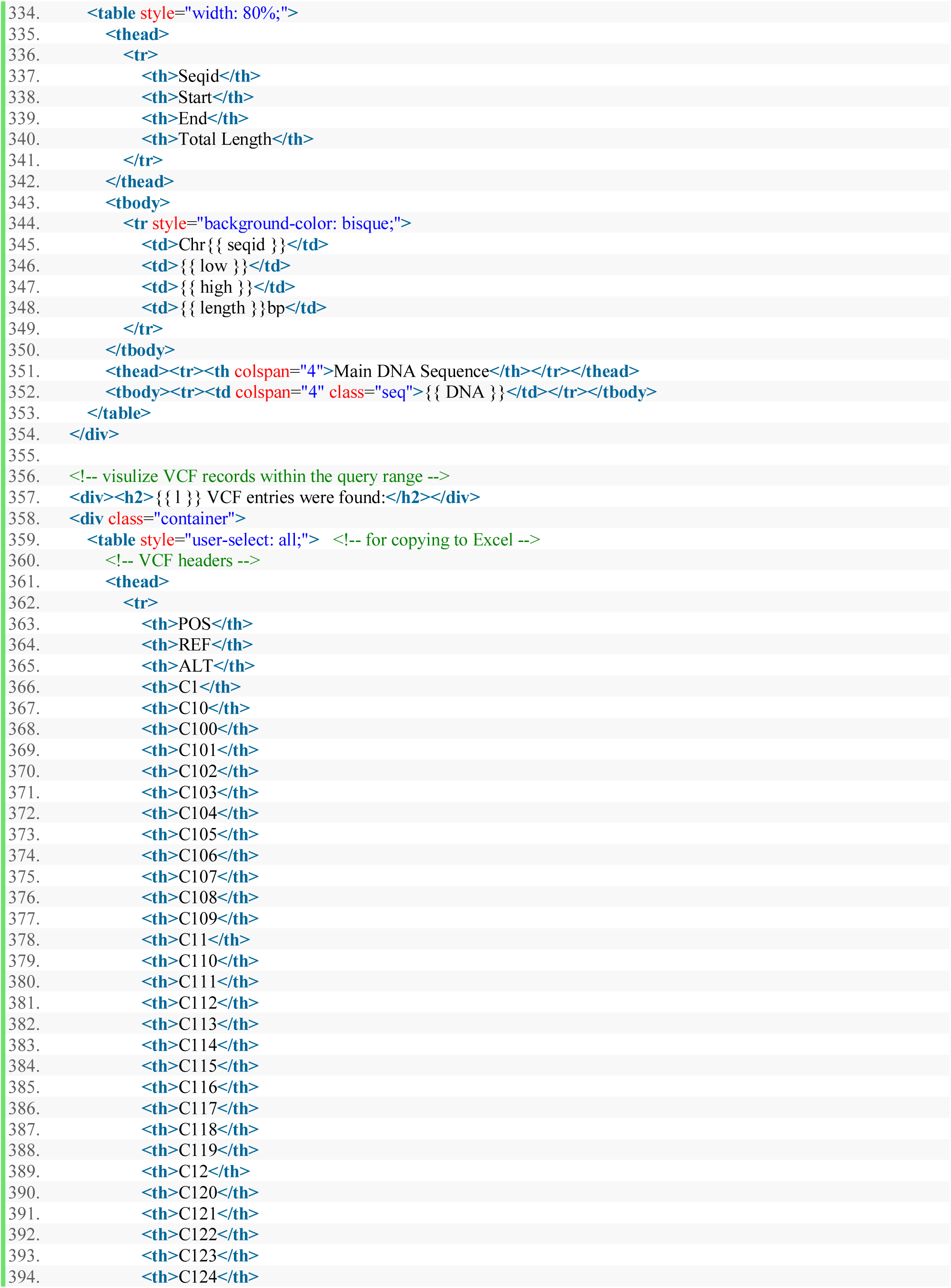
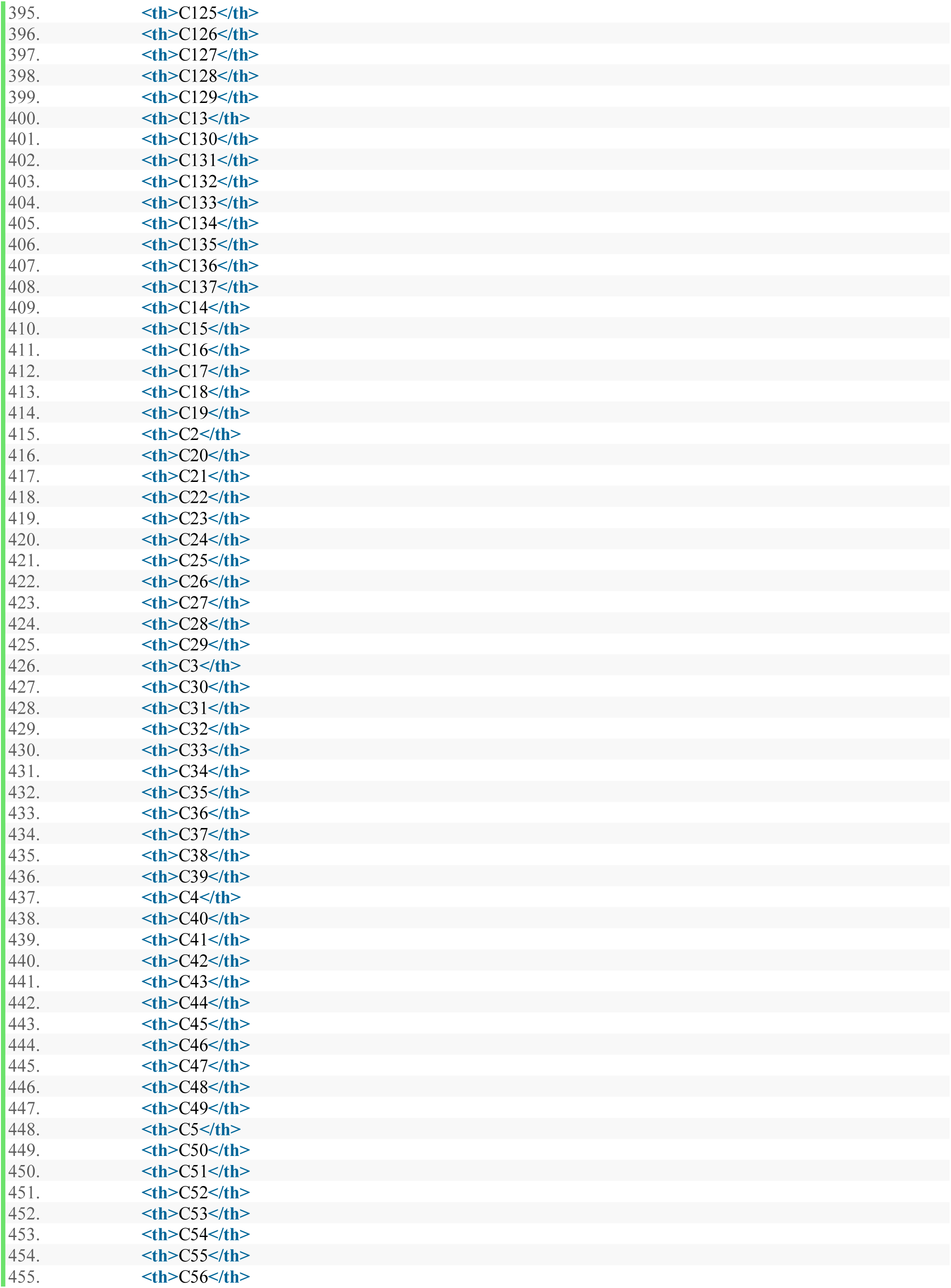
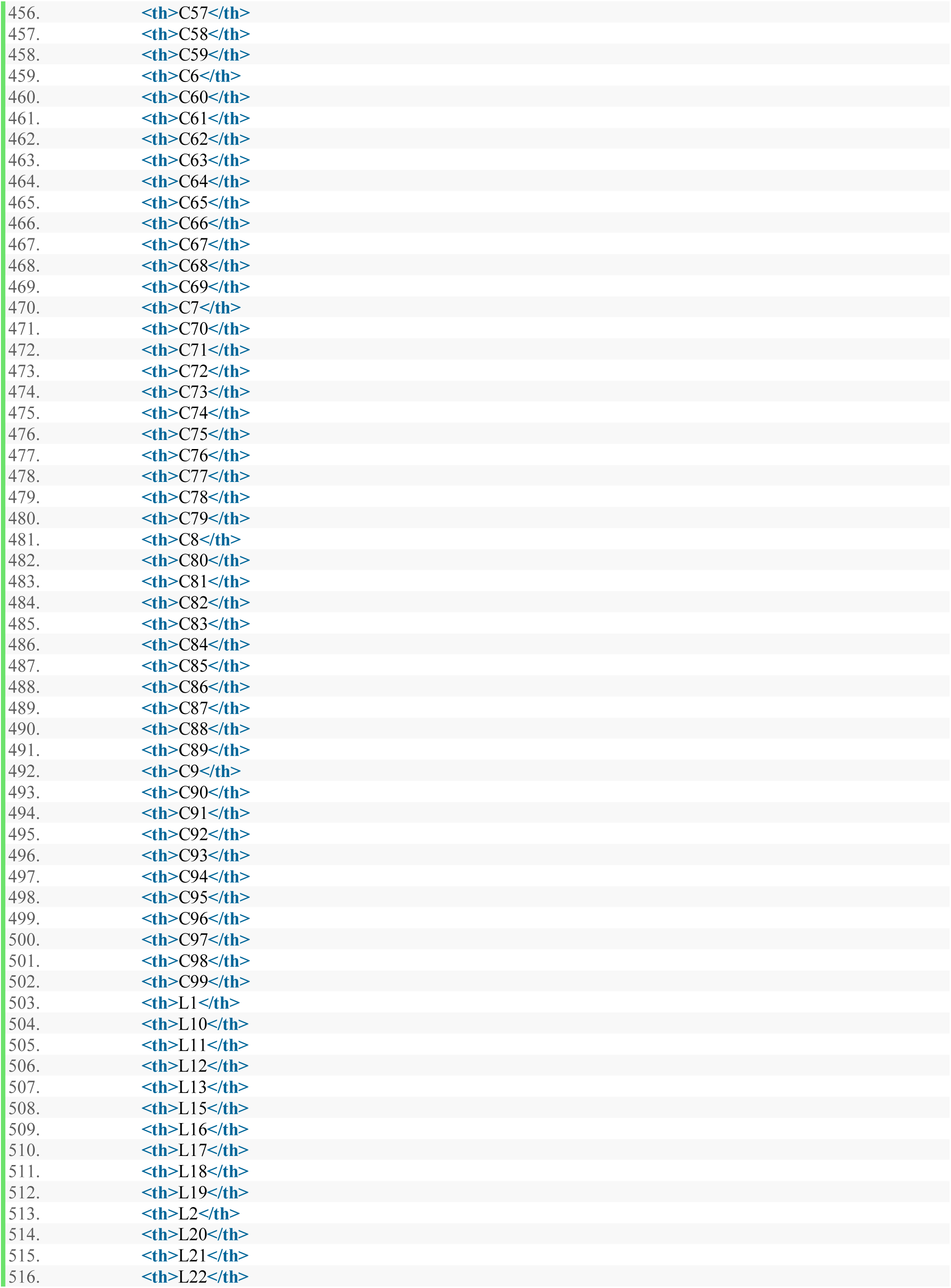
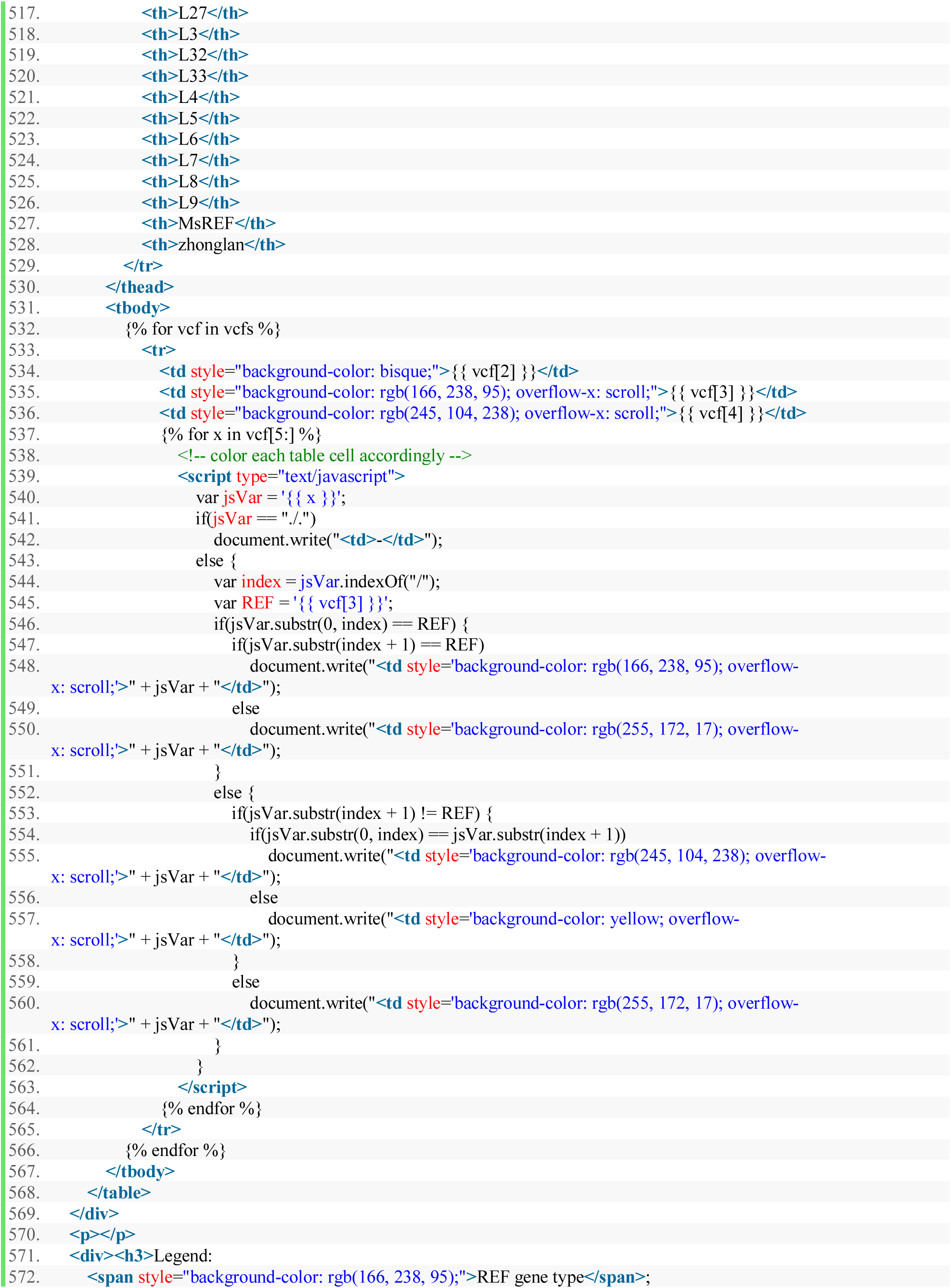
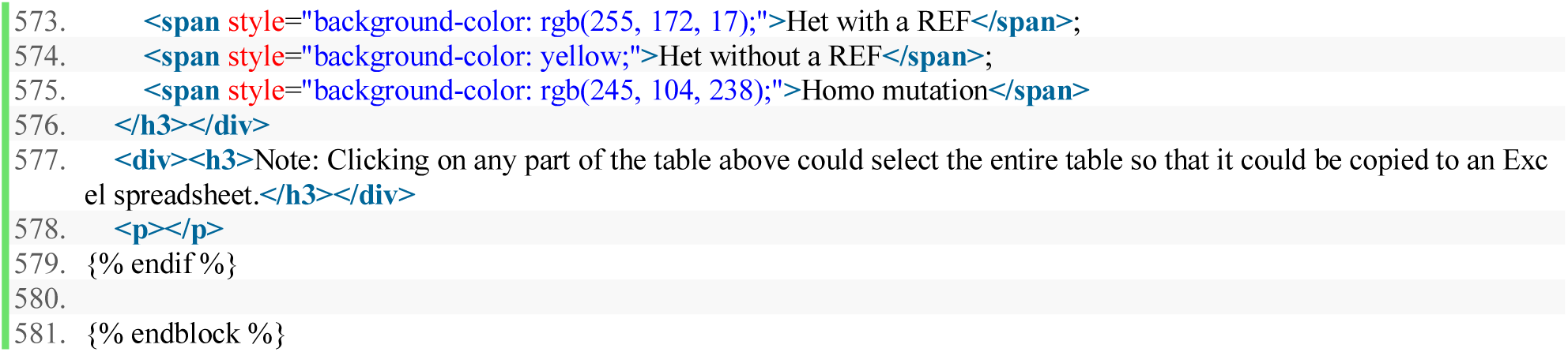

### 10.6 error.html

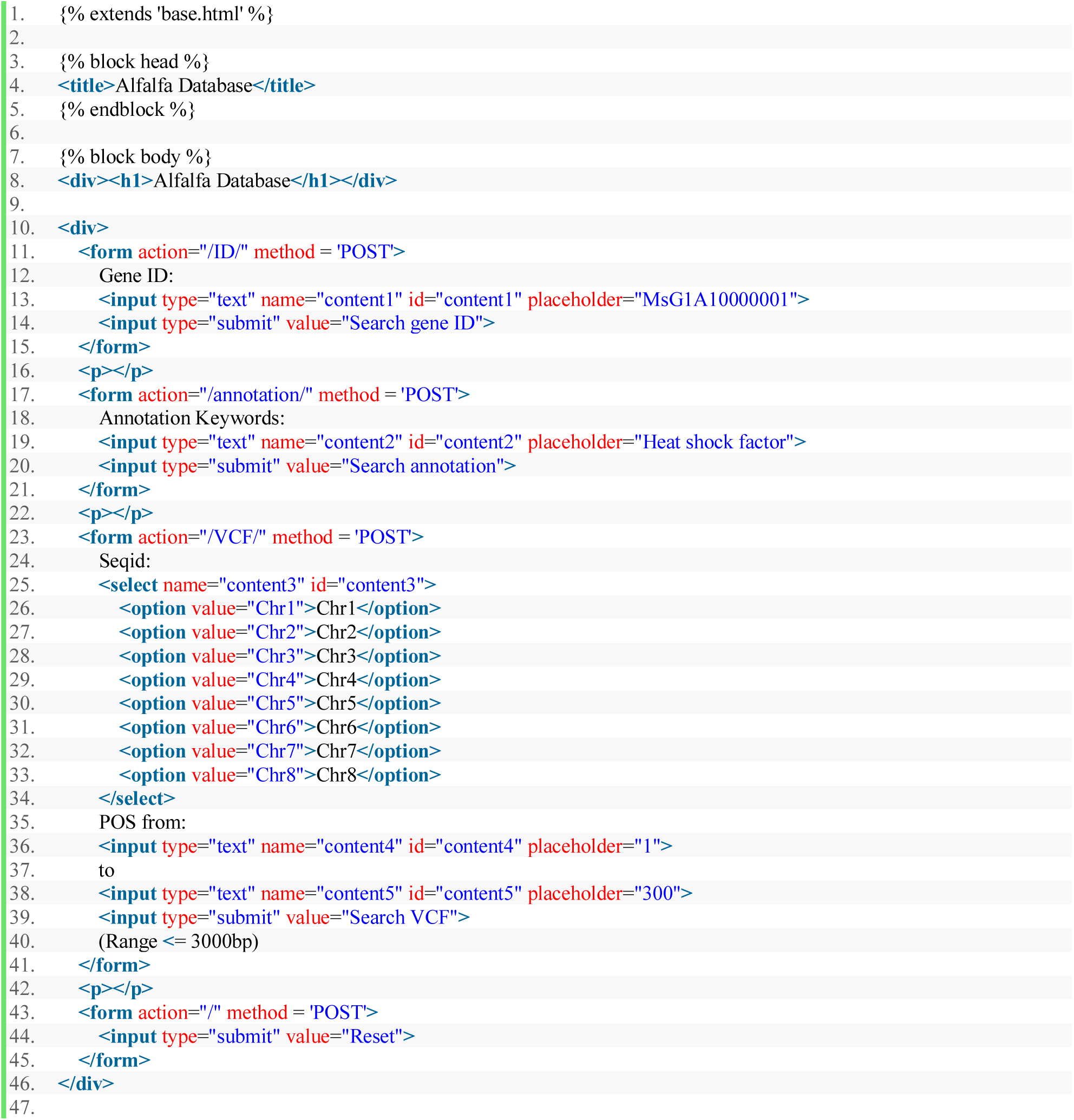
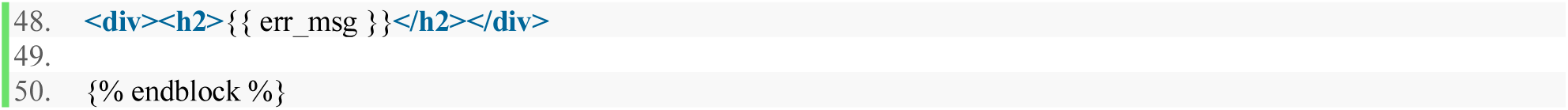

### 10.7 main.css

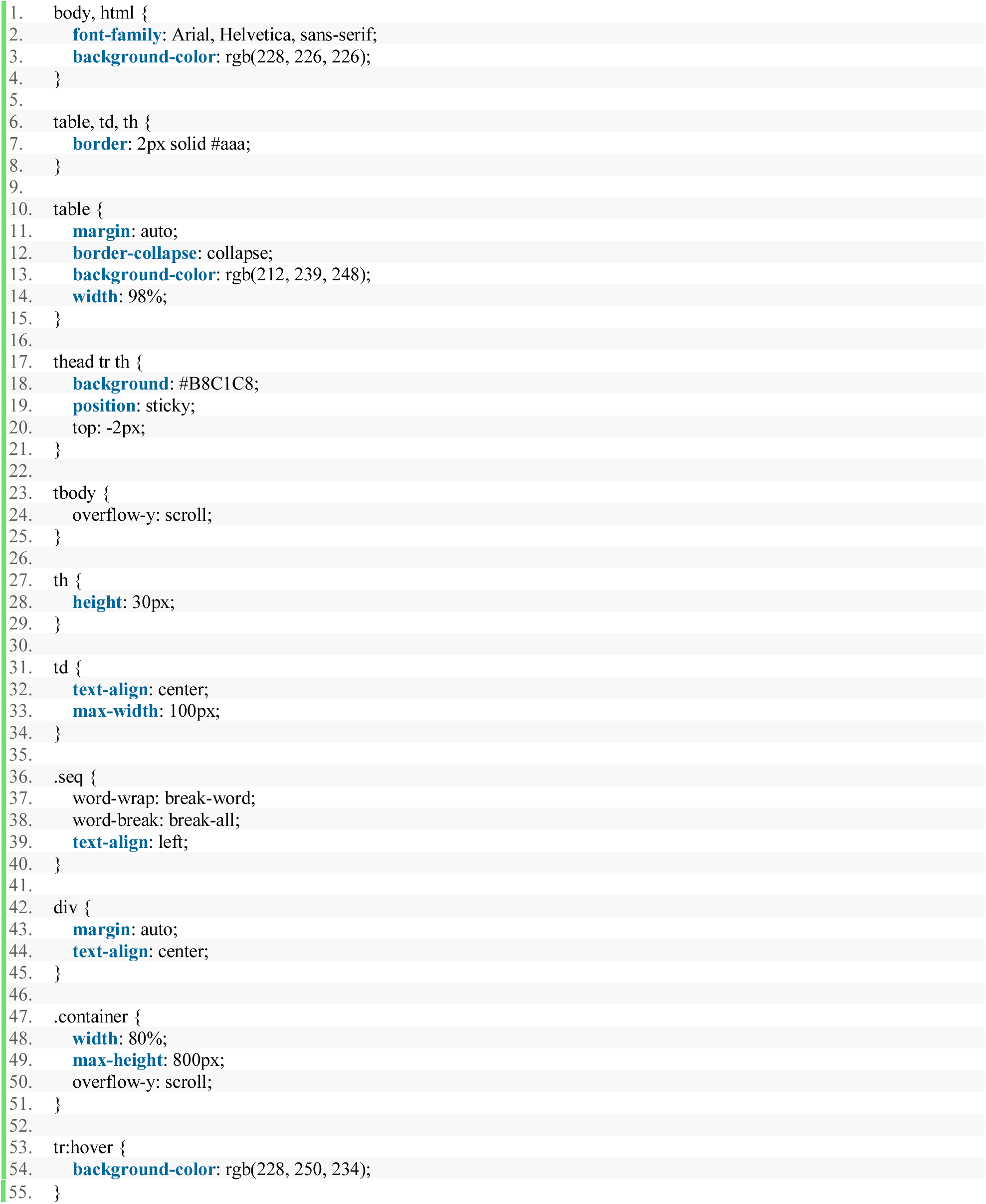

### 10.8 Alfalfa_Database.java

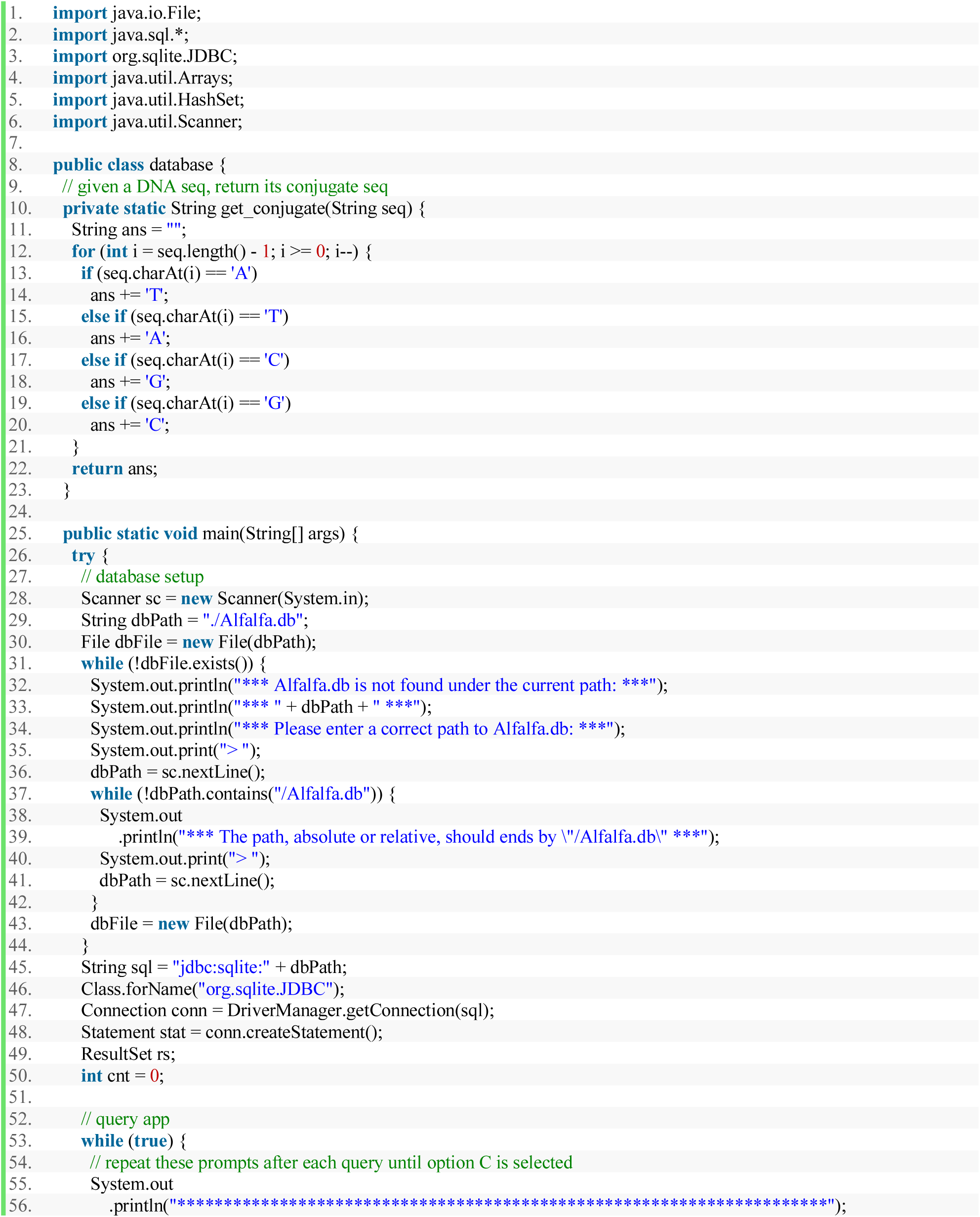
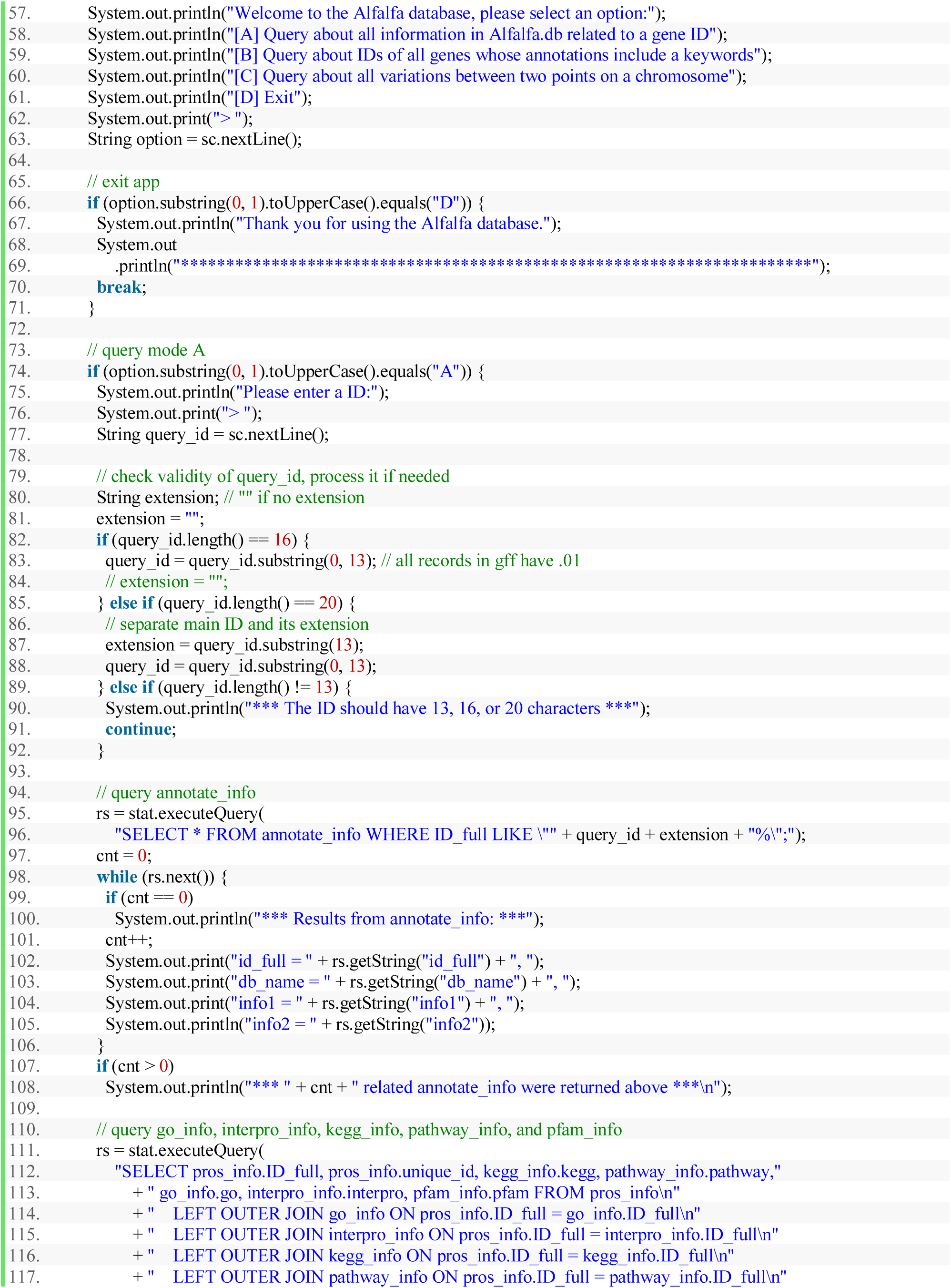
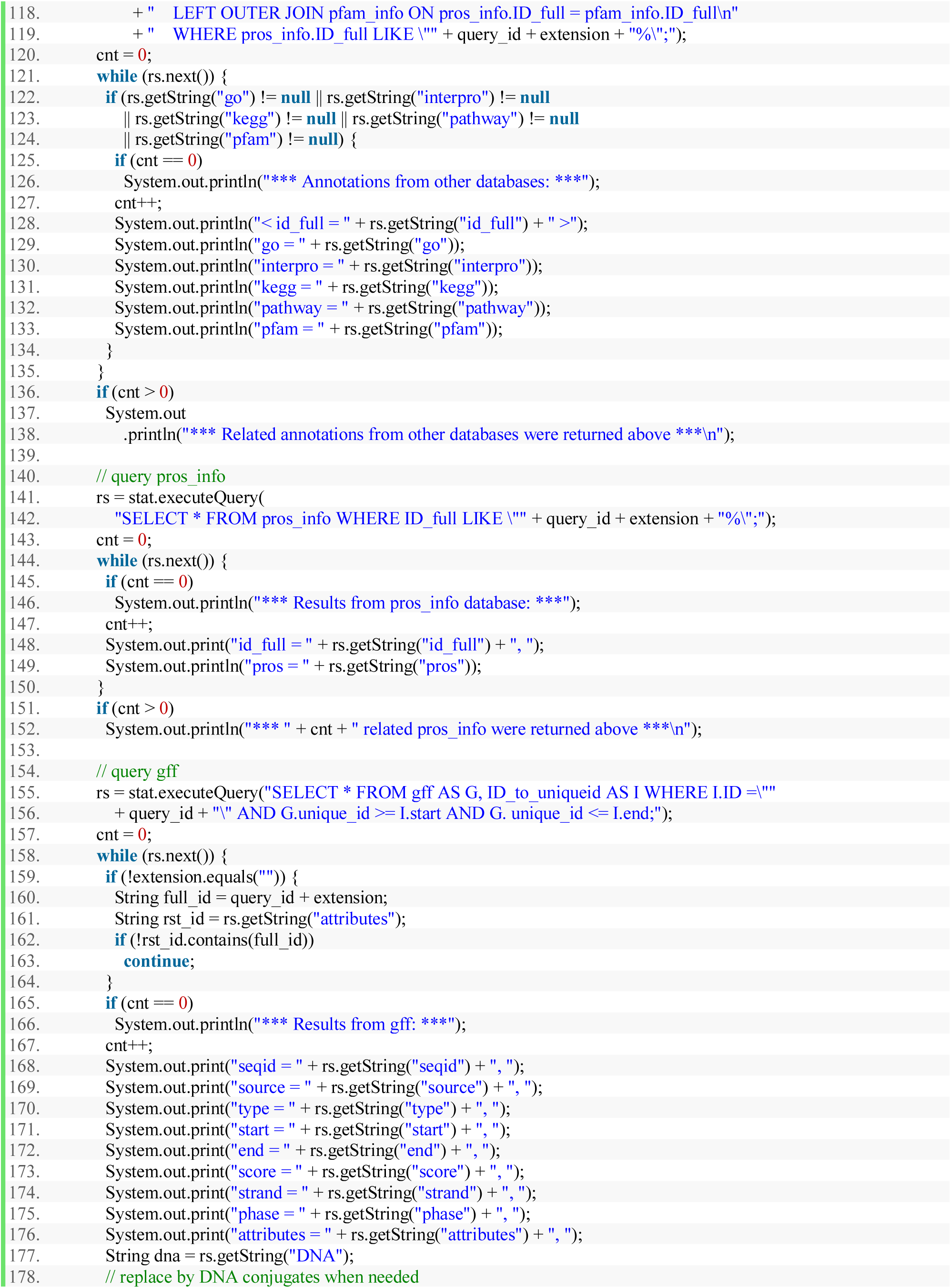
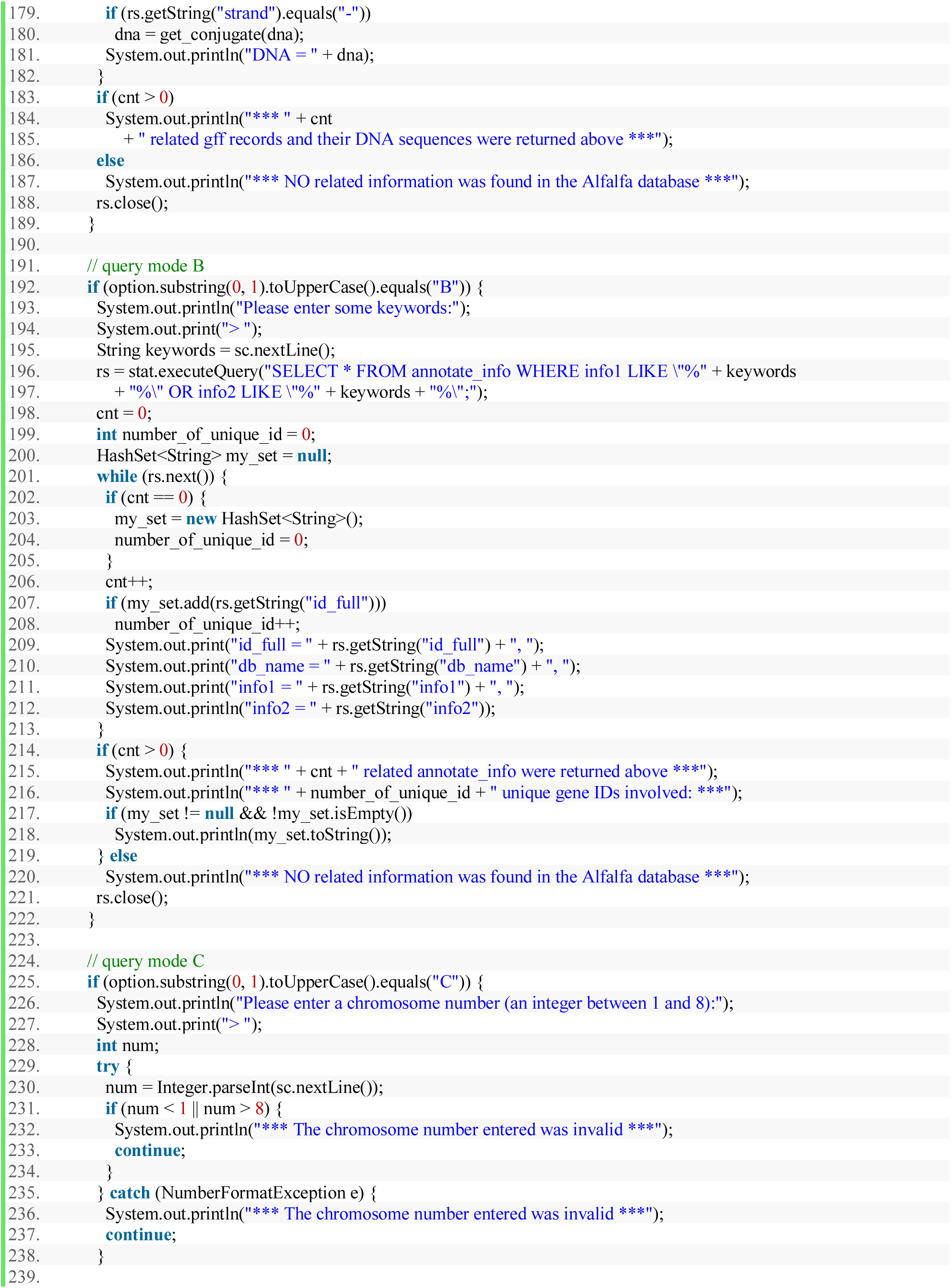
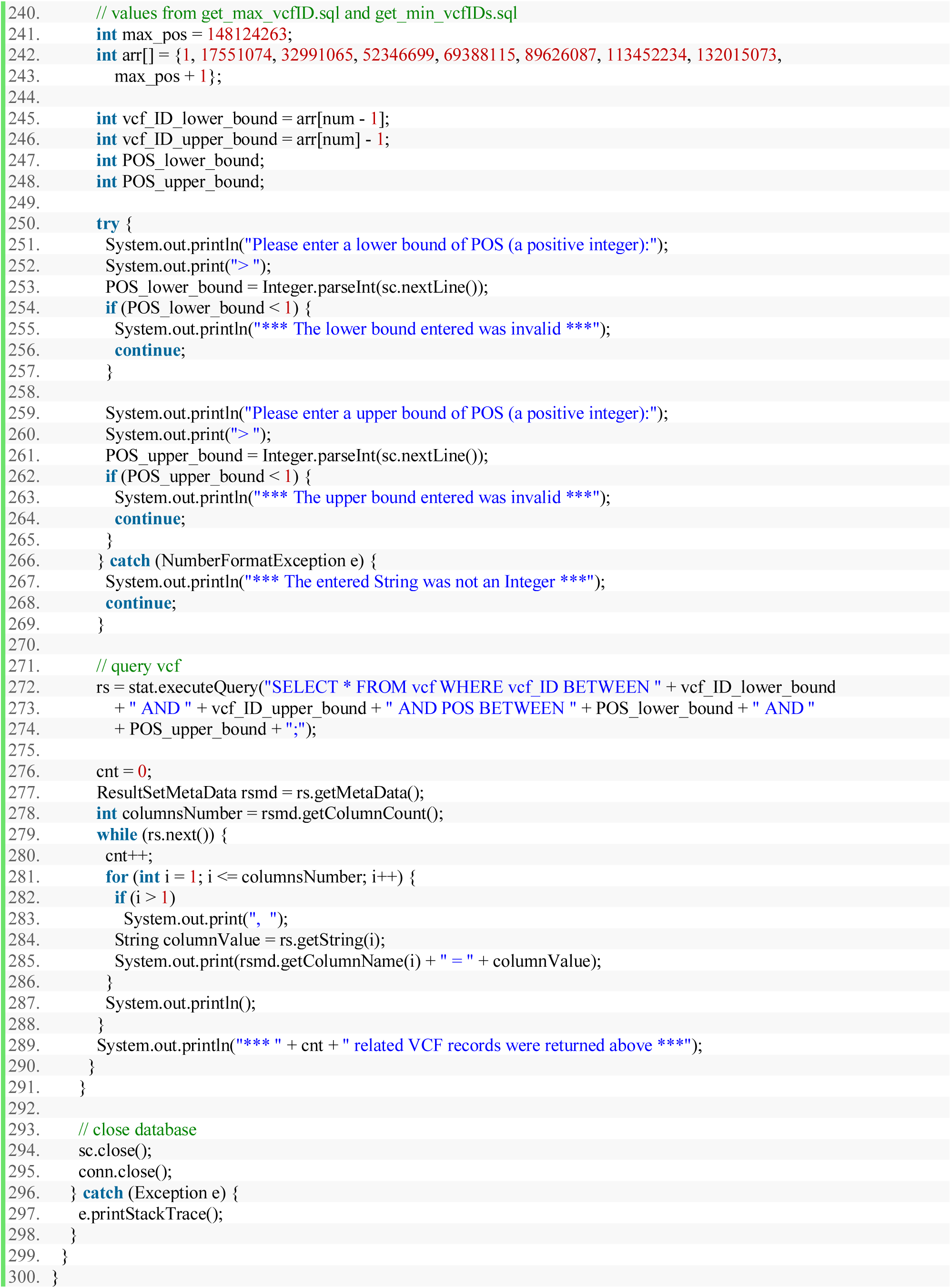

